# A tissue time machine to forward-predict single-timepoint data applied to tuberculosis granulomas

**DOI:** 10.64898/2026.10.09.758083

**Authors:** Pariksheet Nanda, Erin F. McCaffrey, Christian T. Michael, Alea C. Delmastro, Gabrielle Michael, Joshua T. Mattila, JoAnne L. Flynn, R. Michael Angelo, Denise Kirschner

## Abstract

Recent advances have improved understanding of infection; however many detailed datasets are based on single timepoints post-infection and cannot generate longitudinal insights needed to predict infection outcomes. For example, pulmonary infection with *Mycobacterium tuberculosis* (Mtb) initiates a complex system of immune and bacterial interactions that leads to granuloma formation, yet single snapshots in time need elaboration long-term. We develop a new approach, the *tissue time machine*, applied to predict outcomes of non-human primate granulomas from initiation to endpoint using single timepoint tissue datasets including high-dimensional spatial features. This involves identifying granuloma *digital partners*, simulations most closely resembling real granulomas from a virtual library of over 50,000. By analyzing digital partners, we predict infection outcomes of 22/28 granulomas, and that Mtb-specific, IFN-γ-producing T cell recruitment rates influence uncertainty for the 6/28. This method is applicable to any system that allows for mechanistic predictions over time using heterogeneous, incomplete, and high-dimensional datasets.

## Introduction

Recent technologic advances like single-cell RNA sequencing and spatial transcriptomics have accelerated the development of therapeutics by providing a cornucopia of high-dimensional, often spatial datasets^1,2^. While exciting statistical and machine-learning-based tools are emerging to parse these datasets, most data are derived from single-time-point samples, and a need remains to predict datasets forward- or backward-in time. Here, we introduce the *tissue time machine*, a generalizable methodology to use mechanistic computational models to forward- or backward-predict states of a biological object (here, a tissue) in systems where high-dimensional single time-point data are available, biological mechanisms are known, yet system outcomes are heterogeneous and longitudinal data are sparse. We apply this method to study pulmonary tuberculosis (TB) using recent high-dimensional spatial data^3–6^.

TB remains the world’s leading cause of death from a single infectious pathogen; there is no effective vaccine^7^. Following aerosolized exposure to *Mycobacterium tuberculosis* (Mtb), the immune response drives formation of lung granulomas—distinct, spatially heterogeneous structures comprising immune cells that combat Mtb yet serve as sites of long-term Mtb persistence^8–10^. The ability for granulomas to control infection determines whether a host experiences clinically latent infection or active disease (symptomatic, infectious).

Granuloma spatial organization has recently begun to be elaborated as a significant contributor to anti-Mtb immunity^11,12^. Our recent work has shown that granuloma-level immune control correlates with cellular-scale spatial features and networks of interactions between cell types^13^. To further elucidate granuloma infection trajectories, we analyze a unique spatial proteomic dataset from non-human primate (NHP) granulomas harvested between 9–12 weeks post-infection^13^ by creating their *digital partners*—i.e., the set of mechanistic computational simulations that most closely match a specific real-world counterpart^14^.

Our approach identifies several spatial features of granulomas predictive of bacterial burden (CFU), including how these predictors shift over time. As expected, spatial features predictive of later-stage CFU are distinct between early-stage (1–5 weeks post-infection) and mid and later stage infections (5-14, and > 14 weeks post-infection). Additionally, we find that a cluster of spatial features are predictive only during a transient window (weeks 5–14 post-infection) where early granuloma formation dynamics transition to longer-term maintenance dynamics. We identify mechanisms driving CFU levels including the distribution of IFN-γ-producing T cells throughout tissue. Together, these results establish a framework that uses spatial datasets from a single-time-point granuloma image to reconstruct prior infection dynamics, predict future bacterial trajectories, and identify mechanisms that tilt the balance of granuloma success or failure.

## Results

Attributing spatial features of lung granulomas to Mtb infection severity has been difficult and has limited the use of spatial datasets. Single-time-point tissue samples of Mtb-infected granulomas collected from NHPs have not been used to predict longer-term granuloma outcomes. To do this, we synthesize spatial datasets with our computational model *GranSim* (Extended Data 1 and Methods).

### An experimental library of lung granuloma tissue samples

Our first goal is to assemble an *experimental granuloma library* of non-sterile granulomas representative of typical TB pathology for outcome prediction. For this, we select published NHP granuloma datasets collected at necropsy 9–12 weeks post-infection (see Methods) and imaged using multiplexed ion beam imaging by time-of-flight (MIBI-TOF, see Methods)^13,15^ (Fig. 1). We quantify immune cell counts of major lineages from MIBI-TOF images and separately quantify CFU for each sample in the experimental library^13^.

**Fig. 1.**
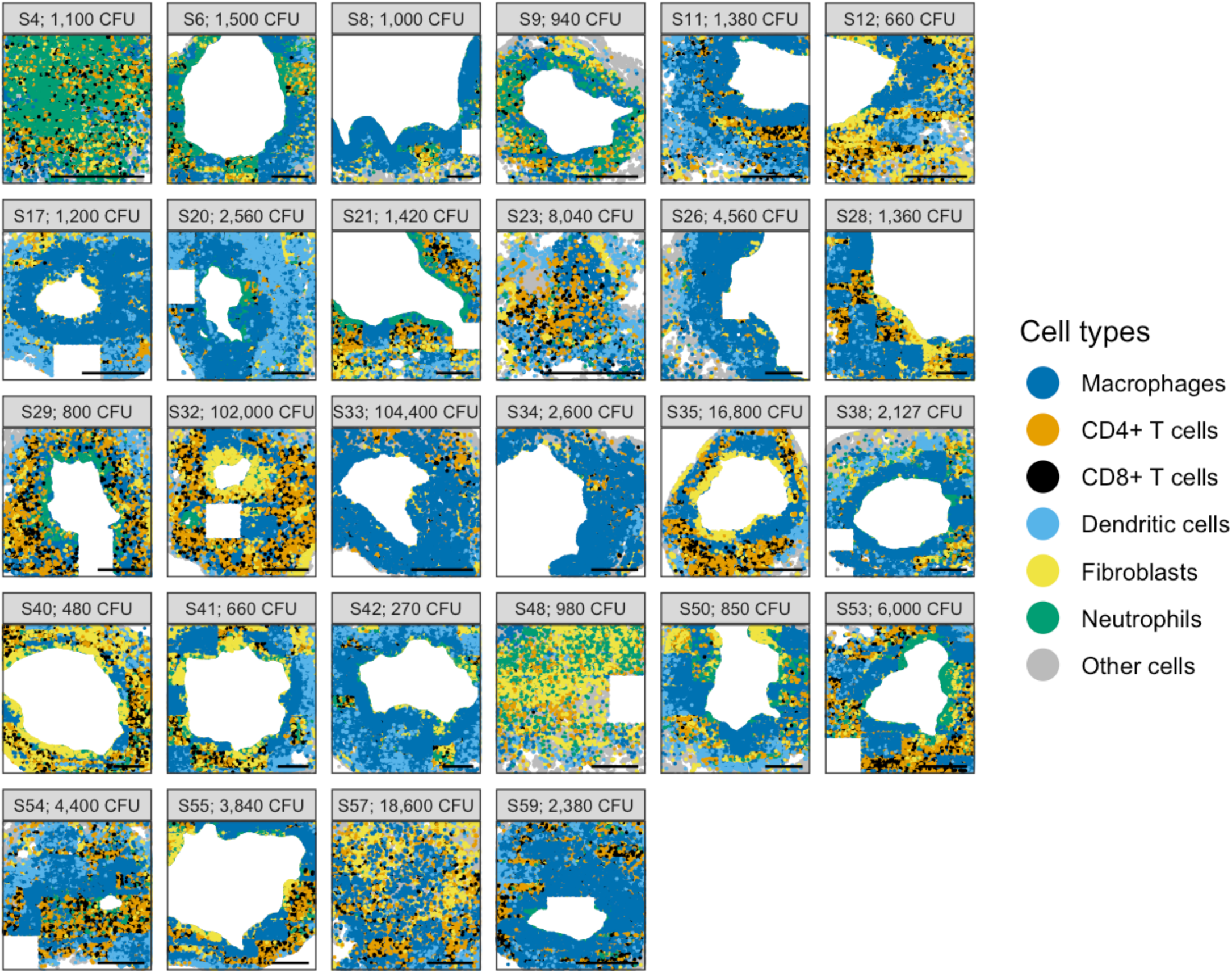
MIBI-TOF-generated experimental lung granuloma library. We show MIBI-TOF images and CFU for each of the NHP granulomas in our experimental library. For comparison, we group MIBI-TOF cells into categories comparable to *GranSim* agents. No samples contain single-cell data within necrotic regions. Some samples have missing tiles that were not captured during the imaging process (e.g., bottom right of sample 8), but these represent a small fraction of each image, and we do not expect them to affect our results. Scale bars at the bottom-right of each image are 500 μm; the imaged tissue area and apparent scale bar size vary across samples because the number of constituent non-overlapping tiles differs to accommodate each tissue section. We consider granulomas that are non-sterile and also have at most 1 necrotic core.

### Simulated virtual library of lung granulomas matches curated spatio-temporal datasets

Our next goal is to generate a *virtual granuloma library* that encompasses the breadth of plausible granuloma outcomes using our *GranSim* computational model of TB^16–18^. We ensure that this library is densely sampled across parameter space such that any granuloma in the experimental library will have many virtual granulomas with similar spatial features (Fig. 2). We chose 22 weeks as an appropriate later time point for a forward-prediction, doubling the average 11-week post-infection necropsy time for most samples. We simulate our virtual library as a set of 60,000 granulomas (20,000 samples × 3 stochastic replicates) over a 22-week timeframe; each granuloma represents interactions of multiple cell types comparable to data (Suppl. Fig. 1). We validate our simulations by matching both immune cell counts and CFU over time with data from 545 granulomas taken serially from previous NHP studies^19–22^ (Fig. 2A and Methods). Fig. 2B shows representative simulated granuloma snapshots at 22 weeks post-infection illustrating sterilization, bacterial control, and uncontrolled growth (Suppl. Fig. 2–Suppl. Fig. 4 for additional representative simulations).

**Fig. 2.**
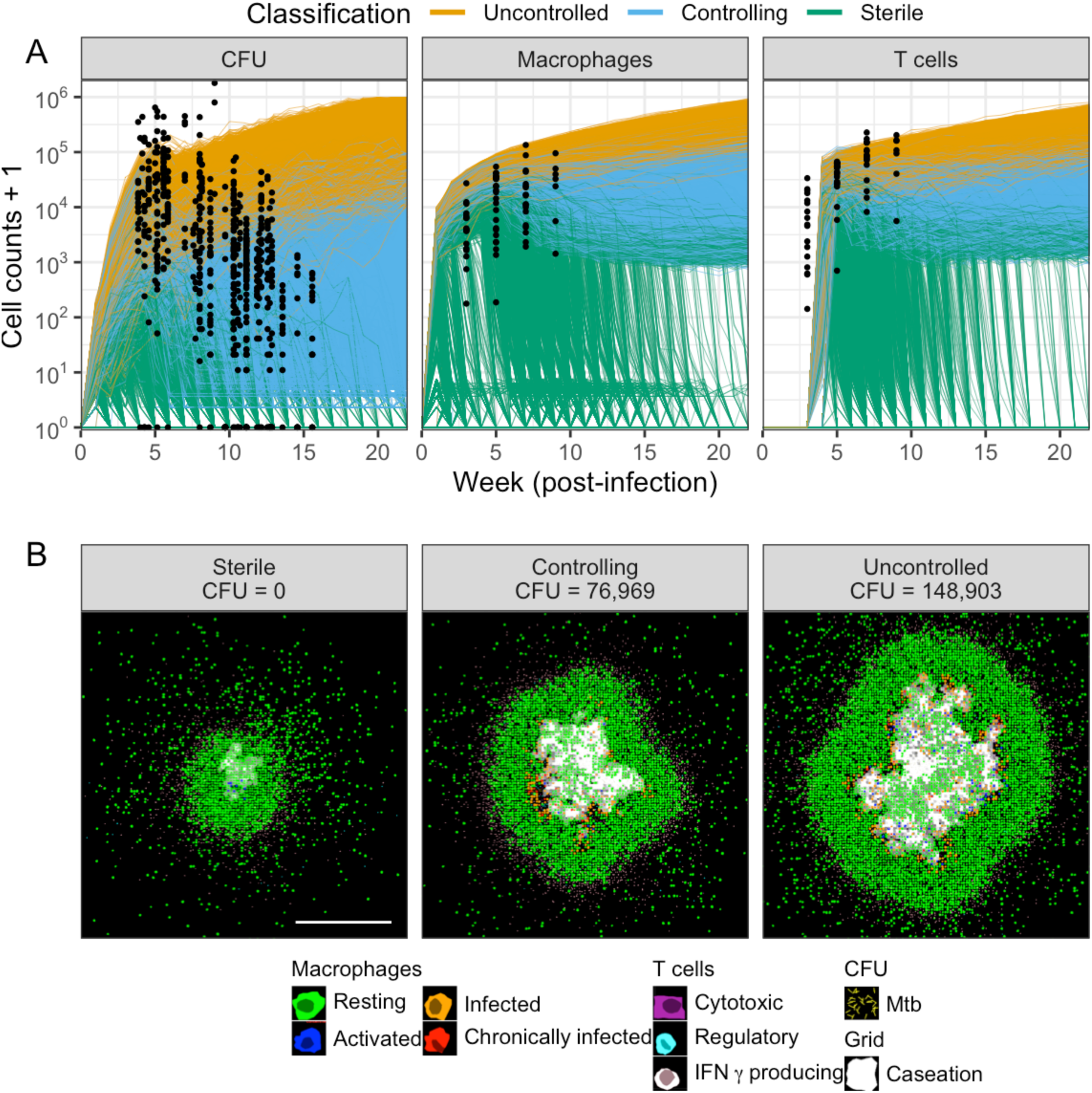
**Virtual granuloma library compared to historical non-human primate datasets**. We simulated 60,000 *GranSim* virtual granulomas (20,000 samples × 3 replicates) from an inoculum of 1 bacterium until 22 weeks post-infection. (A) Of the 53,514 simulations with CFU < 10^6^. Individual granuloma CFU trajectories are colored with individual granulomas assigned to one of 3 groups: sterile (green), controlled (blue), or high/uncontrolled CFU (orange). Sterile granulomas have 0 CFU at week 22 and controlled granulomas have fewer than 10,000 CFU at week 22 or less than a two-fold increase in CFU between weeks 15 and 22 (see Methods). Black dots represent 545 granuloma-scale datasets derived from NHP necropsy samples, including flow cytometry (macrophages and T cells) and growth assay (total CFU), collected weekly. Note that several granulomas are obtained from each non-human primate at the time of necropsy. To show sterilization, counts are offset by 1 before transforming to log scale. (B) Simulations at 22 weeks showing granulomas of each type: sterile, controlled, and high/uncontrolled CFU (read left to right). Additional granuloma images of each type are in **Error! Reference source not found.**–**Error! Reference source not found.**. White scale bar at bottom-right of the sterile granuloma image is 500 μm; the same scale applies to all images.

To characterize the dynamics of Mtb-infected granulomas, we classify the infection state of each virtual granuloma as sterile (0 CFU at 22 weeks post-infection), controlling (<10,000 CFU at 22 weeks, or less than a two-fold CFU increase from week 15 to week 22), or having uncontrolled growth (otherwise; see Methods) (Fig. 2A-B).

In NHPs, ∼50% of granulomas sterilize whether the NHP exhibits latent or active infection^23^. In our virtual library, 52% of our granulomas naturally sterilize, 25% control infection, and 23% experience uncontrolled infection for a total of n=25,503 non-sterilizing virtual granulomas. These values are similar to NHPs, as ∼45% of all infected NHPs progress to active disease^23^, which results from at least one single, uncontrolled granuloma^24^.

### Clusters of cross-sectional spatial features predict granuloma formation-to-maintenance transition

The next step in our pipeline is to generate a set of spatial features predictive of infection outcomes and measurable in both libraries. To compute spatial features, we use CellProfiler, an open-source software for measuring spatial features from cell images^25^. Our pipeline computes a total of 2,240 candidate spatial features for both *GranSim* granulomas and analogous measurements for MIBI-TOF images (see Suppl. Fig. 1 and Suppl. Table 1). At the cellular-scale, features describe the position of cells relative to their neighbors and cell-colocalized cytokine levels (Extended Data 2). At the tissue-scale, features include aggregate arrangements of cells, cytokine distributions, and granuloma architecture.

To explore how granuloma organization associates with bacterial burden, we use our virtual library to temporally identify the features most predictive of granuloma log-CFU. We compute each spatial feature for each MIBI-TOF sample; we repeat the same for all virtual granulomas for weeks 1-22 post-infection. Using linear regression, we rank spatial features by their predictive power for virtual granuloma log-CFU at 22 weeks and select the top 20. We repeat this ranking independently for each week, allowing the most predictive features of week-22 CFU, out of the initial 2,240 features, to change over time. We use a clustering analysis to further narrow our focus to spatial features that are included in the top 20 predictive of later-stage CFU for at least four of the 22 simulated weeks contiguously (Methods). In this study, we consider only non-sterilizing granulomas since sterilizing virtual granulomas would more appropriately be compared with experimental “time-to-sterilization” datasets, which are not available. Moreover, by isolating non-sterile granulomas, we focus on dynamics that characterize late-stage CFU during latent and progressive infection.

Examining top predictive spatial features from non-sterilizing granulomas in our virtual library, we find that they group into three distinct temporal clusters (Fig. 3). The first cluster includes features that are predictive during early granuloma formation, spanning the first five weeks of infection (Fig. 3, blue cluster). This cluster is characterized by macrophage-adjacent TNFα (produced in our model by macrophages and T cells), the size of the macrophage population perimeter in a granuloma, and the mean macrophage-to-macrophage distance. Over subsequent weeks, another cluster emerges that we term the “granuloma maintenance cluster” (Fig. 3, green cluster). This cluster is dominated by IFN-γ-mediated macrophage activation near the outside of the granuloma and the population diameters of macrophages and T cell subtypes. We also observe a final cluster that we refer to as the “transient cluster” (Fig. 3, orange cluster). This cluster is characterized by macrophages activated by IFN-γ near the inside of the granuloma and coincides with T-cell recruitment from lung-draining lymph nodes to the granuloma^26–28^. Therefore, in addition to the expected granuloma formation and maintenance clusters, our analysis reveals a transient phase (weeks 5–14 post-infection) whose spatial features may be critical for determining later-stage granuloma outcomes. Coincidentally, all samples in our experimental library were collected within the window of 9–12 weeks, suggesting that they were collected during the transient phase.

**Fig. 3:**
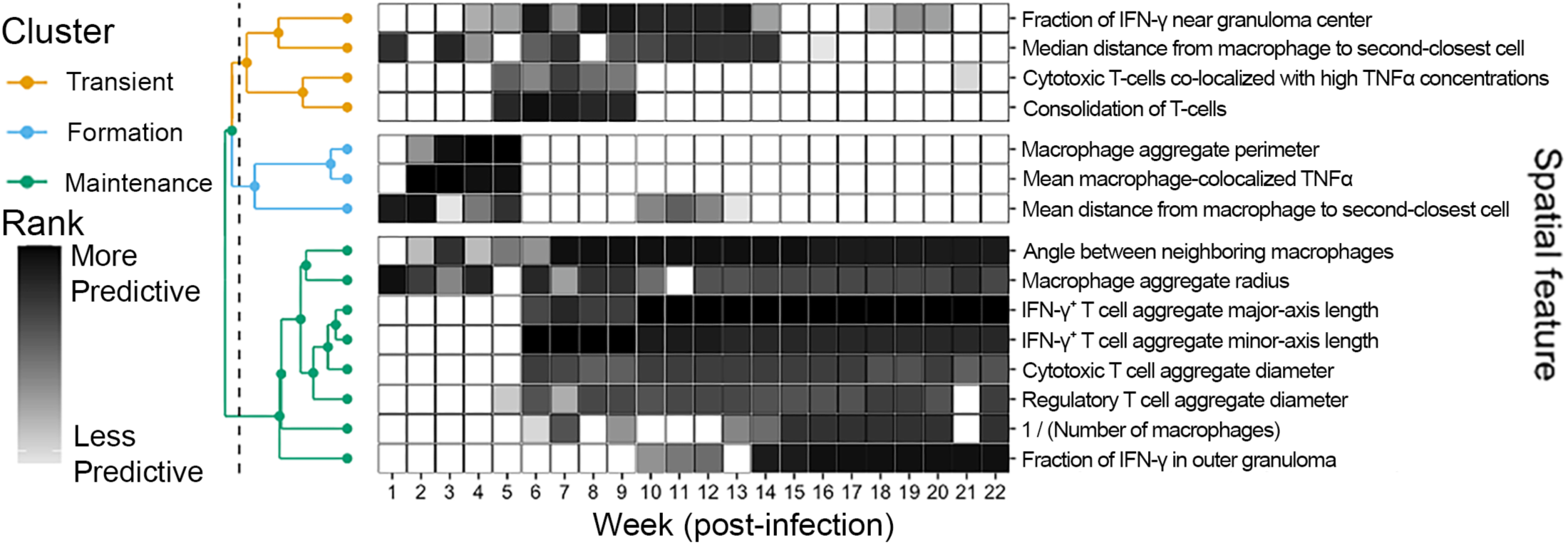
**Top predictive granuloma spatial features form temporally organized clusters in non-sterilizing granulomas**. We identify three clusters: features associated with granuloma formation (up to five weeks, blue dendrogram), features associated with a transient phase when T cells are first recruited from lung-draining lymph nodes (5–14 weeks, orange dendrogram), and features associated with granuloma maintenance (six weeks onwards, green dendrogram). We do not show spatial features that appear less than four times contiguously during the 22 weeks here. We cluster predictive spatial features using the Canberra distance metric (see Methods). We provide mappings of CellProfiler feature names to our descriptive feature names in **Error! Reference source not found.**.

### Both tissue-scale and cellular-scale spatial features are necessary to predict bacterial burden

To determine the most important features for predicting bacterial burden at a final timepoint (and thus for selecting digital partners) we use linear regression to assess how well the top 20 spatial features predict CFU in the virtual library. Using Bayesian variable selection (see Methods), we find that a small collection of 12 features is sufficient to recapitulate the heterogeneous distribution of bacterial burden among non-sterilizing virtual granulomas at 22 weeks post-infection (Extended Data 3; Suppl. Fig. 8–Suppl. Fig. 9). These 12 features capture the dominant mechanisms driving CFU at 22 weeks (Extended Data 4 and Suppl. Fig. 10). We find that granuloma size and shape have a complex relationship to bacterial burden, and we discuss these in detail in the Supplemental Material.

The predictive power of IFN-γ has opposing effects on CFU when it is spatially distributed toward granuloma center versus edge, reflecting the importance of T-cell penetration into the myeloid core. Higher IFN-γ at the granuloma center (Extended Data 4 inner fraction) correlates with lower CFU, indicating that the deeper cells expressing IFN-γ penetrate into granulomas, the lower CFU is. Although IFN-γ is not typically observed near the centers of experimental granulomas, this suggests that CFU would be low in granulomas if T-cells could penetrate deeper within, and infection might possibly resolve even before granulomas become large enough to observe. Previous studies suggested that T cells have difficulty penetrating into the core of a granuloma once a macrophage-rich granuloma has formed, prior to T cell recruitment^26^. Conversely, IFN-γ distributed toward the granuloma edge correlates with higher CFU, indicating that when T cells remain confined to the lymphocytic cuff they fail to activate macrophages near bacteria located closer and within the granuloma center. IFN-γ^+^ T cells are likely recruited and activated earlier via signals like TNFα, evident by the appearance of TNFα-T-cell co-localization in the transient cluster (Fig. 3). This underscores how macrophages are a double-edged sword in TB; activated macrophages are required to kill Mtb, but resting macrophages are the main cellular host for Mtb and can sequester Mtb from bactericidal, T-cell-activated macrophages.

We identify only partial overlap between the two small feature sets predictive of week-22 CFU: one using spatial immune features from week 11 (as representative of the 5–14-week transient window) (Fig. 3) and one using week-22 features identified via Bayesian regression (Extended Data 4). Both feature sets originate from the same 2,240 candidate features identified using CellProfiler (Extended Data 2) and share negative correlates of CFU (e.g., increased macrophage activation rates within a granuloma, and less densely distributed macrophages). However, not all features are shared, notably TNFα proximity to cytotoxic CD8^+^ T cells. This suggests that the necessary mechanisms driving CFU outcomes are time-dependent, with some drivers emerging specifically during the transient phase of granuloma development (weeks 5-14, Fig. 3).

### Novel identification of digital partners of experimental tissue samples

Our next goal is to use spatial features common to both libraries (Extended Data 1) to identify *digital partners* for each experimental granuloma. We digitally partner each of the 28 MIBI-TOF samples (single-timepoint, non-sterile NHP granulomas) with non-sterilizing granulomas from our virtual library. This allows us to make forward- and backward-looking temporal predictions of the outcomes for the individual samples, as if we had been able to observe them longitudinally (namely, a *tissue time machine* predicting both forward and backward in time). To digitally partner simulated granulomas with experimentally derived samples, we apply a two-step process: first identifying candidate digital partners (CDPs) based on log-CFU similarity to the experimental samples, and second ranking the CDPs based on spatial-immunological similarity using the features discussed above (Fig. 4).

**Fig. 4.**
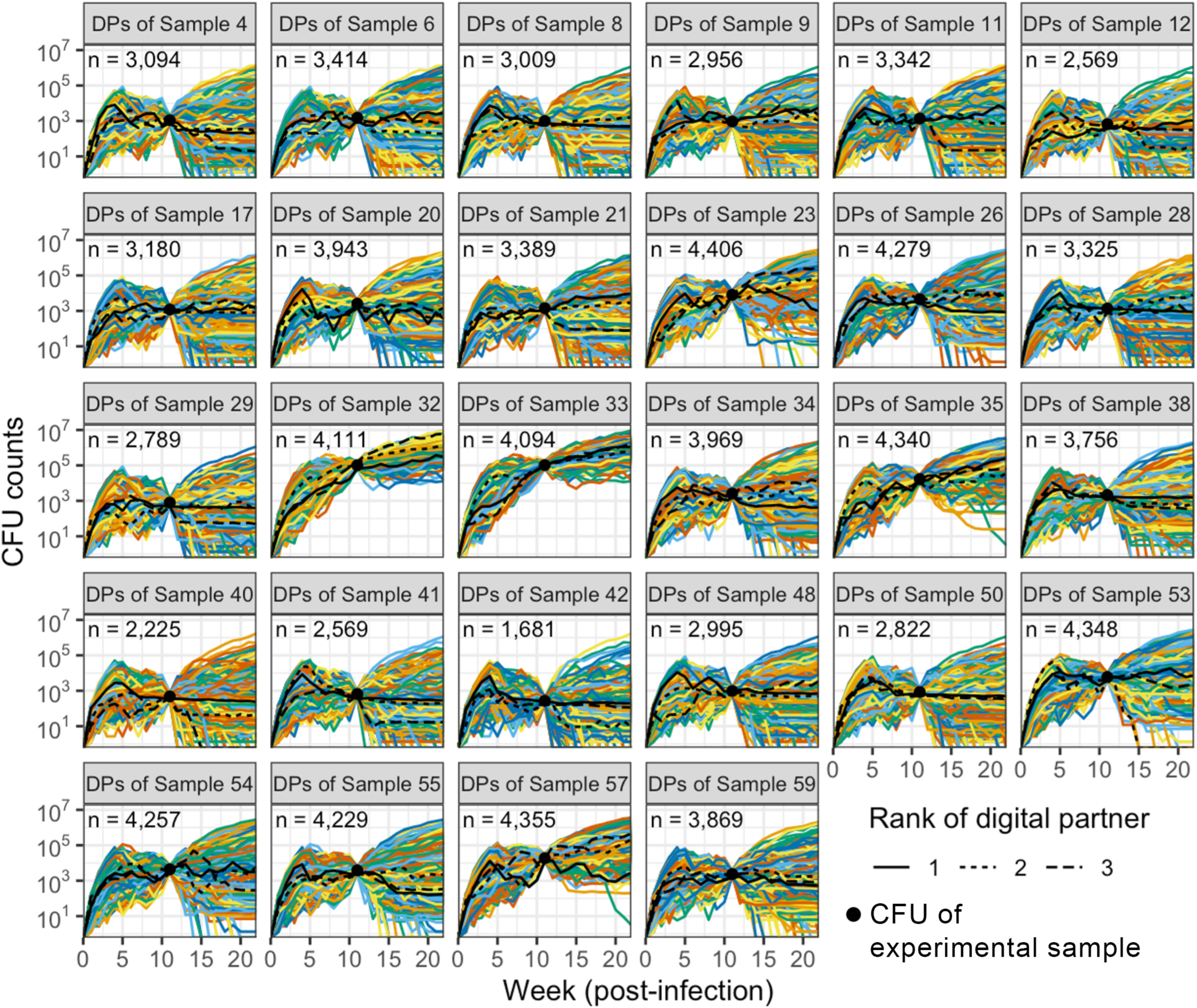
**Digital partners for 28 NHP granulomas**. We match candidate digital partners (CDPs; all curves) from the virtual library to each experimental library sample by comparing the distance between experimental and simulated log-CFU (see Methods). We use optimal transport (OT) to rank the closeness of each MIBI-TOF sample to each CDP (see Methods). We select ranked digital partner simulations by successive elimination (see Methods). We indicate the nearest digital partners to the experimental NHP granulomas with black curves.

To predict forward and backward temporal predictions for each experimental granuloma (had they remained *in vivo*), we use each sample’s top-ranked digital partners. We identify their DPs at week 11 post-infection and examine those partners’ simulated infection outcomes at 22 weeks post-infection. We predict the NHP granuloma outcome would be the same as the simulated outcomes of the top three DPs if they are in consensus, and indeterminate if they are not. Fig. 5 shows three of the 28 representative granulomas and their forward and backward temporal predictions from the top-ranked DPs (all others are shown in Suppl. Fig. 12–Suppl. Fig. 21). Predicted outcomes of all 28 granulomas are in Extended Data 5.

**Fig. 5.**
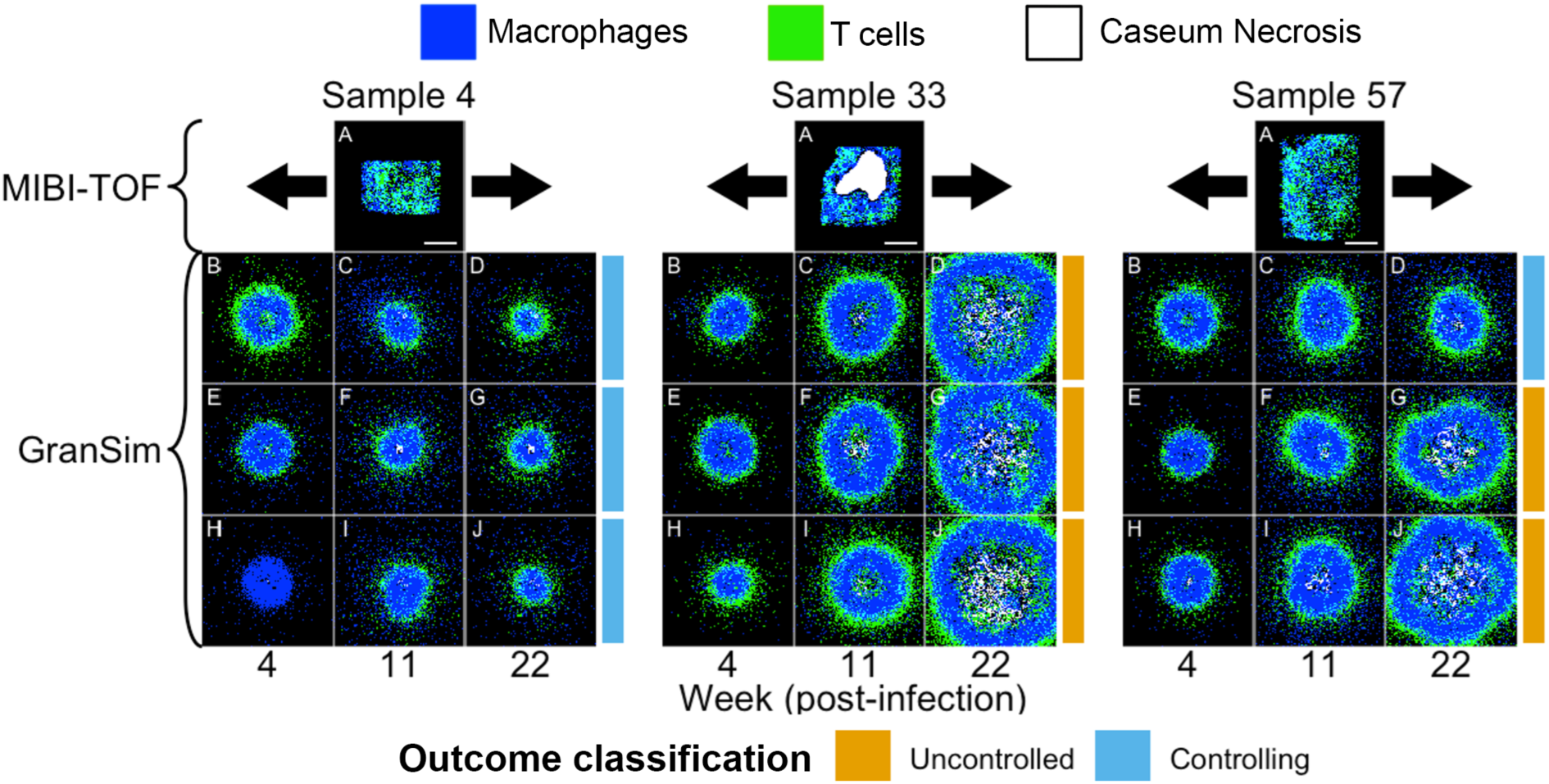
**Tissue time machine simulations for the closest digital partners of three NHP lung granulomas**. We show the three closest digital partners identified by OT and successive elimination for MIBI-TOF samples 4, 33, and 57. These three samples illustrate a range of predicted outcomes: all DPs controlling (sample 4), all DPs uncontrolled (sample 33), and an indeterminate mix of controlling and uncontrolled (sample 57). Consistent with the colors in Fig. 2, we indicate controlling (blue) and uncontrolled (orange) outcomes for each digital partner at 22 weeks. For each MIBI-TOF sample, the corresponding images are as follows: (A) MIBI-TOF sample showing macrophages (blue), T-cells (green), and caseous necrosis (white). The MIBI-TOF have rectangular fields of view composed of multiple 1024 × 1024 non-overlapping tiles. The white scale bar at the bottom-right is 500 μm for all images. (B-D) We show the closest ranked digital partner (rank 1) at weeks 4, 11, and 22. (E-J) We similarly show the second and third closest digital partners (ranks 2 and 3) at weeks 4, 11, and 22. Note that (B-J) are zoomed in to show detail at the same scale as the MIBI-TOF sample (namely 2 mm × 2 mm of the simulations), but the simulated grid size is 6 mm × 6 mm. We present digital partners for all MIBI-TOF-imaged granulomas in **Error! Reference source not found.**–**Error! Reference source not found.** and their corresponding outcomes in **Error! Reference source not found.**.

Ideally, the top digital partners for a given sample should agree with each other in their predicted later-stage outcomes. This happens for 22 of 28 MIBI-TOF samples, where each of the top 3 DPs agrees with the predicted granuloma outcome at 22 weeks (Extended Data 5). In 20 samples, all 3 DPs were controlling, and, in 2 MIBI-TOF samples all 3 DPs were uncontrolled. For the other 6 of 28 granulomas, the DP prediction is indeterminate (Extended Data 5). Nevertheless, the pipeline predicts a consensus outcome for 22 of 28 samples (∼80%). This strongly suggests that additional factors absent from either the simulation or the tissue measurements are needed to further refine the pipeline. Out of all virtual granulomas at 11 weeks post-infection, 66% (21,251 of 32,101) were CDPs for at least one experimental library sample. We explore the differences between these outcomes next. We also surmised that 3 DPs were insufficient to reliably forecast all cases, and for all subsequent analyses, we use the top 50 digital partners for each experimental sample to better capture outcome heterogeneity.

### Mechanisms driving granuloma infection outcomes across tissue and cellular scales

Whereas above we compare spatial features to infection outcomes, here we examine mechanistic drivers underpinning those infection outcomes. We do this by examining the 50 closest-ranked digital partners for each of the 22 experimental granulomas whose top-3 digital partners reached consensus predictions (as described above). This yields 1,100 virtual granulomas that we use to predict the 22-week outcomes of experimental granulomas (collected at 9-12 weeks post-infection). We use the partial rank correlation coefficients (PRCC) method, which uses nonlinear correlations to identify mechanisms driving infection severity (Fig. 6A-G).

**Fig. 6.**
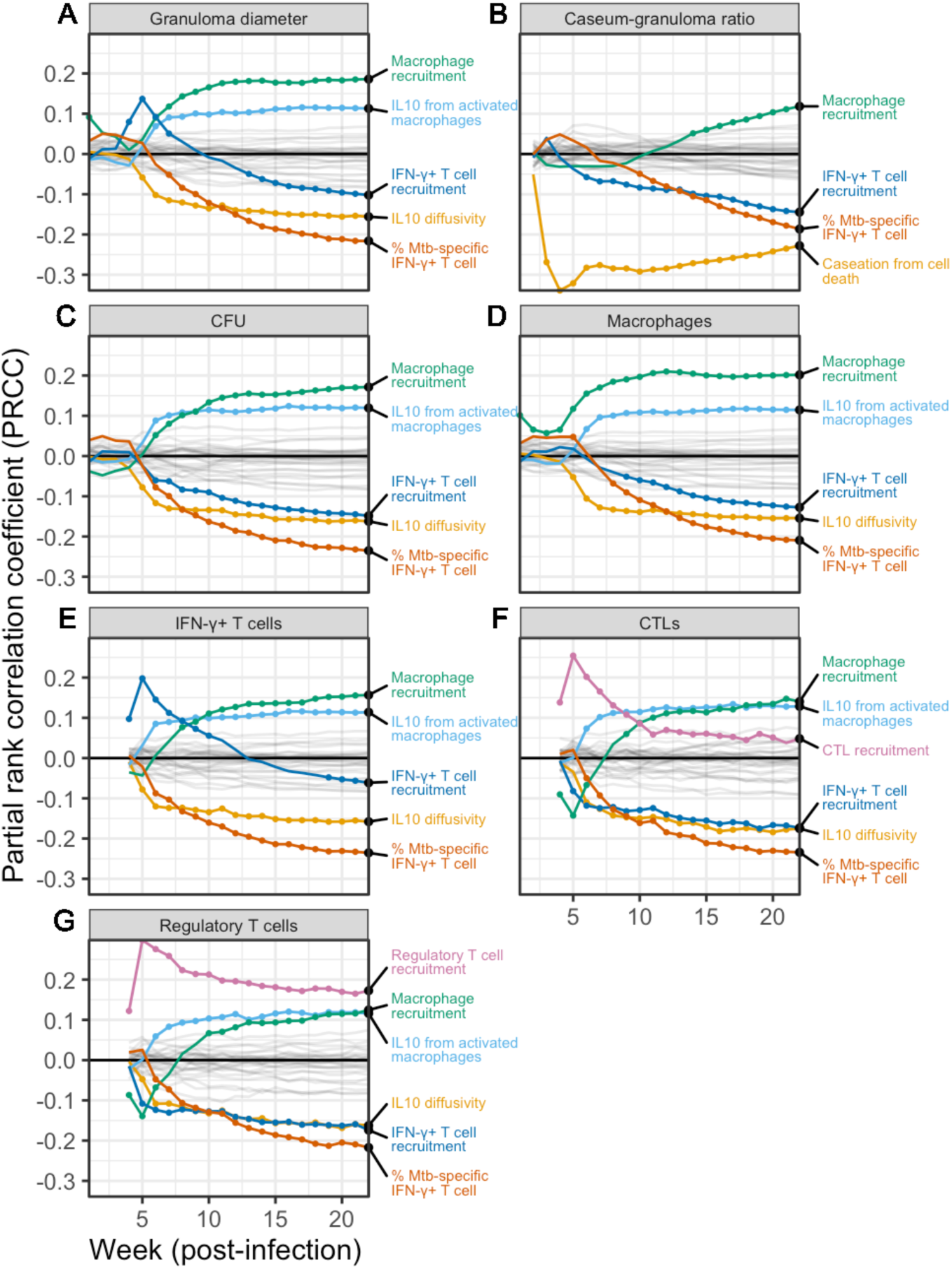
Identification of significant mechanisms driving tissue-scale features, CFU, and immune cell levels during infection. Shown are partial-rank correlation coefficients (PRCCs) derived from uncertainty and sensitivity analysis (U/SA) of the 50 closest ranked digital partners for each of the 22 MIBI-TOF tissue samples with predicted outcomes in consensus (first group in **Error! Reference source not found.**). We vary model parameters and calculate PRCCs driving tissue-scale outputs at each week from 1-22 for all digital partners (50 for each sample, yielding a total of n = 1,100 digital partners). Each panel shows the PRCC values a single simulation outcome: (A) granuloma diameter, (B) caseous-to-cellular-area ratio, (C) total CFU, (D) Macrophage count, (E) IFN-γ^+^ T cell count, (F) CD8^+^ Cytotoxic T lymphocyte count, and (G) regulatory T cell count. We identified influential mechanisms for each outcome using the boxplot whisker outlier test (see Methods) and are shown as colored curves. Timepoints of those impactful mechanisms with p-value < 0.05, corrected by the Benjamini–Hochberg false discovery rate (BH-FDR), are shown as colored dots along their respective curves. Non-influential reference parameters are shown as semi-transparent gray curves and not labeled (a total of 52 mechanisms characterized by 63 model parameters were considered in this analysis).

We explore parameters influencing two tissue-scale *GranSim* outputs: granuloma diameter and caseum-to-cellular area ratio. Fig. 6A shows that IFN-γ^+^ T cells have a nonlinear impact on granuloma diameter; early recruitment stimulates an inflammatory response leading to a larger, immune-dense granulomas, while improved T cell recruitment later during infection supports bacterial control and smaller granuloma sizes. The correlation coefficients change sign between weeks 7 and 15, aligning with the transient phase identified above (Fig. 3). Macrophage activation and IL-10 secretion also influence granuloma outcomes; both the number of macrophages and the rate of activated macrophage production of IL-10 is associated with increased regulatory T cell levels and increased granuloma diameters. Our finding that the role of IL-10 changes over time is consistent with previous studies of primary granulomas in NHPs^29–31^. Most of the same mechanisms drive cellular-scale outcomes (Fig. 6C–G); we discuss the impact of macrophages and IL-10 on granuloma outcomes along with extended analysis of cell-scale outcomes in more depth in the supplementary material.

We repeat this analysis comparing results with the 6 of 28 experimental granulomas whose digital partner forecasts were indeterminate (Extended Data 6). Specific to the indeterminate granulomas, regulatory T cell recruitment positively correlates with granuloma diameter, caseum-to-cellular area ratio, CFU, macrophages, and IFN-γ^+^ T cells. The frequency of Mtb specificity among cytotoxic CD8^+^ T cells negatively correlates with granuloma diameter, CFU, macrophages, IFN-γ^+^ T cells, and regulatory T cells. Finally, CFU correlates with secretion rates of IL-10 from infected macrophages. This suggests that the indeterminate virtual granulomas are at an immunological tipping point that is influenced by suppressive drivers. To predict outcomes of such indeterminate granulomas may require using datasets that track these mechanisms as part of digital partner selection.

## Discussion

Tuberculosis granuloma trajectories are challenging to predict, both because of their spatial complexity and current limitations faced by available experimental methods. New spatially-detailed imaging techniques like MIBI-TOF^13,15^ capture granuloma organization that is associated with granuloma outcome^13,26,32–34^, although such data are only available at a single time point per granuloma. To predict longitudinal outcomes of real lung granulomas, we present the *tissue time machine*: a new approach that combines detailed single-time-point tissue data with our spatio-temporal computational model. Here, we apply our tissue time machine approach to predict both early and late infection states of 28 NHP granulomas by synthesizing mechanisms known to drive infection outcomes with NHP granuloma datasets.

While the digital partners of each NHP granuloma match multiple measurable features of the sample at 11 weeks post-infection, the tissue states both temporally prior to and later than 11 weeks can differ substantially. Thus, there are many infection trajectories that can converge to the tissue state we observe in individual MIBI-TOF images, and some feature-equivalent images may have diverging outcomes. However, digital partners frequently predicted the same outcome states, suggesting that the decisive point of inflection for granuloma fate occurs during granuloma formation (up to five weeks post-infection). Our sensitivity analysis indicates that prediction accuracy will be improved by collecting NHP samples for spatial analytes of IL-10 and IFN-γ-producing Mtb-specific T cells, as they are not currently measured in the NHP MIBI-TOF dataset.

For samples whose outcomes could not be consistently predicted, we find that regulatory T cells drive the heterogeneity in our forecasts. Regulatory T cells are known to have a complex impact on granuloma outcomes, both by possibly impairing T cell activation^32^ and cytotoxic T cells activity, but also in resolving inflammation. Cytotoxic T cells combined with low levels of regulatory T cell activity is a marker of active TB^35^, despite cytotoxic T cells typically playing a protective role^36^. Additionally, recent spatial biology datasets show higher regulatory T cell infiltration in the granuloma myeloid core that has the potential to impair T cell activation^32^.

Our analysis also uncovers a cluster of transient features in between these stages, characterized by cytotoxic T cells, IFN-γ, TNFα, and a consolidation of the T cell population. Our analyses suggest that TNFα helps accelerate a robust adaptive immune response (as it appears in the transient cluster), laying the groundwork for a protective response associated with co-localized Mtb-specific T cells with macrophages deep within granulomas. We further validate the predictive power of the top-ranked spatial features by reproducing the distribution of bacterial burden using a minimal set of 12 features.

Mtb-specific IFN-γ-producing T cells prominently arises in our analyses. As tetramer studies are currently not possible on human granulomas and rare even in NHPs, how the frequency of Mtb-specific T cells impacts granuloma outcome remains unknown^37^. However, our analysis suggests that the timing and magnitude of these cells help orchestrate immune control, and recent data show that long-lasting protection against TB is CD4^+^ T cell-dependent^31^.

All models are incomplete, having both technical and resource limitations. We base our predictions on the top three digital partners, and more work is required for determination of an appropriate number; selection of this number may indeed characterize predictability of a biological system, its corresponding mechanistic model, or datasets involved, e.g., moving to fifty CDPs for our sensitivity analysis added significant statistical power to our predictions. In the future, we may employ more advanced methods such as convolutional neural network analysis to uncover nonlinear relationships among spatial features, rather than linear regressions. Lastly, we may compare our 2-dimensional (2-D) *GranSim* model results with our 3-D *GranSim* model outputs^38^ as the 2-D model is more sensitive to artifacts related to cell-cell co-localization^38^. While computational limitations currently makes the 3-D model infeasible for this application, we minimize the impact of this simplification by scaling our 2-D outputs into 3-D as we have previously described^39^.

Here, we have used the tissue time machine method in the context of Mtb infection, but we can apply this approach to study any system where longitudinal data are scarce, single-time-point data are available, and a mechanistic model can be developed. Much like medical digital twins^40^, the digital partner/tissue-time-machine approach is not restricted to MIBI-TOF data; one may use any high-dimensional and/or multimodal datasets taken from an individual subject for forward prediction. The process to extend the tissue time machine approach forward or backward to predict a state of a different system is straightforward (e.g., a tumor or an organ). First, generate a large, heterogeneous library of virtual subjects with sufficient mechanistic detail to produce emergent features measurable in the subject. Second, establish a procedure to use all available datasets to isolate the subject’s most-similar digital partner set. Third, examine the digital partner outcomes of the subject. If the subject’s digital partners are in consensus, then the prediction is robust and the digital partners and virtual library can be studied via sensitivity analysis to identify key driving features. If the subject’s digital partners are not in consensus, then analyzing the digital partners may identify which mechanisms or data are missing for subject-specific predictions. The outputs that can be studied are any feature that can be credibly predicted by the model framework.

## Extended Data and supplementary material

PDF documents containing Extended Data 1–7 and the Supplementary Material are available on bioRxiv. Extended Data citations refer to the Reference list of the main text.

## Acknowledgments

We thank Paul Wolberg for his support with computer programming. This research was supported by NIH Grants R01 AI50684 (DEK) and by funding from the Wellcome Leap ΔTissue Program awarded to DEK, MA, JM and JLF. ACD was supported by NIH Grant F31 AI191425. This work used Anvil at the Purdue University Rosen Center for Advanced Computing and Expanse at the San Diego Supercomputer Center through allocation MCB140228 from the Advanced Cyberinfrastructure Coordination Ecosystem: Services & Support (ACCESS) program, which is supported by National Science Foundation grants #2138259, #2138286, #2138307, #2137603, and #2138296. This research was supported in part through computational resources and services provided by Advanced Research Computing (ARC), a division of Information and Technology Services (ITS) at the University of Michigan, Ann Arbor. This research was supported in part through computational resources and services provided by the University of Michigan Medical School, which is supported by NIH grant NIH S10 OD026825. The authors would like to thank the Consulting for Statistics, Computing and Analytics Research (CSCAR) at the University of Michigan for providing valuable insights on developing the data analysis pipeline.

## Author contributions

**All authors**: Writing - reviewing and editing. **Denise Kirschner**: Conceptualization (lead); Funding acquisition; Project administration; Resources; Supervision; Writing - original draft (supporting). **JoAnne L. Flynn**: Conceptualization (supporting); Funding acquisition; Resources. **Michael Angelo**: Funding acquisition; Project administration; Resources; Supervision. **Joshua T. Mattila**: Conceptualization (supporting); Funding acquisition. **Gabrielle Michael**: Investigation; Software; Visualization. **Christian T. Michael**: Methodology; Validation; Writing - original draft (supporting). **Alea C. Delmastro**: Data curation; Software. **Erin F. McCaffrey**: Conceptualization (supporting); Data curation; Investigation; Software. **Pariksheet Nanda**: Data curation; Investigation; Methodology; Software; Supervision; Validation; Visualization; Writing - original draft (lead).

## Methods

### Tissue Time Machine Method Overview

TB is a disease for which little longitudinal data are measurable, thus we seek to extend detailed single-time-point datasets forward and backward in time. The dataset we are starting from (i.e., the experimental library) is a spatial proteomic dataset from NHP granulomas harvested between 9–12 weeks post-infection^13^ and imaged using multiplexed ion beam imaging by time-of-flight (MIBI-TOF)^13,15^, a mass spectrometry-based multiplexed imaging platform. Our experimental library excluded non-sterile granulomas because we do not know when, during the course of infection, those granulomas sterilized. However, an advantage of our pipeline is that the types of granulomas we analyzed can be tailored or extended as data become available. We considered granulomas with at most 1 central necrotic core.

We previously introduced the concept of digital partners as the set of mechanistic computational simulations that most closely match a given real-world counterpart, such as a given experimental observation, clinical patient, or biological sample^14^. Similar to digital twins^40^, the goal of digital partners are to predict specific outcomes. However, a critical component of digital twins are a continuous stream of information from the real-world counterpart into the digital twin for dynamic updating^40–43^, which is not possible in systems where longitudinal data or repeated sampling are not available. Moreover, in complex and/or heterogeneous systems where many mechanisms are known but insufficient data are available to precisely calibrate to patient specific data, models may not be identifiable, especially when one attempts to calibrate that model to a specific patient. Thus, the digital partner framework is to simulate a large ensemble of virtual counterparts capturing a vast set of plausible outcomes and, from that population, isolate a set of candidate digital partners (CDPs), simulations that are consistent with observable data. By observing the late-stage behavior of those digital partners, we forward-predict the outcomes of the subject in question, had it remained *in situ*.

To create our virtual granuloma library, we employ our established mechanistic computational model of TB granulomas, *GranSim*. *GranSim* is a stochastic, agent-based, computational platform that simulates spatio-temporal dynamics of TB lung granulomas and has been continuously curated with both human and NHP datasets over 20 years^16–18^. *GranSim* captures immune cells in various states (primed, effector, and regulatory T cells; resting, active, and infected macrophages), chemokines (CCL2, CCL5, CXCL9), cytokines (TNFα, IFN-γ, IL-10), individual Mtb in different metabolic states, and a caseous necrotic core. Using these components, *GranSim* yields emergent virtual granulomas that recapitulate spatial dynamics of lung granulomas over time.

We use *GranSim* to simulate a virtual library of over 50,000 TB granulomas that span the spectrum of plausible human and NHP granuloma outcomes. We identify multiple digital partners for each granuloma by quantifying similarity across multiple weighted spatial measurements, discussed later in the Methods section. We then use these partners to trace mechanisms, forecast outcomes, and uncover underlying drivers of observable spatial features^14,44^. Granuloma-scale simulation is relevant to host fate because a single uncontrolled granuloma is sufficient to drive progression to active disease^20,23^.

Pairing high-dimensional, multiplexed tissue images with simulations has been limited to models that must be initialized by spatial immunolabels^45,46^. We overcome this methodological limitation using optimal transport (OT), which has been shown to effectively couple heterogeneous datasets, including dynamical models with single-cell transcriptomic data^47–50^. To our knowledge, this is the first instance of applying OT to predict TB outcomes. Our *tissue time machine* pipeline uses OT to couple spatial features from both the tissue samples and simulation images, eliminating the strict requirement that all the model parameters have directly corresponding spatial immunolabels. This allows for the use of sophisticated spatio-temporal computational models that extend what is directly observable in tissue.

### Selection of NHP granulomas

We required each granuloma in the library to have at most one, centralized, caseous necrotic core both to match the typical granuloma presentation and to ensure compatibility with the single-core architectures of the granulomas in our virtual library (Fig. 1). In this analysis, we are only considering non-sterile granulomas. Thus, we omit sterile NHP granulomas because we cannot know the timing of sterilization. Previously when we explored *GranSim* granuloma sterilization we found that control was associated with rapid CD4^+^ T-cell priming^51^ and that the frequency of Mtb specificity directed the granuloma cytokine milieu^26^ (and, by extension, granuloma spatial organization). We also predicted sterilization using machine learning methods trained on synthetic data^33^, though such approaches do not yield interpretable spatial features.

In future work, we will expand our studies to also include sterilized granulomas. More work needs to be done to directly compare *GranSim* sterilization with experimental datasets, as we need a new way to identify digital partners without knowing the time of Mtb sterilization in the animal datasets. The digital partner methodology allows us to create principled, mechanistically-grounded forward-predictions that are otherwise biologically inaccessible and can integrate known biological mechanisms to predict the impact of interventions such as novel TB vaccines.

### Non-human primate lung granulomas acquisition

For all of our analyses, we use published MIBI-TOF datasets generated in a previous study^13^. There, a total of 52 archival tissue samples of granulomas from 16 cynomolgus macaques (*Macaca fascicularis*) were analyzed. These macaques had been infected with low-dose Mtb Erdman (2-13 CFU) by way of bronchoscopic instillation and followed until 9–12 weeks post-infection, followed by necropsy. Granulomas from these macaques included histologically solid granulomas, cellular granulomatous inflammation surrounding central necrosis, fibrotic granulomas, and multifocal granulomas^13^. Formalin-fixed paraffin-embedded tissue sample blocks were stored in the University of Pittsburgh’s tissue repository. From those samples, serial 5–μm sections stained with hematoxylin and eosin were examined to identify Mtb infection. Full details are available in^13^.

### Antibody preparation, tissue staining, and MIBI-TOF imaging

The MIBI-TOF data utilized in this study are part of the dataset generated by McCaffrey & Delmastro et al^13^ and complete methodological details are described there. Briefly, the first step in generation of this dataset was identification of antibodies that cross-react with the cynomolgus macaque epitopes imaged here. To achieve this, we screened human-specific reagents with single-plex chromogenic immunohistochemistry (IHC) in cynomolgus macaque control tissues (lymph node, spleen, lung, and colon) to confirm epitope cross-reactivity and appropriate staining patterns. All antibodies were then conjugated to isotopic metal reporters with the IonPath conjugation kit (IonPath, Menlo Park, USA). All antibodies were lyophilized and stored long-term at −20°C. To optimize the performance of the antibodies for MIBI-TOF, antibodies were reconstituted and combined into a single panel. The appropriate titer for each antibody was determined by staining control tissues with serial dilutions of each reagent between 0.0625–3 μg/mL. The final titer was chosen based on optimal signal-to-noise in these tissues.

To prepare the TB granuloma cohort, all specimens were first sectioned and mounted on slides. One section (5 μm) was placed on a glass slide, stained with hematoxylin-and-eosin (H&E), and inspected by an anatomic pathologist to screen for the presence of granulomatous inflammation. The second serially adjacent section (5 μm) was mounted on a MIBI-TOF slide (IonPath, Menlo Park, USA). For cohort staining, we reconstituted fresh aliquots of all antibodies, combined them into a single master mix, and stained all slides simultaneously. Interactive protocols for all these steps can be found at:

• Reagent preparation^52^

• IHC staining^53^

• MIBI staining^54^

• Sequenza staining^55^

• Antibody lyophilization^56^

• Lyophilized antibody reconstitution^57^

### MIBI-TOF data generation

All MIBI-TOF data was generated on a commercial MIBIScope instrument (IonPath, Menlo Park, USA). To image each ROI, we broke them down into 400 μm × 400 μm fields-of-view (FOVs). Each FOV was (1024 pixels)^2^ and were imaged using the “Coarse” settings on the MIBIScope instrument. Between all runs we conducted an imaging test on the molybdenum and polymethylmethacrylate (PMMA) standards per the manufacturer’s recommendations and settings. When the detector sensitivity dropped below the recommended levels, we adjusted the detector gain to minimize FOV-to-FOV sensitivity differences. One slide and one ROI was imaged at a time. However, to minimize image artifacts from instrument drift, we randomized the acquisition of each FOV within an ROI. All default image processing (background correction and denoising) were disabled. Low-level processing of the MIBI-TOF images was conducted with the publicly available, in-house pipeline, Toffy^58^.

### Single cell segmentation and phenotyping

To extract single cell data from the MIBI-TOF images, we utilized the image analysis pipeline, ark-analysis^59^. First, we used Mesmer to segment individual cells from all images. The CD45 and CD14 channels were combined and input as the cell membrane channel and the HH3, H3K9Ac, and H3K27me3 channels were combined to form a single nuclear channel. We used the default Mesmer parameters. To phenotype these cells, we next conducted pixel clustering with Pixie with the following parameters: blur_factor = 2 and subset_proportion = 0.1. We used the CD3, CD4, CD8a, CD11c, CD14, CD20, CD31, CD45, CD68, Vimentin, CD163, CD206, Calprotectin, Chymase/Tryptase, ɑSMA, DC-LAMP, Fibronectin, HLA-DR, and CD36 channels for clustering resulting in 22 metaclusters. We then used this information to quantify the proportion of each pixel cluster in each segmented cell object. These pixel cluster profiles were used to cluster the cells into 20 metaclusters. Giant cells were manually assigned their phenotype based on the previously described giant cell segmentation approach^32^. The resulting data provided a census of for each granuloma of individual cell phenotype and spatial positioning.

### *GranSim* model of lung granulomas

Immunobiological model of infection progression

*GranSim* is a multi-scale, hybrid agent-based model that simultaneously explores molecular, cellular, and tissue scales to describe complex biological processes that cannot be accurately captured at a single biological scale^60^. Our model has been published since 2004 and is continuously updated and curated with new biological datasets^16–18^. The hybrid agent-based model architecture allows us to compare model cell agents and cytokine levels to experimental images. *GranSim* tracks Mtb-infection associated activated Mtb-specific T cells (CD69^+^, T-bet, ROR-γT), CD8^+^ αβ T cells, IFN-γ cytokine signaling, and TNFα signaling along with other values such as chemokine diffusion and macrophage cell steps. Mycobacteria are represented in 3 different metabolic states within lung granulomas: non-replicating Mtb trapped within caseum necrosis, Mtb within macrophages (the preferred replicative niche for Mtb), and Mtb in extracellular spaces. *GranSim* tracks cytokine-pathway phosphorylated proteins at the molecular-scale, immune cell counts and bacterial counts at the cellular-scale, and spatial distributions of cells and macromolecules at the tissue-scale across time. The full list of rules and executable programs are available on the *GranSim* software website^61^.

Creation and validation of a virtual granuloma library

We begin a virtual granuloma library by simulating 20,000 *GranSim* granulomas sampled with 3 replicates across 63 parameter ranges to capture the heterogeneity observed in lung granulomas^19,20,23,62,63^ (Fig. 2B) including the variation of immune cell counts and CFU^19–22^ (Fig. 2A). In each simulation, we characterize mechanisms by a set of 63 biologically-relevant parameter values.

Intra-host and inter-host heterogeneity is caused by slight differences between the levels of activity of mechanisms at multiple scales, which *GranSim* captures by setting distinct parameter values (e.g., cell life lengths or cytokine clearance rates) for each mechanism within a biologically-relevant range. We establish *GranSim* parameter ranges from literature or experimental studies or by estimating them using model calibration^18,64,65^. We use Latin hypercube sampling (LHS), a stratified sampling-without-replacement technique^66–68^ to efficiently ensure a biologically heterogeneous virtual library (see Supplement on *GranSim* Rules, appendix B for details on the 63 varied parameters). We discard granulomas with extremely high CFU (>10^6^ CFU) that are typically not observed except during fulminant disease with uncontrolled pathology^23,69,70^, leaving a total of n=53,514 virtual granulomas.

To ensure that our virtual library is biologically plausible, we must quantitatively compare each virtual granuloma to the span of available biological data. To this end, we compare cell counts and CFU from each virtual granuloma against a longitudinal NHP dataset of the same. This dataset contains macrophage and T-cell counts (flow cytometry) and total CFU (growth assay) that have been collected over several years^19–22^ (Fig. 2A).

We classify Mtb infection in granulomas as either sterilizing, controlling, or uncontrolled using the CFU output by the *GranSim* model. Sterilizing granulomas are any granuloma whose CFU reaches 0. All other granulomas are either uncontrolled or controlling. As in ^71^, we assume that all granulomas with less than 10,000 CFU are controlling. Remaining granulomas are uncontrolled if they meet one of two criteria and controlling otherwise: (1) granulomas having more than a two-fold increase in CFU between days weeks 15 and 22 post-infection are uncontrolled, and (2) granulomas exceeding 100,000 CFU are uncontrolled. Therefore, if a non-sterile granuloma has less than a two-fold CFU increase from weeks 15 to 22 or has less than 10,000 CFU we classify that granuloma as controlling Mtb infection. We remove simulated granulomas that reach more than 10^6^ CFU at any timepoint to match the bacterial burden observed in experimental data^23,69,70^.

### Computational implementation

*GranSim* is programmed in portable C++14 code that executes on GNU/Linux and macOS operating systems. *GranSim* uses the Boost C++ library for general functionality^72^, the GNU Scientific Library (GSL) for linear algebra^73^, and the FFTW library for diffusion kinetics^74^. To visualize simulations, we use the Qt C++ library with its OpenGL module^75^. This work used *GranSim* r9959 compiled against Boost v1.79, GSL v2.7, FFTW v3.3.10, and Qt v5.12.8. Simulations with *GranSim*—especially the high number of simulations needed to sufficiently describe the large number of spatial features analyzed in this work—are computationally intensive and require HPC resources; we are granted time on national HPC systems to perform simulations for this work (see Acknowledgements).

### Spatial feature selection, extraction from imaging data, and correlation to infection state

#### Visualization of MIBI-TOF and *GranSim* TIFF images

TIFF composite color images were generated from TIFF binary channel masks described above of agent locations (*GranSim*) or cell locations (MIBI-TOF) using an ImageJ v1.54p^76^ macro provided in the supplementary code^77^. Image panels of MIBI-TOF with their digital partner images (Fig. 5 and Suppl. Fig. 12–Suppl. Fig. 21) were initially prototyped using FigureJ^78^ and then automated using the EBImage R package v4.48.0^79^ in the supplementary code^77^.

#### Selection of biologically-relevant candidate spatial features

We design our lists of spatial features of Mtb tissue pathology to measure known correlates of CFU in NHP Mtb granulomas in both the simulated and experimental images as described below; we do this to minimize the possibility of spurious spatial correlations from model implementation or non-Mtb pathological outcomes. From our work and the literature, known correlates for control of Mtb infection include IFN-γ, TNFα, IL-17 cytokines from proinflammatory CD4^+^ Th1 and Th17 cells^19,80–90^, and cytotoxic effector CD8^+^ αβ T cells^82,91^. The model directly simulates TNFα cytokines, INF-γ functionality with probabilistic interactions^61^, and simulates T cell agents equivalent to INF-γ producing proinflammatory CD4^+^ and CD8^+^ T cells T cells, cytotoxic effector T cells, and regulatory T regulatory cells (CD8^+^), though *GranSim* does not explicitly track IL-17. The MIBI-TOF immunolabels do not capture IL-17 or TNFα. Therefore, spatial measurements of these cell types and cytokines, described next, were chosen to correlate with Mtb infection tissue pathology. For more details about mapping immunolabels to *GranSim* agents see Suppl. Table 1.

We measure tissue-scale features from *GranSim* that are observable in the MIBI-TOF images (cluster number selected based on exploratory analysis (Suppl. Fig. 5–Suppl. Fig. 7). We use unsupervised clustering to identify temporal groupings among these features and then perform independent cross-sectional analyses at each week post-infection.

#### Using CellProfiler to compute spatial features from TIFF images

As an overview, we measure tissue scale features by examining *aggregates* that we construct for each cell type, intuitively similar to *channels* used for fluorescence images (see Extended Data 2; code available in supplementary materials). For example, the macrophage aggregate of a *GranSim* simulation is blank except for at locations containing macrophages. From this, we can derive measurements such as cellular distribution throughout granulomas, or how close the shape of a lymphocyte cuff is to a perfect circle (i.e., circularity). We then apply all cellular- and tissue-scale CellProfiler measurements to all cell types in *GranSim* that are also known correlates of bacterial burden.

For both *GranSim* and MIBI-TOF analyses, we use CellProfiler v4.2.6^25^, an object-based image analysis software. We use CellProfiler to read multiple TIFF images per-simulation or MIBI-TOF sample corresponding to different *GranSim* cell types or *GranSim* cytokine signals, or MIBI-TOF cell types, respectively. For each sample or simulation, we build (1) set of *objects* that record information about each individual cell (e.g., position) and (2) images that record chemical concentrations. We further process the individual cell objects into a single *aggregate* object per-cell-type (<u>Extended Data 2</u>). From the cell objects, aggregates, and images associated to a sample or simulation, we use CellProfiler to compute spatial relationships summarized in <u>Extended Data 7</u>.

#### Large-scale extraction and rescaling of spatial features from *GranSim* and MIBI-TOF data

From the virtual granuloma library, we use spatial information for every week up to 22 weeks post-infection as follows. Data files of agent locations and cytokine levels are converted into cell-identifying mask-integer-TIFF images and 32-bit floating point TIFF images, respectively. We do this using a purpose-built C program, gransimg^92^, using LibTIFF v4.5.1^93^ accelerated with OpenMP^94^. We analyze these TIFF images using CellProfiler as described above. We determine the neighbors of a cell *C* by identifying all cells within a 60 μm Chebyshev radius (using MeasureObjectNeighbors module in CellProfiler). This ensures that our results are comparable to those obtained using the 50 μm threshold used in published MIBI-TOF cell neighbor analyses; our 60 μm threshold is a multiple of the 20 μm grid compartment size used in *GranSim*.

We created and published a Spack^95^ package for CellProfiler v4.2.6 (that has since been accepted and merged into the Spack “builtin” repository) to analyze simulation TIFF images on the Purdue University Anvil HPC system^96^ and ran CellProfiler in batch mode followed by R post-processing using GNU Parallel v20220522^97^ in the job submission script. The R post-processing of CellProfiler individual cell object features consists of summarizing their mean, median, and interquartile range (IQR), as described in the last row of <u>Extended Data</u> <u>7</u>.

We run two nearly identical CellProfiler pipelines for *GranSim* and MIBI-TOF to accommodate their different image calibrations. The digitized MIBI-TOF cell centroids have a calibration of 0.390 μm per pixel. We round these to 1 μm per pixel to run CellProfiler on a reduced image size. The only time we need to control for the difference in these calibrations is when we perform optimal transport, where we rescale specific measurements from MIBI-TOF to *GranSim* as appropriate for the spatial feature measurement type (length, area, etc.).

The simulation CellProfiler pipeline, the MIBI-TOF CellProfiler pipeline, R code, and job submission scripts are included with the supplementary code^98^.

### Ranking experimental and virtual features using cross-sectional linear regression

#### Overview

Our goal is to identify the top spatial features that most highly correlate MIBI-TOF images to their log_10_(CFU + 1) levels (simplified as log-CFU). To do this, all spatial features of the virtual granulomas are measured at every week and at each of these weeks they are compared against their predicted log-CFU at 22 weeks. To identify the top spatial features that predict log-CFU, we begin with our 2,240 candidate spatial features most relevant to disease severity, discussed above, which we calculate for each granuloma in the virtual and experimental libraries. We then apply linear regression to determine the top predictors of CFU. Due to computational limitations described in the next paragraph, performing the linear regression requires two steps: firstly, computing the least absolute shrinkage and selection operator (LASSO) and secondly, computing the Bayesian variable selection. The regression is univariate, namely, as a primary indicator of infection severity, we only predict the log-CFU using spatial features for each virtual granuloma.

To be computationally tractable, our ranking of candidate digital partners requires examining virtual granulomas at a single representative week within the 9–12-week timespan of when the experimental granulomas were collected. This is because matching virtual granulomas to each MIBI-TOF sample at its own specific collection timepoint adds an additional time-dimension, and computing DPs using only one week took over 50 TB of space (see Methods). We choose week 11 to perform the matches because our exploratory feature clustering analyses (Suppl. Fig. 5–Suppl. Fig. 7) suggest that week 11 contains the most informative set of spatial features within the 9–12-week range. (Note that our exploratory analyses did not impose the same 4-week-contiguous criterion we imposed for our analysis in Fig. 3.) Additional support for our choice of using week 11 is that, in previous work from one of our groups, week 10 was considered a pivotal timepoint at which the balance between bacterial control and uncontrolled growth is determined in granulomas^82^.

#### Linear regression

We process each virtual granuloma week separately using LASSO regression analysis with the R v4.3.3 lars package v1.3^99^. However, querying for top predictive features using the lars package returns several additional predictive features than requested due to ties. Therefore, we break these ties using more computational expensive Bayesian variable selection with the projpred package v2.8.0^100^. Computational complexity imposes a practical limit against using Bayesian variable selection to subset more than 20 spatial features. Thus, from the 2,240 spatial features we applied both lars linear regression and projpred Bayesian variable selection to identify the top 20 spatial features that discriminate log-CFU infection severity.

#### Hierarchical clustering

We performed hierarchical clustering analysis on the top 20 spatial features for each week to determine potentially related sets of spatial features across the 22 weeks. We performed clustering (Suppl. Fig. 5–Suppl. Fig. 7) using the ggalign R package v1.2.0^101^ with the Canberra distance metric^102,103^. To use Canberra distance with ranking values, we first subtracted the ranks of their maximum value plus one so that rank 1 values became distance 20; this is necessary because distance metrics assume higher significance for higher values. Similarly, we assigned unranked predictive spatial features an infinite distance prior to the Canberra distance calculation.

#### Spatial feature contributions

To perform the next step of digital partner assignment of simulations to MIBI-TOF tissue samples, we use the calculated the expected log predictive density (ELPD)^104^ obtained during Bayesian variable selection (Suppl. Fig. 8–Suppl. Fig. 9). The ELPD is equivalent to minimizing the Kullback–Leibler divergence between the predictive distributions of the reference statistical model with all spatial features to the candidate submodel containing fewer spatial features. By ranking submodels that each successively add a single spatial variable leading up to the reference model with all spatial features, we thus estimate the contribution of each spatial feature by their difference in ELPD.

### Pairing each MIBI-TOF image with a unique simulation using unbalanced optimal transport

#### Overview

We obtain CDPs for each MIBI-TOF sample as follows. From the 25,503 non-sterilizing virtual granulomas, we select CDPs to be simulations with week-11 log-CFU similar to the that of the experimental sample. In Fig. 4, we compare MIBI-TOF log-CFU (single black dot) with CDP log-CFU (colored curves). We describe details of CDP selection in Methods. On average there are ∼3,500 virtual granulomas in the CDP set for each experimental granuloma. In general, the number of CDPs depends upon the parameter-density of the virtual library, the resolution of the experimental data, and the strictness of the similarity threshold. The precise number of CDPs is less important than the outcome heterogeneity of the most-similar CDPs. Using our method, 66% (21,251 of 32,101) were CDPs for at least one experimental library sample. The other 34% of virtual granulomas have substantially higher or lower CFU than the granulomas present in our small experimental library.

To rank similarity between MIBI-TOF samples and their CDPs, we use unbalanced optimal transport (OT). OT is a family of methods that match heterogeneous datasets by globally coupling two distributions and minimizing coupling costs^105–108^ (e.g., minimizing the amount of displacement required to transform one dataset into another). This method allows us to quantify how similar CDPs are to MIBI-TOF samples (Suppl. Fig. 11 ). For each granuloma, we select the top 50 ranked candidates as its final digital partner set (top three highlighted as black curves in Fig. 4).

#### Detailed procedure for ranking candidate digital partners

To find the closest matching simulations from the virtual library to each granuloma in the experimental library, we solve an unbalanced OT problem using all 28 MIBI-TOF granulomas to find their 28 corresponding *GranSim* granulomas. This type of OT problem is called unbalanced because there are more *GranSim* granulomas than MIBI-TOF granulomas—a balanced OT problem would have the same number of granulomas for both MIBI-TOF and *GranSim*. The OT maps the top spatial features (Extended Data 4) from MIBI-TOF to *GranSim* by setting the cost function with the ELPD obtained from the Bayesian variable selection step as described in the previous Methods subsection.

We express our granuloma spatial feature datasets with a linear programming formulation (eq. 1–eq. 4) of a classic transport problem (see Chapter 3 in ^109^). These problems map origins (*O*) to destinations (*D*), which are separated by an abstracted “distance”, computed using a given cost function *c*(*i*, *j*) where *i* ∈ *O* and *j* ∈ *D*. The goal of OT is to assign a pairing to each destination from a pre-specified number of origins (transport demand) in a way that minimizes total cost, i.e., ∑_j_ *c* (Origins of *j*, *j* ). Here, the origins are our simulations, and the destinations are the MIBI-TOF granulomas, and the objective function captures immunological similarity using the features identified in our Bayesian variable selection (computation of *c* is described later in this section). In our formulation, the destinations have a transport demand of one, namely, each destination MIBI-TOF sample can only accept one unique origin simulated granuloma to form the pairing (eq. 2). The origins have a transport supply of at most one, namely, each origin can send at most one simulated granuloma (i.e., parameter set) to form the pairing (eq. 3). Together these constraints ensure a one-to-one mapping (i.e., no simulated granuloma is matched to more than one MIBI-TOF granuloma).

We record our *O* → *D* pairing using a matching matrix [*x_i_*_,j_] where *x_i_*_,j_ is 1 *i* pairs to *j* and 0 otherwise. By assigning *x_i_*_,j_ as binary, we apply a redundant constraint to limit the solver from trying to assign the same origin simulated granulomas to a multiple destination MIBI-TOF granulomas. As with the classical transport problem, *x_i_*_,j_ is fixed to zero for any combination of origin and destination that is “forbidden” by external constraint criteria. Using this, we establish reasonable origins (i.e., the candidate digital partners) that may be paired to destinations (i.e., corresponding MIBI-TOF granulomas) by requiring origins to have log-CFU similar to their destinations according to Silverman’s bandwidth estimate^110^. (This was recently applied for OT pairing in super-resolution microscopy^111^.) In order to pose our pairing as a classic transport problem, we use a vector of spatial features (*F*), found earlier in our previous methods section of Bayesian variable selection, to build the cost matrix (*c*) using the source simulated *GranSim* granulomas (*G*) and destination MIBI-TOF granulomas (*M*).

In order to return a scalar value for *c*(*i*, *j*) using the vector *F*, we incorporate the features measured from both *M* and *G*, weighted by the ELPD for each feature (*E_k_* for *k* ∈ *F*) (eq. 4). Individually the dimensions of the matrices containing simulation and sample features are (#features × #simulations) and (#features × #*samples*), but it is more helpful to represent them as *G*^′^ and *M*^′^, both of dimension (#features × #simulations × #*samples*), where the added dimensions repeat their matrix contents or add zeros as appropriate. This allows us to match the dimensions of the feature matrices and, by assigning 0 as needed, consider only valid pairings. Among the 3 replicates of each *GranSim* granuloma (*G*) for each MIBI-TOF granuloma (*M*), we choose the replicate with the lowest cost, because some replicates of a given *GranSim* granuloma (*G*) may contain not-a-number values (NAs) in their spatial vector when they were analyzed by CellProfiler; in other words taking an average or other summary metric across the *GranSim* replicates would not work due to inconsistent dimensions. The cost matrix (*c*) is shown as a hierarchical cluster in Suppl. Fig. 11 .

The solution of equations eq. 1–eq. 4 yield a single optimal simulated pairing from *G* to each individual MIBI-TOF granuloma in *M*. To find next most similar digital partner, we use a process of successive elimination, where the optimal simulations (and replicates simulated using the same parameter values) are removed from *G* and the optimization is re-run to choose the next-most-similar set of simulations. We do this to generate and rank the 50 closest features for each sample in *M*.

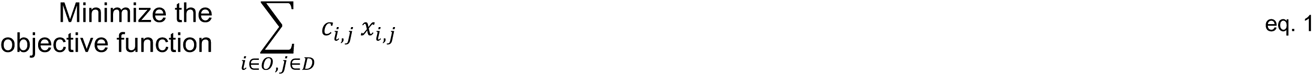

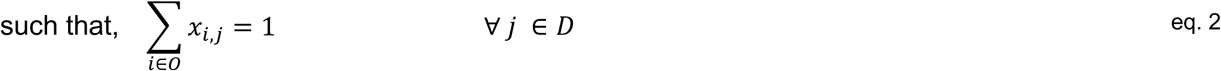

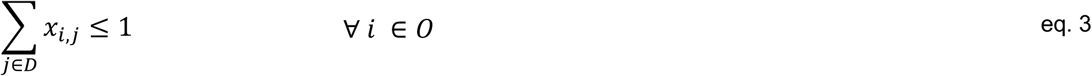

Where,

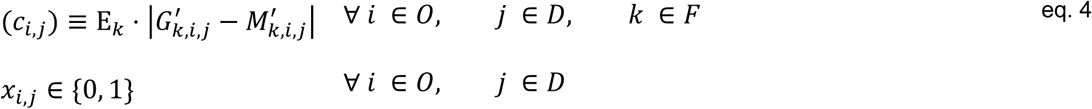

Note that the MIBI-TOF granulomas have a temporal resolution of 1 day, but we group all 9–12-week granulomas as 11 weeks to reduce complexity as mentioned in the Results as follows. Modifying two steps in the analysis would make it possible to use the 1-day resolution of MIBI-TOF: first, all equations eq. 1–eq. 4 would require adding a time dimension as well as an additional constraint equation to only choose one pairing among all timepoints. Second, we would need to simulate virtual granulomas at all timepoints corresponding to those for which we have MIBI-TOF samples; as a practical note, CellProfiler unsummarized per-cell metrics introduce data storage challenges including our HPC cluster data storage quotas.

We use the GNU Linear Programming Kit (GLPK) via the Rglpk R package version 0.6-5.1^112^ to solve equations eq. 1–eq. 4; in particular, the sparse matrix support available in Rglpk is necessary for memory efficiency. Note that the R implementation simplifies the 3 dimensions formulated above to 2 dimensions with detailed explanations in the R code comments^77^. Additionally, the R code to reproduce the digital partners Fig. 4 in the paper includes a minimal example using toy data (available in the supplementary code^77^ in the section 5.1 “Toy example of digital partners”).

In the future, we can apply an unbalanced OT approach from a purely mathematical to a hybrid mathematical-statistical framework by retaining our current CFU-matching approach while improving spatial feature matching through regularized, scale-invariant optimal transport methods like those in Python Optimal Transport^113^. We expect that our approach (experimentally informed subsampling of model simulations using unbalanced OT) will both improve the predictive power of computational intervention studies and enable the incorporation of additional multi-modal datasets.

### Sensitivity analysis of model simulations matching MIBI-TOF images

Due to the complex nonlinear interactions between *GranSim* mechanisms, standard correlations provide limited insight (Suppl. Fig. 22). To identify key factors driving behaviors of the system we use partial rank correlation coefficients (PRCC), a sensitivity analysis approach refined by our group^68^. We first use LHS to sample the parameter space capturing uncertainty of the model system based on parameters. For the ∼1,100 digitally partnered virtual granulomas (Fig. 6 and Extended Data 6), we test vary 63 model parameters characterizing mechanisms of interest against 7 model outputs. We then use PRCC to identify non-linear correlations between parameter values and simulation outcomes. Parameters with the highest PRCC values are the most influential over the observed outcomes, and this allows us to identify which mechanisms drive infection outcomes. We identify which PRCC values are largest using the boxplot whisker outlier test in R via the boxplot.stats built-in function, where the whiskers are computed as the sum of 1.5 times the interquartile range and Tukey’s hinge^114,115^. All other sensitivity analysis calculations are run with MATLAB R2023b.

For direct comparison between drivers of indeterminate granulomas and granulomas with consensus predictions, we use the top 184 digital partners for each of the 6 indeterminate parameter samples (yielding 1,104 total, comparable to the 1,100 analyzed in Fig. 6). Beyond the parameters we identify in Fig. 6 for the CDPs that were consistent, this PRCC analysis uncovers additional drivers of heterogeneity specific to the indeterminate granulomas. (Extended Data 6 visualizes these correlations.)

### Reproducibility

For reproducibility, all analyses that generate Fig. 1–Fig. 6 are made available in the uploaded supplementary code and data^77,98^; this is available on GitHub at https://github.com/kirschner-lab/spatialmatch. Briefly, the GitHub repository contains two major sets of supplemental code that are required to reproduce all figures, tables, and supplementary data. The primary set of code is a collection of R scripts that perform the novel procedures for the *Tissue Time Machine* approach, namely CDP identification using optimal transport. The README file indicates which specific files generate each figure. These R scripts comprise the end-phase analyses shown in this manuscript and require inputs that come from a secondary set of code.

This secondary set of code contains the specific commands to run *GranSim*, convert the raw *GranSim* outputs into TIFF images, compute their spatial features using CellProfiler, then performs LASSO and Bayesian variable selection. The GitHub README describes each step of the workflow, and the associated code is in a .zip file archived in Zenodo. Running the secondary code set takes nearly a million hours of compute time to generate so, we have also included a Zenodo archive that includes the relevant outputs of the secondary code set used for generating Fig. 1–Fig. 6 (located at ^98^).

All figures use the more accessible color-blindness friendly Okabe–Ito color palette^116^ built into R; the raster images were not remapped to Okabe–Ito but the legibility of the raster images is minimally impacted because red hues are never used with only blue-green-white colors in use.

## Notes

### Competing Interest Statement

The authors have declared no competing interest.

## References

1. Van de Sande B, Lee JS, Mutasa-Gottgens E, Naughton B, Bacon W, Manning J, et al. Applications of single-cell RNA sequencing in drug discovery and development. Nat Rev Drug Discov. 2023 Jun;22(6):496–520. doi:10.1038/s41573-023-00688-4

2. Chen TY, You L, Hardillo JAU, Chien MP. Spatial Transcriptomic Technologies. Cells. 2023 Jan;12(16):2042. doi:10.3390/cells12162042

3. Giesen C, Wang HAO, Schapiro D, Zivanovic N, Jacobs A, Hattendorf B, et al. Highly multiplexed imaging of tumor tissues with subcellular resolution by mass cytometry. Nat Methods. 2014 Apr;11(4):417–22. doi:10.1038/nmeth.2869 PubMed PMID: 24584193.

4. Lin JR, Izar B, Wang S, Yapp C, Mei S, Shah PM, et al. Highly multiplexed immunofluorescence imaging of human tissues and tumors using t-CyCIF and conventional optical microscopes. eLife. 2018 Jul 11;7:e31657. doi:10.7554/eLife.31657 PubMed PMID: 29993362; PubMed Central PMCID: PMC6075866.

5. Black S, Phillips D, Hickey JW, Kennedy-Darling J, Venkataraaman VG, Samusik N, et al. CODEX multiplexed tissue imaging with DNA-conjugated antibodies. Nat Protoc. 2021 Aug;16(8):3802–35. doi:10.1038/s41596-021-00556-8 PubMed PMID: 34215862; PubMed Central PMCID: PMC8647621.

6. Liu CC, Bosse M, Kong A, Kagel A, Kinders R, Hewitt SM, et al. Reproducible, high-dimensional imaging in archival human tissue by multiplexed ion beam imaging by time-of-flight (MIBI-TOF). Laboratory Investigation. 2022 Jul;102(7):762–70. doi:10.1038/s41374-022-00778-8 PubMed PMID: 35351966; PubMed Central PMCID: PMC10357968.

7. Global Tuberculosis Report 2024. 1st ed. Geneva: World Health Organization; 2024. 1 p.

8. Bold TD, Ernst JD. Who benefits from granulomas, mycobacteria or host? Cell. 2009 Jan 9;136(1):17–9. doi:10.1016/j.cell.2008.12.032S0092-8674(08)01638-3 [pii] PubMed PMID: 19135882.

9. Davis JM, Ramakrishnan L. The role of the granuloma in expansion and dissemination of early tuberculous infection. Cell. 2009 Jan 9;136(1):37–49. doi:S0092-8674(08)01443-8 [pii] 10.1016/j.cell.2008.11.014 PubMed PMID: 19135887.

10. Ramakrishnan L. Revisiting the role of the granuloma in tuberculosis. Nat Rev Immunol. 2012 May;12(5):352–66. doi:10.1038/nri3211 PubMed PMID: 22517424.

11. Liu H, Li P, He S, Yang H, Yuan T, He X, et al. The role and mechanisms of multiple immunoregulatory cells in pulmonary tuberculosis. Front Immunol. 2026 Jun 2;17:1841581. doi:10.3389/fimmu.2026.1841581

12. Flynn JL, Chan J. Immune cell interactions in tuberculosis. Cell. 2022 Dec 8;185(25):4682–702. doi:10.1016/j.cell.2022.10.025 PubMed PMID: 36493751.

13. McCaffrey EF, Delmastro AC, Fitzhugh I, Ranek JS, Douglas S, Peters JM, et al. The immunometabolic topography of cellular organization and bacterial control in tuberculosis granulomas. Nat Immunol. 2026 Apr;27(4):867–80. doi:10.1038/s41590-026-02431-8 PubMed PMID: 41731147; PubMed Central PMCID: PMC13361869.

14. Michael CT, Almohri SA, Linderman JJ, Kirschner DE. A framework for multi-scale intervention modeling: virtual cohorts, virtual clinical trials, and model-to-model comparisons. Frontiers in Systems Biology. 2024;3. doi:10.3389/fsysb.2023.1283341 PubMed PMID: 39310676.

15. Sinha A, Weichhart T. Spatial immunometabolic zonation in tuberculosis granulomas. Nat Rev Immunol. 2025 Sep;25(9):635–635. doi:10.1038/s41577-025-01210-0 PubMed PMID: 40624263.

16. Segovia-Juarez JL, Ganguli S, Kirschner D. Identifying control mechanisms of granuloma formation during M. tuberculosis infection using an agent-based model. Journal of Theoretical Biology. 2004;231(3):357–76. doi:10.1016/j.jtbi.2004.06.031 PubMed PMID: 15501468.

17. Warsinske HC, DiFazio RM, Linderman JJ, Flynn JL, Kirschner DE. Identifying mechanisms driving formation of granuloma-associated fibrosis during Mycobacterium tuberculosis infection. Journal of Theoretical Biology. 2017 Jun 19. doi:10.1016/j.jtbi.2017.06.017 PubMed PMID: 28642013.

18. Nanda P, Budak M, Michael CT, Krupinsky K, Kirschner DE. Development and Analysis of Multiscale Models for Tuberculosis: From Molecules to Populations. In: Aguiar M, Bellomo N, Chaplain M, editors. Predicting Pandemics in a Globally Connected World, Volume 2 [Internet]. Cham: Springer Nature Switzerland; 2024 [cited 2026 Apr 4]. p. 11–43. (Modeling and Simulation in Science, Engineering and Technology). Available from: https://link.springer.com/10.1007/978-3-031-56794-0_2 doi:10.1007/978-3-031-56794-0_2

19. Gideon HP, Phuah J, Myers AJ, Bryson BD, Rodgers MA, Coleman MT, et al. Variability in tuberculosis granuloma T cell responses exists, but a balance of pro- and anti-inflammatory cytokines is associated with sterilization. PLoS Pathog. 2015 Jan;11(1):e1004603. doi:10.1371/journal.ppat.1004603 PubMed PMID: 25611466.

20. Martin CJ, Cadena AM, Leung VW, Lin PL, Maiello P, Hicks N, et al. Digitally Barcoding Mycobacterium tuberculosis Reveals In Vivo Infection Dynamics in the Macaque Model of Tuberculosis. MBio. 2017 May 9;8(3). doi:10.1128/mBio.00312-17 PubMed PMID: 28487426.

21. Darrah PA, DiFazio RM, Maiello P, Gideon HP, Myers AJ, Rodgers MA, et al. Boosting BCG with proteins or rAd5 does not enhance protection against tuberculosis in rhesus macaques. NPJ Vaccines. 2019;4:21. doi:10.1038/s41541-019-0113-9 PubMed PMID: 31149352; PubMed Central PMCID: PMC6538611.

22. Joslyn LR, Linderman JJ, Kirschner DE. A virtual host model of Mycobacterium tuberculosis infection identifies early immune events as predictive of infection outcomes. Journal of Theoretical Biology. 2022 Apr 21;539:111042. doi:10.1016/j.jtbi.2022.111042 PubMed PMID: 35114195; PubMed Central PMCID: Pmc9169921.

23. Lin PL, Ford CB, Coleman MT, Myers AJ, Gawande R, Ioerger T, et al. Sterilization of granulomas is common in active and latent tuberculosis despite within-host variability in bacterial killing. Nat Med. 2014 Jan;20(1):75–9. doi:10.1038/nm.3412 PubMed PMID: 24336248.

24. Lin PL, Flynn JL. The End of the Binary Era: Revisiting the Spectrum of Tuberculosis. J Immunol. 2018 Nov 1;201(9):2541–8. doi:10.4049/jimmunol.1800993 PubMed PMID: 30348659; PubMed Central PMCID: PMC6217958.

25. Stirling DR, Swain-Bowden MJ, Lucas AM, Carpenter AE, Cimini BA, Goodman A. CellProfiler 4: improvements in speed, utility and usability. BMC Bioinformatics. 2021 Dec;22(1):433. doi:10.1186/s12859-021-04344-9 PubMed PMID: 34507520; PubMed Central PMCID: PMC8431850.

26. Millar JA, Butler JR, Evans S, Grant NL, Mattila JT, Linderman JJ, et al. Spatial Organization and Recruitment of Non-Specific T Cells May Limit T Cell-Macrophage Interactions Within Mycobacterium tuberculosis Granulomas. Front Immunol. 2021 Jan 20;11. doi:10.3389/fimmu.2020.613638 PubMed PMID: 33552077; PubMed Central PMCID: PMC7855029.

27. Gong C, Mattila JT, Miller M, Flynn JL, Linderman JJ, Kirschner D. Predicting lymph node output efficiency using systems biology. Journal of Theoretical Biology. 2013 Oct 21;335:169–84. doi:10.1016/j.jtbi.2013.06.016 PubMed PMID: 23816876; PubMed Central PMCID: PMC3783027.

28. Eileen A. Wong, Louis Joslyn, Nicole Grant, Edwin Klein, Philana Ling Lin, Denise Kirschner, et al. Low Levels of T Cell Exhaustion in Tuberculous Lung Granulomas. Infection and Immunity. 2018. doi:10.1128/IAI.00426-18 PubMed PMID: 29891540.

29. Cadena AM, Hopkins FF, Maiello P, Carey AF, Wong EA, Martin CJ, et al. Concurrent infection with Mycobacterium tuberculosis confers robust protection against secondary infection in macaques. PLOS Pathogens. 2018;14(10):e1007305. doi:10.1371/journal.ppat.1007305 PubMed PMID: 30312351; PubMed Central PMCID: PMC6200282.

30. Wong EA, Evans S, Kraus CR, Engelman KD, Maiello P, Flores WJ, et al. IL-10 Impairs Local Immune Response in Lung Granulomas and Lymph Nodes during Early Mycobacterium tuberculosis Infection. Vol. 204. 2020 Feb 1;204(3):644–59. doi:10.4049/jimmunol.1901211 PubMed PMID: 31862711.

31. Bromley JD, Ganchua SKC, Nyquist SK, Maiello P, Chao M, Borish HJ, et al. CD4+ T cells re-wire granuloma cellularity and regulatory networks to promote immunomodulation following *Mtb* reinfection. Immunity. 2024 Oct 8;57(10):2380–2398.e6. doi:10.1016/j.immuni.2024.08.002 PubMed PMID: 39214090.

32. McCaffrey EF, Donato M, Keren L, Chen Z, Delmastro A, Fitzpatrick MB, et al. The immunoregulatory landscape of human tuberculosis granulomas. Nat Immunol. 2022 Feb;23(2):318–29. doi:10.1038/s41590-021-01121-x PubMed PMID: 35058616; PubMed Central PMCID: PMC8810384.

33. Joslyn LR, Renardy M, Weissman C, Grant NL, Flynn JL, Butler JR, et al. Temporal and Spatial Analyses of TB Granulomas to Predict Long-Term Outcomes. In: Vodovotz Y, An G, editors. Complex Systems and Computational Biology Approaches to Acute Inflammation [Internet]. Cham: Springer International Publishing; 2021 [cited 2026 Sep 15]. p. 273–91. Available from: http://link.springer.com/10.1007/978-3-030-56510-7_15 doi:10.1007/978-3-030-56510-7_15

34. Lyu J, Narum DE, Baldwin SL, Larsen SE, Bai X, Griffith DE, et al. Understanding the development of tuberculous granulomas: insights into host protection and pathogenesis, a review in humans and animals. Front Immunol. 2024 Dec 9;15:1427559. doi:10.3389/fimmu.2024.1427559 PubMed PMID: 39717773; PubMed Central PMCID: PMC11663721.

35. Flores-Gonzalez J, Ramón-Luing LA, Falfán-Valencia R, Batista CVF, Soto-Alvarez S, Huerta-Nuñez L, et al. The presence of cytotoxic CD4 and exhausted-like CD8+ T-cells is a signature of active tuberculosis. Biochimica et Biophysica Acta (BBA) - Molecular Basis of Disease. 2024 Aug;1870(6):167219. doi:10.1016/j.bbadis.2024.167219 PubMed PMID: 38734321.

36. Shen X, Wu T, Ji X, Yang K, Wang L, Peng Y, et al. Mycobacterium tuberculosis infection depressed cytotoxic T cells activity owing to decreasing NKG2C and increasing NKG2A expression. Molecular Immunology. 2023 Oct;162:133–42. doi:10.1016/j.molimm.2023.08.014 PubMed PMID: 37683324.

37. Wei H, Wang R, Yuan Z, Chen CY, Huang D, Halliday L, et al. DR*W201/P65 Tetramer Visualization of Epitope-Specific CD4 T-Cell during M. tuberculosis Infection and Its Resting Memory Pool after BCG Vaccination. Pai M, editor. PLoS ONE. 2009 Sep 4;4(9):e6905. doi:10.1371/journal.pone.0006905 PubMed PMID: 19730727; PubMed Central PMCID: PMC2731856.

38. Marino S, Hult C, Wolberg P, Linderman JJ, Kirschner DE. The Role of Dimensionality in Understanding Granuloma Formation. Computation (Basel). 2018 Dec;6(4):58. doi:10.3390/computation6040058 PubMed PMID: 31258937; PubMed Central PMCID: PMC6599587.

39. Renardy M, Wessler T, Blemker S, Linderman J, Peirce S, Kirschner D. Data-Driven Model Validation Across Dimensions. Bulletin of mathematical biology. 2019 Jun;81(6):1853–66. doi:10.1007/s11538-019-00590-4 PubMed PMID: 30830675; PubMed Central PMCID: PMC6494696.

40. Laubenbacher R, Mehrad B, Shmulevich I, Trayanova N. Digital twins in medicine. Nat Comput Sci. 2024 Mar;4(3):184–91. doi:10.1038/s43588-024-00607-6 PubMed PMID: 38532133.

41. Masison J, Beezley J, Mei Y, Ribeiro H, Knapp AC, Sordo Vieira L, et al. A modular computational framework for medical digital twins. Proceedings of the National Academy of Sciences. 2021 May 18;118(20):e2024287118. doi:10.1073/pnas.2024287118 PubMed PMID: 33972437; PubMed Central PMCID: PMC8157963.

42. Knapp AC, Cruz DA, Mehrad B, Laubenbacher RC. Personalizing computational models to construct medical digital twins. bioRxiv. 2024 Nov 7;2024.05.31.596692. doi:10.1101/2024.05.31.596692 PubMed PMID: 39574674; PubMed Central PMCID: PMC11580862.

43. Kimpton LM, Paun LM, Colebank MJ, Volodina V. Challenges and opportunities in uncertainty quantification for healthcare and biological systems. Philosophical Transactions of the Royal Society A: Mathematical, Physical and Engineering Sciences. 2025 Mar 13;383(2292):20240232. doi:10.1098/rsta.2024.0232 PubMed PMID: 40078151; PubMed Central PMCID: PMC11904623.

44. Laubenbacher R, Adler F, An G, Castiglione F, Eubank S, Fonseca LL, et al. Toward mechanistic medical digital twins: some use cases in immunology. Front Digit Health. 2024 Mar 7;6:1349595. doi:10.3389/fdgth.2024.1349595 PubMed PMID: 38515550; PubMed Central PMCID: PMC10955144.

45. Wong HS, Park K, Gola A, Baptista AP, Miller CH, Deep D, et al. A local regulatory T cell feedback circuit maintains immune homeostasis by pruning self-activated T cells. Cell. 2021 Jul;184(15):3981–3997.e22. doi:10.1016/j.cell.2021.05.028 PubMed PMID: 34157301; PubMed Central PMCID: PMC8390950.

46. Hickey JW, Agmon E, Horowitz N, Tan TK, Lamore M, Sunwoo JB, et al. Integrating multiplexed imaging and multiscale modeling identifies tumor phenotype conversion as a critical component of therapeutic T cell efficacy. Cell Systems. 2024 Apr;15(4):322–338.e5. doi:10.1016/j.cels.2024.03.004 PubMed PMID: 38636457; PubMed Central PMCID: PMC11030795.

47. Schiebinger G, Shu J, Tabaka M, Cleary B, Subramanian V, Solomon A, et al. Optimal-Transport Analysis of Single-Cell Gene Expression Identifies Developmental Trajectories in Reprogramming. Cell. 2019 Feb;176(4):928–943.e22. doi:10.1016/j.cell.2019.01.006 PubMed PMID: 30712874; PubMed Central PMCID: PMC6402800.

48. Tong A, Huang J, Wolf G, van Dijk D, Krishnaswamy S. TrajectoryNet: A Dynamic Optimal Transport Network for Modeling Cellular Dynamics. Proc Mach Learn Res. 2020 Jul;119:9526–36. PubMed PMID: 34337419; PubMed Central PMCID: PMC8320749.

49. Huguet G, Magruder DS, Tong A, Fasina O, Kuchroo M, Wolf G, et al. Manifold Interpolating Optimal-Transport Flows for Trajectory Inference. Adv Neural Inf Process Syst. 2022 Dec;35:29705–18. PubMed PMID: 37397786; PubMed Central PMCID: PMC10312391.

50. Sha Y, Qiu Y, Zhou P, Nie Q. Reconstructing growth and dynamic trajectories from single-cell transcriptomics data. Nat Mach Intell. 2023 Nov 30;6(1):25–39. doi:10.1038/s42256-023-00763-w PubMed PMID: 38274364; PubMed Central PMCID: PMC10805654.

51. Marino S, Kirschner DE. A Multi-Compartment Hybrid Computational Model Predicts Key Roles for Dendritic Cells in Tuberculosis Infection. Computation (Basel). 2016 Oct 21;4(4). doi:10.3390/computation4040039 PubMed PMID: 28989808; PubMed Central PMCID: PMC5627612.

52. Mb Bosse M. MIBI and IHC solutions v1 [Internet]. 2020 [cited 2026 Aug 17]. Available from: https://www.protocols.io/view/mibi-and-ihc-solutions-bhmej43e doi:10.17504/protocols.io.bhmej43e

53. Mb Bosse M. IHC staining v1 [Internet]. 2020 [cited 2026 Aug 17]. Available from: https://www.protocols.io/view/ihc-staining-bf6ajrae doi:10.17504/protocols.io.bf6ajrae

54. Mb Bosse M. MIBI staining v2 [Internet]. 2022 [cited 2026 Aug 17]. Available from: https://www.protocols.io/view/mibi-staining-b9b3r2qn doi:10.17504/protocols.io.dm6gprk2dvzp/v5

55. Mb Bosse M, Bendall S, Angelo M. Staining Sequenza v1 [Internet]. 2020 [cited 2026 Aug 17]. Available from: https://www.protocols.io/view/staining-sequenza-bmc6k2ze doi:10.17504/protocols.io.bmc6k2ze

56. Camacho C, Mb Bosse M, Bendall S, Angelo M. Antibody lyophilization v1 [Internet]. 2020 [cited 2026 Aug 17]. Available from: https://www.protocols.io/view/antibody-lyophilization-bhmgj43w doi:10.17504/protocols.io.bhmgj43w

57. Mb Bosse M, Angelo M, Bendall S. Lyophilized metal-antibody reconstitution v1 [Internet]. 2020 [cited 2026 Aug 17]. Available from: https://www.protocols.io/view/lyophilized-metal-antibody-reconstitution-bjd6ki9e doi:10.17504/protocols.io.bjd6ki9e

58. angelolab. GitHub [Internet]. [cited 2026 Aug 17]. GitHub - angelolab/toffy: Scripts for interacting with and generating data from the commercial MIBIScope. Available from: https://github.com/angelolab/toffy

59. ark-analysis: Integrated pipeline for multiplexed image analysis [Internet]. GitHub: Angelo Lab. Available from: https://github.com/angelolab/ark-analysis

60. Gammack D, Ganguli S, Marino S, Segovia-Juarez J, Kirschner DE. Understanding the immune response in tuberculosis using different mathematical models and biological scales. Multiscale Model Sim. 2005;3(2):312–45. doi:10.1137/040603127

61. GranSim [Internet]. Denise Kirschner Laboratory at the University of Michigan Medical School; 2025. Available from: http://malthus.micro.med.umich.edu/GranSim/

62. Flynn JL, Gideon HP, Mattila JT, Lin PL. Immunology studies in non-human primate models of tuberculosis. Immunol Rev. 2015 Mar;264(1):60–73. doi:10.1111/imr.12258 PubMed PMID: 25703552.

63. Cadena AM, Fortune SM, Flynn JL. Heterogeneity in tuberculosis. Nature reviews Immunology. 2017 Nov;17(11):691–702. doi:10.1038/nri.2017.69 PubMed PMID: 28736436.

64. Joslyn L, Kirschner D, Linderman JJ. CaliPro: A calibration protocol that utilizes parameter density estimation to explore parameter space and calibrate complex biological models. Cel Mol Bioeng. 2020 Sep 15;31–47. doi:10.1007/s12195-020-00650-z PubMed PMID: 33643465; PubMed Central PMCID: PMC7878626.

65. Nanda P, Kirschner DE. Calibration methods to fit parameters within complex biological models. Front Appl Math Stat. 2023;9. doi:10.3389/fams.2023.1256443 PubMed PMID: 38222943; PubMed Central PMCID: PMC10785782.

66. M. D. McKay, R. J. Beckman, W. J. Conover. A comparison of three methods for selecting values of input variables in the analysis of output from a computer code. Technometrics. 2000;42(1):55–61. doi:10.2307/1271432

67. Helton JC, Johnson JD, Sallaberry CJ, Storlie CB. Survey of sampling-based methods for uncertainty and sensitivity analysis. Reliability Engineering & System Safety. 2006 Oct 1;The Fourth International Conference on Sensitivity Analysis of Model Output (SAMO 2004)91(10):1175–209. doi:10.1016/j.ress.2005.11.017

68. Marino S, Hogue IB, Ray CJ, Kirschner DE. A methodology for performing global uncertainty and sensitivity analysis in systems biology. J Theor Biol. 2008 Sep 7;254(1):178–96. doi:10.1016/j.jtbi.2008.04.011 PubMed PMID: 18572196; PubMed Central PMCID: PMC2570191.

69. Maiello P, DiFazio RM, Cadena AM, Rodgers MA, Lin PL, Scanga CA, et al. Rhesus Macaques Are More Susceptible to Progressive Tuberculosis than Cynomolgus Macaques: a Quantitative Comparison. Infect Immun. 2018 Feb;86(2). doi:10.1128/IAI.00505-17 PubMed PMID: 28947646; PubMed Central PMCID: PMC5778369.

70. Hult C, Mattila JT, Gideon HP, Linderman JJ, Kirschner DE. Neutrophil Dynamics Affect Mycobacterium tuberculosis Granuloma Outcomes and Dissemination. Front Immunol. 2021;12:712457. doi:10.3389/fimmu.2021.712457 PubMed PMID: 34675916; PubMed Central PMCID: PMC8525425.

71. Budak M, Via LE, Weiner DM, Barry III CE, Nanda P, Michael G, et al. A systematic efficacy analysis of tuberculosis treatment with BPaL-containing regimens using a multiscale modeling approach. CPT: Pharmacometrics & Systems Pharmacology. 2024;13(4):673–85. doi:10.1002/psp4.13117 PubMed PMID: 38404200.

72. Abrahams D, Dawes B, Rivera R. Boost C++ Libraries [Internet]. 1998. Available from: https://www.boost.org

73. Galassi M, Theiler J, Gough B, Jungman G. GNU Scientific Library [Internet]. 2001. Available from: https://www.gnu.org/software/gsl/

74. Frigo M, Johnson SG. Fastest Fourier Transform in the West (FFTW) [Internet]. 2003. Available from: https://www.fftw.org

75. Qt [Internet]. The Qt Company; 1995. Available from: https://www.qt.io

76. Schneider CA, Rasband WS, Eliceiri KW. NIH Image to ImageJ: 25 years of image analysis. Nat Methods. 2012 Jul;9(7):671–5.

77. spatialmatch: Figures and code for ABM spatial matching paper [Internet]. GitHub; 2025. Available from: https://github.com/kirschner-lab/spatialmatch

78. Mutterer J, Zinck E. Quick-and-clean article figures with FigureJ. Journal of Microscopy. 2013 Oct;252(1):89–91. doi:10.1111/jmi.12069 PubMed PMID: 23906423.

79. Pau G, Fuchs F, Sklyar O, Boutros M, Huber W. EBImage—an R package for image processing with applications to cellular phenotypes. Bioinformatics. 2010 Apr 1;26(7):979–81. doi:10.1093/bioinformatics/btq046 PubMed PMID: 20338898; PubMed Central PMCID: PMC2844988.

80. Lin PL, Plessner HL, Voitenok NN, Flynn JL. Tumor necrosis factor and tuberculosis. J Investig Dermatol Symp Proc. 2007 May;12(1):22–5.

81. Lin PL, Myers A, Smith L, Bigbee C, Bigbee M, Fuhrman C, et al. Tumor necrosis factor neutralization results in disseminated disease in acute and latent Mycobacterium tuberculosis infection with normal granuloma structure in a cynomolgus macaque model. Arthritis Rheum. 2010 Feb;62(2):340–50. doi:10.1002/art.27271 PubMed PMID: 20112395.

82. Gideon HP, Hughes TK, Tzouanas CN, Wadsworth MH, Tu AA, Gierahn TM, et al. Multimodal profiling of lung granulomas in macaques reveals cellular correlates of tuberculosis control. Immunity. 2022 May;55(5):827–846.e10. doi:10.1016/j.immuni.2022.04.004 PubMed PMID: 35483355; PubMed Central PMCID: PMC9122264.

83. Algood HM, Lin PL, Flynn JL. Tumor necrosis factor and chemokine interactions in the formation and maintenance of granulomas in tuberculosis. Clin Infect Dis. 2005 Aug 1;41 Suppl 3:S189-93. doi:CID35769 [pii] 10.1086/429994 PubMed PMID: 15983898.

84. Khader SA, Bell GK, Pearl JE, Fountain JJ, Rangel-Moreno J, Cilley GE, et al. IL-23 and IL-17 in the establishment of protective pulmonary CD4+ T cell responses after vaccination and during Mycobacterium tuberculosis challenge. Nat Immunol. 2007 Apr;8(4):369–77.

85. Millington KA, Innes JA, Hackforth S, Hinks TSC, Deeks JJ, Dosanjh DPS, et al. Dynamic Relationship between IFN-γ and IL-2 Profile of *Mycobacterium tuberculosis* -Specific T Cells and Antigen Load. The Journal of Immunology. 2007 Apr 15;178(8):5217–26. doi:10.4049/jimmunol.178.8.5217 PubMed PMID: 17404305; PubMed Central PMCID: PMC2743164.

86. Green AM, Difazio R, Flynn JL. IFN-gamma from CD4 T cells is essential for host survival and enhances CD8 T cell function during Mycobacterium tuberculosis infection. J Immunol. 2013 Jan 1;190(1):270–7. doi:10.4049/jimmunol.1200061 PubMed PMID: 23233724.

87. O’Garra A, Redford PS, McNab FW, Bloom CI, Wilkinson RJ, Berry MP. The immune response in tuberculosis. Annu Rev Immunol. 2013;31:475–527. doi:10.1146/annurev-immunol-032712-095939 PubMed PMID: 23516984.

88. Lyadova IV. Neutrophils in Tuberculosis: Heterogeneity Shapes the Way? Mediators of inflammation. 2017;2017:8619307. doi:10.1155/2017/8619307 PubMed PMID: 28626346.

89. Scriba TJ, Coussens AK, Fletcher HA. Human Immunology of Tuberculosis. Microbiol Spectr. 2017 Jan;5(1). doi:10.1128/microbiolspec.TBTB2-0016-2016 PubMed PMID: 28155806.

90. Lyadova IV, Panteleev AV. Th1 and Th17 Cells in Tuberculosis: Protection, Pathology, and Biomarkers. Hwang SA, editor. Mediators of Inflammation. 2015 Jan;2015(1):854507. doi:10.1155/2015/854507 PubMed PMID: 26640327; PubMed Central PMCID: PMC4657112.

91. Esaulova E, Das S, Singh DK, Choreño-Parra JA, Swain A, Arthur L, et al. The immune landscape in tuberculosis reveals populations linked to disease and latency. Cell Host & Microbe. 2021 Feb;29(2):165–178.e8. doi:10.1016/j.chom.2020.11.013 PubMed PMID: 33340449; PubMed Central PMCID: PMC7878437.

92. Nanda P. gransimg: C program to export GranSim TIFF image output [Internet]. GitHub; 2024. Available from: https://github.com/kirschner-lab/gransimg

93. libtiff: TIFF Library and Utilities [Internet]. GitLab; 2023. Available from: https://gitlab.com/libtiff/libtiff

94. Dagum L, Menon R. OpenMP: an industry standard API for shared-memory programming. IEEE Comput Sci Eng. 1998 Mar;5(1):46–55. doi:10.1109/99.660313

95. Gamblin T, LeGendre M, Collette MR, Lee GL, Moody A, De Supinski BR, et al. The Spack package manager: bringing order to HPC software chaos. In: Proceedings of the International Conference for High Performance Computing, Networking, Storage and Analysis [Internet]. Austin Texas: ACM; 2015 [cited 2026 Aug 17]. p. 1–12. Available from: https://dl.acm.org/doi/10.1145/2807591.2807623 doi:10.1145/2807591.2807623

96. Song XC, Smith P, Kalyanam R, Zhu X, Adams E, Colby K, et al. Anvil - System Architecture and Experiences from Deployment and Early User Operations. In: Practice and Experience in Advanced Research Computing [Internet]. Boston MA USA: ACM; 2022 [cited 2026 Aug 17]. p. 1–9. Available from: https://dl.acm.org/doi/10.1145/3491418.3530766 doi:10.1145/3491418.3530766

97. Tange O. GNU Parallel 20220522 (’NATO’) [Internet]. Zenodo; 2022 [cited 2026 Aug 17]. Available from: https://zenodo.org/record/6570228 doi:10.5281/ZENODO.6570228

98. Nanda P. Simulated and processed data input for publication figures generated by the spatialmatch R package [Internet]. Zenodo; 2025 [cited 2026 Aug 17]. Available from: https://zenodo.org/doi/10.5281/zenodo.15102620 doi:10.5281/ZENODO.15102620

99. Efron B, Hastie T, Johnstone I, Tibshirani R. Least angle regression. Ann Statist. 2004 Apr 1;32(2). doi:10.1214/009053604000000067

100. Piironen J, Paasiniemi M, Vehtari A. Projective inference in high-dimensional problems: Prediction and feature selection. Electron J Statist. 2020 Jan 1;14(1). doi:10.1214/20-EJS1711

101. Peng Y, Wang S, Yu G. ggalign: A ggplot2 Extension for Consistent Axis Alignment [Internet]. CRAN; 2025. Available from: https://CRAN.R-project.org/package=ggalign

102. Lance GN, Williams WT. Computer Programs for Hierarchical Polythetic Classification (“Similarity Analyses”). The Computer Journal. 1966 May 1;9(1):60–4. doi:10.1093/comjnl/9.1.60

103. Lance GN, Williams WT. Mixed-Data Classificatory Programs I - Agglomerative Systems. Aust Comput J. 1967;1(1):15–20.

104. Vehtari A, Ojanen J. A survey of Bayesian predictive methods for model assessment, selection and comparison. Statist Surv. 2012 Jan 1;6(none). doi:10.1214/12-SS102

105. Monge G. Mémoire sur la théorie des déblais et des remblais. 1781.

106. Brenier Y. Polar factorization and monotone rearrangement of vector-valued functions. Comm Pure Appl Math. 1991 Jun;44(4):375–417. doi:10.1002/cpa.3160440402

107. Villani C. Topics in optimal transportation. Reprinted with corrections. Providence, Rhode Island: American Mathematical Society; 2016. 378 p. (Graduate studies in mathematics; no. 58).

108. Kantorovich LV. On the Translocation of Masses. J Math Sci. 2006 Mar;133(4):1381–2. doi:10.1007/s10958-006-0049-2

109. Fourer R, Gay DM, Kernighan BW. AMPL: a modeling language for mathematical programming. 2. ed. San Francisco, Calif.: Scientific Pr.; 1996.

110. Silverman BW. Density Estimation for Statistics and Data Analysis. Boca Raton: Routledge; 2018. 1 p. (Chapman and Hall/CRC Monographs on Statistics and Applied Probability; v. 26).

111. Naas J, Nies G, Li H, Stoldt S, Schmitzer B, Jakobs S, et al. MultiMatch: geometry-informed colocalization in multi-color super-resolution microscopy. Commun Biol. 2024 Sep 13;7(1):1139. doi:10.1038/s42003-024-06772-8 PubMed PMID: 39271907; PubMed Central PMCID: PMC11399439.

112. Theussl S, Hornik K. Rglpk: R/GNU Linear Programming Kit Interface [Internet]. CRAN; 2024. Available from: https://CRAN.R-project.org/package=Rglpk

113. Flamary R, Courty N, Gramfort A, Alaya MZ, Boisbunon A, Chambon S, et al. POT: Python Optimal Transport. Journal of Machine Learning Research. 2021;22(78):1–8.

114. Mcgill R, Tukey JW, Larsen WA. Variations of Box Plots. The American Statistician. 1978 Feb;32(1):12–6. doi:10.1080/00031305.1978.10479236

115. Chambers JM, editor. Graphical methods for data analysis. 6. [print.]. Pacific Grove, Calif: Wadsworth & Brooks/Cole; 1983. 395 p. (The Wadsworth & Brooks/Cole statistics probability series).

116. Okabe M, Ito K. Color universal design (CUD): How to make figures and presentations that are friendly to color blind people [Internet]. 2002 Nov 20. Available from: https://jfly.uni-koeln.de/html/color_blind/

