## Extended Data for "A tissue time machine to forward-predict single-timepoint data applied to tuberculosis granulomas"

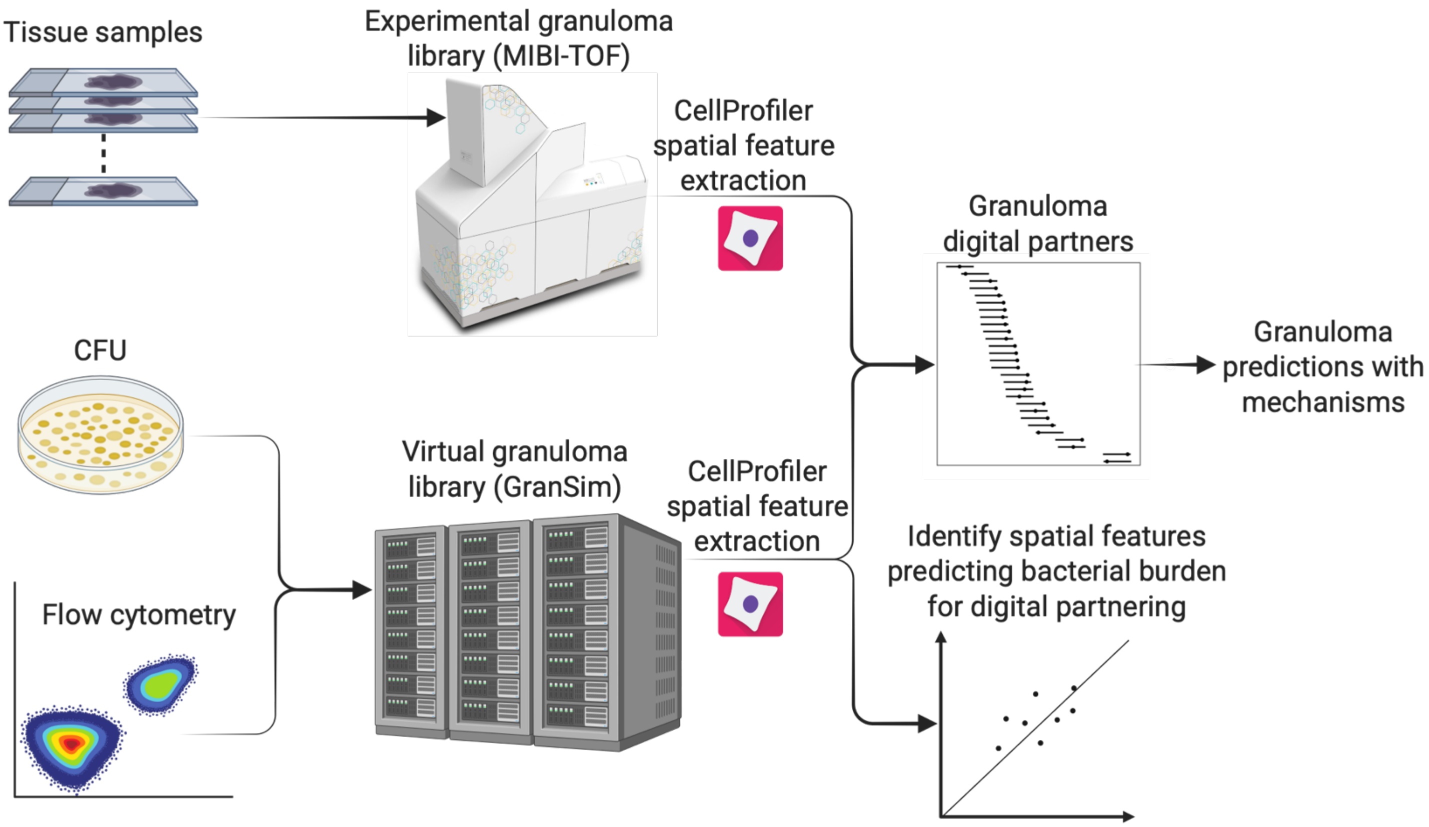


Extended Data 1 **Overview of hybrid experimental-computational workflow**. We use virtual granulomas to predict infection outcomes of 28 MIBI-TOF-imaged granulomas. We immunolabel lung granuloma tissue samples from cynomolgus macaques with metal isotope-tagged antibodies for immunohistochemistry and imaging with an IonPath MIBI-TOF imager. For each NHP granuloma, curated CFU and historical flow cytometry datasets are available; we used these data to calibrate *GranSim* and then generated a virtual granuloma library of 60,000 granulomas simulated for 22 weeks post-infection. We discarded simulations that resulted in biologically implausible CFU (>$10^6$ CFU) for a final virtual granuloma library of n = 53,514 granulomas. We used CellProfiler to extract 2,240 spatial features for both experimental and virtual granulomas. We then use virtual granuloma CFU outcomes (clearance, controlled, and uncontrolled growth) to rank spatial features most predictive of CFU. We identify virtual granulomas from the library as candidate digital partners based on the proximity between their simulated log-CFU and the experimental granuloma's measured log-CFU. We then rank candidate digital partners by optimal-transport distance across selected Mtb-relevant spatial features. These top-ranked digital partners allow us to both predict infection outcomes and identify mechanistic drivers of granuloma infection progression. Created in BioRender. Nanda, P. (2026) https://BioRender.com/ouex42d

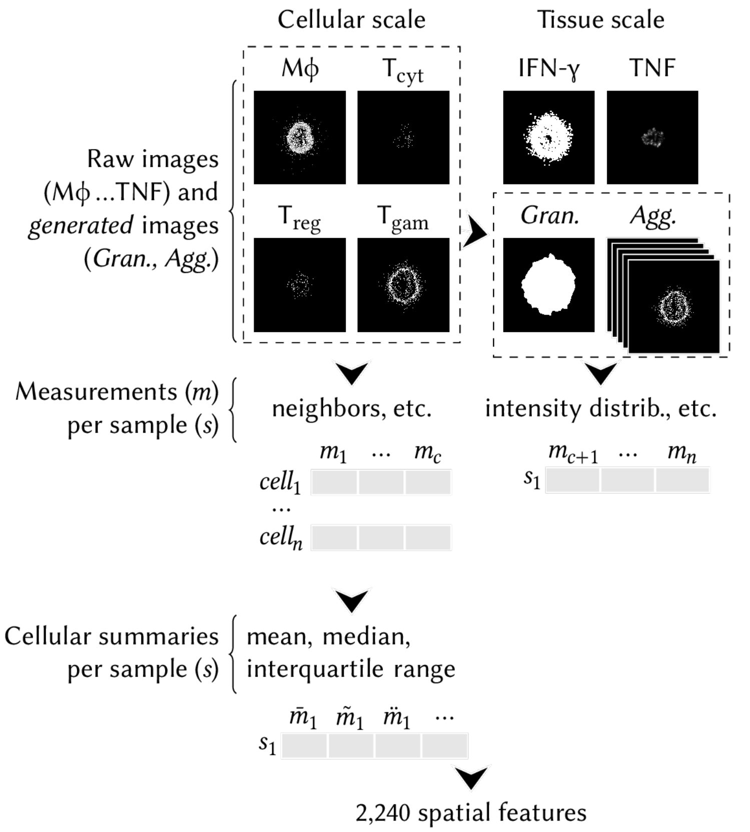


Extended Data 2 **CellProfiler workflow for extracting spatial features from MIBI-TOF experimental granulomas and *GranSim* virtual granulomas**. We aggregate all cellular-scale channels into tissue-scale features to capture both tissue-scale cell distributions and cellular-scale relationships. We create the granuloma composite (*Gran.*) by combining all macrophages and T cells, applying dilation, and filling holes to delineate the granulomatous region. From the cellular-scale images (outlined by dashed lines), we derive aggregated (*Agg.*) images, where measurements are applied across all cells of a given type. We also derive the granuloma-extent mask (*Gran.*) from cellular-scale images and create an additional aggregated image combining all T-cell subtypes. We extract all features using CellProfiler modules described in Extended Data 4. We summarize cellular-scale features as scalar statistics (mean, median, and interquartile range). Example images are from virtual granuloma 988. Minor differences between the virtual and experimental pipelines ensure comparability in downstream processing (see Methods). Abbreviations used in image labels: Mϕ = macrophage, $T_{cyt}$ = cytotoxic $CD8^+$ effector T cell, $T_{reg}$ = regulatory $CD4^+$ T cell, $T_{gam}$ = IFN-γ-producing T cell.

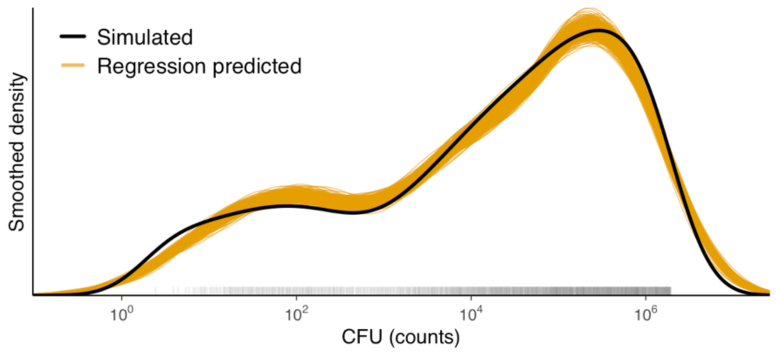


Extended Data 3 **As few as 12 of 2240 Mtb-relevant spatial features from the virtual library are sufficient to predict the distribution of simulated log-CFU at 22 weeks post-infection**. The thick black curve shows the smoothed density of CFU values across non-sterilizing granulomas in the virtual library (subsampled to 1,000 granulomas for visualization; CFU values are the rug-plot ticks on the X-axis). The thin orange curves show 1,000 predicted CFU distributions generated by Bayesian linear regression using only the 12 spatial features in Extended Data 4. Each orange curve represents one plausible regression model drawn from the posterior distribution, i.e., one set of regression coefficients sampled from the range of coefficient values consistent with the actual simulated CFU values. For each linear model, we predict CFU for the same 1,000 granulomas using only their spatial features, then plot the resulting density. The tight overlap among orange curves indicates low uncertainty in the regression coefficients, and their close match to the black curve demonstrates that 12 spatial features are sufficient to recapitulate the full distribution of bacterial burden. Y-axis values are omitted because density plots are normalized to integrate to 1; absolute Y-axis values are very small due to the X-axis values spanning six orders of magnitude.

| Scale | Spatial feature statistic | R | Coef. | SE |
|---|---|---|---|---|
| Cellular | **Angle formed between a macrophage and its two closest macrophage neighbors†:** Relative to a specified macrophage *M*, we measure the angle subtended at *M* by the closest pair of other macrophages within 60 μm* of *M,* or 0 if no other neighbors are nearby. A higher value indicates the closest macrophage neighbors of *M* are distributed toward opposing directions (i.e., in a line); a lower value may indicate that *M* is uniformly surrounded by other macrophages regardless of local macrophage density; a lower value may also indicate that the macrophage neighbors of *M* are all on one side of *M* (e.g., *M* may be flush against caseum necrosis). *In vivo*, this measurement is likely to be slightly different due to not being constrained to a 2-D grid, although the interpretation of the measured values translates between the virtual and experimental contexts. | 0.93 | $3.09\times10^{-2}$ | $1.12\times10^{-3}$ |
| | **Angle formed between a cytotoxic T cell and its two closest cytotoxic T-cell neighbors‡**: Computed exactly as above but instead using cytotoxic T cells. | 0.76 | $1.35\times10^{-3}$ | $9.92\times10^{-5}$ |
| Tissue | **IFN-γ$^+$-T cell aggregate major axis length.** Higher values indicate a larger lymphocytic cuff. | 0.96 | $6.83\times10^{-3}$ | $2.73\times10^{-4}$ |
| | **IFN-γ$^+$-T cell aggregate minor axis length.** Higher values indicate a larger lymphocytic cuff. | 0.94 | $8.69\times10^{-3}$ | $2.76\times10^{-4}$ |
| | **Outer granuloma macrophage activation signal**: Fraction of IFN-γ at the outermost quarter ring of the connected area occupied by macrophages. Higher values indicate that macrophage-colocalized IFN-γ is further from the granuloma center. | 0.94 | 3.71 | $9.22\times10^{-2}$ |
| | **1 / (Number of macrophages in simulation).** Higher values indicate fewer total macrophages. | -0.91 | -486 | 12.6 |
| | **1 / (Number of IFN-γ$^+$-T cells in simulation).** Higher values indicate fewer total IFN-γ$^+$ T cells. | -0.87 | -86.7 | 10.1 |
| | **T-cell aggregate shape-irregularity.** Measure of non-circularity of the aggregate of T cells (including all subtypes). where a minimum value of 1 represents exactly a circle, and higher values are more irregular. Higher values indicate a less circular lymphocytic cuff. | -0.86 | -0.652 | $2.18\times10^{-1}$ |
| | **Inner granuloma macrophage activation signal**: Fraction of IFN-γ at the innermost quarter ring of the connected area occupied by macrophages. Higher values indicate more IFN-γ in macrophages at the center of the granuloma. | -0.86 | -12.6 | $8.55\times10^{-1}$ |
| | **Cytotoxic T cell aggregate diameter.** | 0.86 | $1.54\times10^{-2}$ | $4.97\times10^{-4}$ |
| | **Regulatory T cell aggregate diameter.** | 0.84 | $1.87\times10^{-2}$ | $5.40\times10^{-4}$ |
| | **Macrophage aggregate radius.** | 0.73 | 0.588 | $1.68\times10^{-2}$ |

Extended Data 4 **Minimal set of 12 spatial features out of 2,240 CellProfiler spatial features that recapitulate week-22 CFU distributions**. Using LASSO and Bayesian linear regressions, we find two cellular-scale and ten tissue-scale features that collectively predict log-CFU in non-sterilizing virtual granulomas (see Methods and Suppl. Fig. 8–Suppl. Fig. 10). The R values are calculated against log-CFU. The cellular-scale measurements are computed for the indicated cell types, i.e., macrophages and cytotoxic T cells. We report the final fitted Bayesian linear model regression coefficients (Coef.) and standard error (SE); the linear intercept coefficient and standard error are -5.29 and 0.103, respectively. Spatial features are grouped by scale and ordered by absolute R value within each group. * We chose 60 μm as the neighborhood boundary to be comparable to the 50 μm boundary used for previous MIBI-TOF (see Methods)**.** † Mean across all macrophages. ‡ Median across all cytotoxic T cells.

| MIBI-TOF sample | Rank 1 | Rank 2 | Rank 3 | Agreement |
|---|---|---|---|---|
| **Group 1: Samples with top-3 digital partners in consensus (n=22/28)** | | | | |
| 4 | Controlling | Controlling | Controlling | Controlling |
| 6 | Controlling | Controlling | Controlling | Controlling |
| 8 | Controlling | Controlling | Controlling | Controlling |
| 9 | Controlling | Controlling | Controlling | Controlling |
| 11 | Controlling | Controlling | Controlling | Controlling |
| 12 | Controlling | Controlling | Controlling | Controlling |
| 17 | Controlling | Controlling | Controlling | Controlling |
| 20 | Controlling | Controlling | Controlling | Controlling |
| 21 | Controlling | Controlling | Controlling | Controlling |
| 26 | Controlling | Controlling | Controlling | Controlling |
| 28 | Controlling | Controlling | Controlling | Controlling |
| 29 | Controlling | Controlling | Controlling | Controlling |
| 38 | Controlling | Controlling | Controlling | Controlling |
| 41 | Controlling | Controlling | Controlling | Controlling |
| 42 | Controlling | Controlling | Controlling | Controlling |
| 48 | Controlling | Controlling | Controlling | Controlling |
| 50 | Controlling | Controlling | Controlling | Controlling |
| 54 | Controlling | Controlling | Controlling | Controlling |
| 55 | Controlling | Controlling | Controlling | Controlling |
| 59 | Controlling | Controlling | Controlling | Controlling |
| 32 | Uncontrolled | Uncontrolled | Uncontrolled | Uncontrolled |
| 33 | Uncontrolled | Uncontrolled | Uncontrolled | Uncontrolled |
| **Group 2: samples with top-3 digital partners not in consensus (n=6/28)** | | | | |
| 23 | Controlling | Uncontrolled | Uncontrolled | Indeterminate |
| 34 | Controlling | Controlling | Uncontrolled | Indeterminate |
| 35 | Uncontrolled | Uncontrolled | Controlling | Indeterminate |
| 40 | Controlling | Controlling | Sterile | Indeterminate |
| 53 | Controlling | Sterile | Controlling | Indeterminate |
| 57 | Controlling | Uncontrolled | Uncontrolled | Indeterminate |

Extended Data 5 **Top 3 digital partner outcomes and agreement**. Correspond outcomes for Suppl. Fig. 12–Suppl. Fig. 21 and overall agreement among the 3 digital partner outcomes. Approx. 80% (22/28) are predictive using the time machine. Within those that agree, 71.43% (n = 20) are controlling and 7.14% (n = 2) uncontrolled, 21.43% are indeterminate (i.e., a mix of controlling and uncontrolled) (n = 6/28).

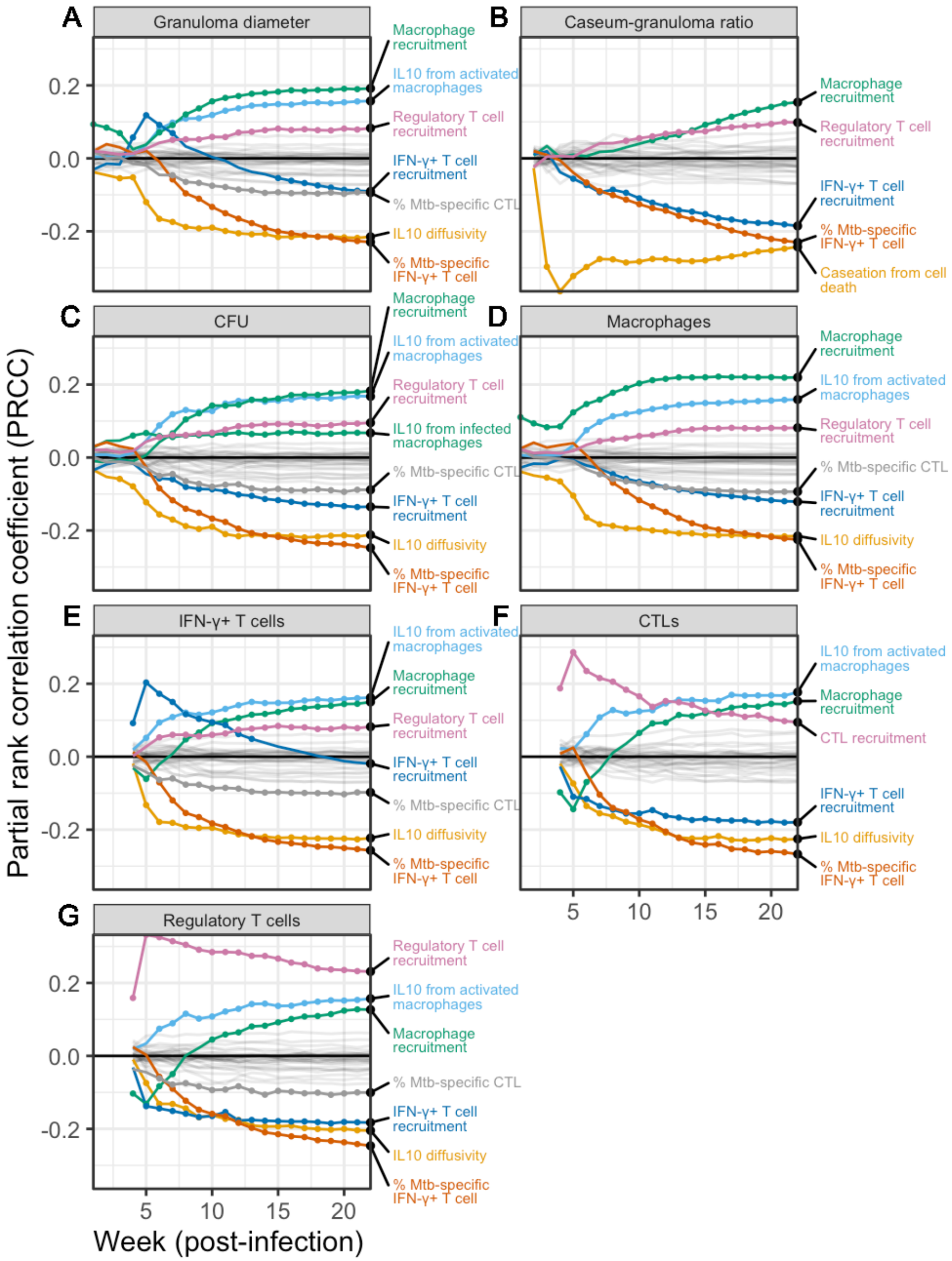


Extended Data 6 **Identification of significant mechanisms driving tissue-scale features, CFU, and immune cell levels during infection for the 6/28 indeterminate samples.** Shown are partial-rank correlation coefficients (PRCCs) derived from uncertainty and sensitivity analysis (U/SA) of the 184 closest ranked digital partners for each of the 6 MIBI-TOF tissue samples without a consensus predicted outcome (second group in Extended Data 5). We vary model parameters and calculate PRCCs driving tissue-scale outputs (granuloma diameter and caseous-to-cellular-area ratios) and cellular-scale outputs at each week from 1-22 for all digital partners (184 for each sample, yielding a total of n = 1,104 digital partners). Influential mechanisms for each outcome were identified using the boxplot whisker outlier test (see Methods) and are shown as colored curves. Timepoints of those impactful mechanisms with p-value < 0.05, corrected by the Benjamini–Hochberg false discovery rate (BH-FDR), are shown as colored dots along their respective curves. Non-influential reference parameters are shown as semi-transparent gray curves and not labeled.

| Category | Scale | Processing using CellProfiler (list of CellProfiler Modules) |
|---|---|---|
| Preprocessing | Tissue | Create tissue objects for each cell type by merging cell object identifier raw data (SplitOrMergeObjects x6) |
| Preprocessing | Tissue | Create T cell tissue object by merging T cell types (CombineObjects x2, SplitOrMergeObjects) |
| Preprocessing | Tissue | Create granuloma tissue object by merging T cell tissue object with macrophage tissue object (CombineObjects, SplitOrMergeObjects, ConvertObjectsToImage, DilateImage) |
| Quality Control | Tissue | Flag multiple granulomas and choose only single-necrotic core granulomas to match MIBI-TOF samples (IdentifyPrimaryObjects, FlagImage) |
| Preprocessing | Tissue | Choose largest granuloma if granuloma has multiple necrotic cores to avoid error (MeasureObjectSizeShape, FilterObjects, FlagImage) |
| Measurement | Cellular | Cytokine intensities in cells (MeasureObjectIntensity) |
| Measurement | Tissue | Cytokine intensity distribution (MeasureObjectIntensity, MeasureObjectIntensityDistibution) |
| Measurement | Both | Aggregate arrangements of cells (MeasureObjectSizeShape) |
| Measurement | Cellular | Cell neighbors (MeasureObjectNeighbors x27) |
| Aggregation | Cellular | Per-cell statistics for each image using mean, median, and IQR (N/A; used an R script) |

Extended Data 7 **Overview of CellProfiler pipeline processing steps of *GranSim* and MIBI-TOF images at each timepoint**. We provide the CellProfiler pipeline files in the supplementary code and data for specifics of how the CellProfiler modules were configured. There are minor differences between the two pipelines (simulated versus experimental) to control for pixel size differences (20 μm per pixel for *GranSim*; 1 μm per pixel for MIBI-TOF), to ensure that binary dilation of cell masks generate tissue-scale masks appropriately, and to convert pixel sizes to make the features comparable in the downstream optimal transport R code. See the Zenodo link to the supplementary HPC code[100].
