## Supplementary Material for "A tissue time machine to forward-predict single-timepoint data applied to tuberculosis granulomas"

#### Extended biological background for TB

Granuloma heterogeneity

Host and pathogen-driven mechanisms interact in complex ways to affect granuloma outcomes of bacterial clearance, control, or overgrowth[1–5]. Immune profiles in the blood do not accurately reflect the lung battle[6,7], thus translational studies of TB outcomes often focus on granulomas.

Prior work has identified that the spatial organization of TB granulomas is critical to limit Mtb growth and dissemination[8–11], likely driven by a balance of pro- and anti-inflammatory mediators[12]. Canonically, granulomas have infected necrotic centers (caseum) encircled by macrophages and other myeloid cells (termed the myeloid core). Most T cells are excluded from the myeloid core, instead surrounding it to form the lymphocytic cuff. These mediators include $CD4^+$ and $CD8^+$ T-cell levels, macrophage differentiation states, proinflammatory cytokine TNFα and IFN-γ, anti-inflammatory cytokine IL-10, lipid mediators, and co-expression of $IDO^+$ and PD-L1 in myeloid cells, and others[8,13–15].

Use of NHP animal models

Acquiring human pulmonary granulomas is challenging and invasive. When human TB specimens are biopsied, there is limited clinical information (i.e., bacterial burden, granuloma age, granuloma trajectory) that can be obtained per specimen. In this work, we examine individual granulomas both from NHPs and simulated using *GranSim*. Experimentally, the cynomolgus macaque NHP model of TB is a powerful tool to investigate the important immunobiological mechanisms that determine these distinct outcomes[16].

The NHP model is the only animal model that (1) forms human-like Mtb granulomas in response to infection and (2) fully reproduces the spectrum of infection outcomes observed in humans: clearance, clinical latency, and active disease[17–19]. Adult primate hosts infected with low-dose Mtb (< 25 colony-forming units or CFU) typically form 8–20 pulmonary granulomas, each with its own outcome trajectory[13,19–22]. A single uncontrolled granuloma is sufficient for a whole animal to progress toward active infection[13,21]. Each granuloma within a host follows its own infection trajectory ranging from complete sterilization to high bacterial burden and dissemination when the immune system fails to contain Mtb[3,23].

Additional biological context regarding Fig. 6 and Extended Data 4

There are similarities between both the spatial features predictive of granuloma outcomes and temporal mechanisms correlating with digital partner outcomes. Notably, we find uncontrolled granulomas are associated with less T-cell penetration into the center of granulomas, as high CFU is predicted by (1) macrophage activation being predominantly relegated to the outermost portion of granulomas and (2) more circular lymphocyte cuffs, which we have observed previously[20]. This complements earlier studies using single-cell RNA sequencing that found that macrophage-produced IFN-γ was associated with high-CFU granulomas at 10 weeks post-infection[24]. When considering both spatial and temporal analyses together, we find that reduced CFU is associated with more effective macrophage activation close to the granuloma center: CFU negatively correlates with both IFN-$\gamma^+$ T-cell Mtb-specificity and IFN-γ-macrophage colocalization near the granuloma center.

#### Extended Results

Comparison between spatial feature clusters (Fig. 3)

In our simulations, all three clusters in Fig. 3 involve pro-inflammatory activity of macrophages. The transient cluster includes a radial measure of macrophages near IFN-γ, positioned away from macrophage–Mtb interaction sites; this pattern was recently identified by immunotopography[25]. The post-formation clusters (transient and maintenance) include IFN-γ-producing T cells, cytotoxic $CD8^+$ T cells, and regulatory $CD4^+$ T cells. Cytotoxic $CD8^+$ T cells play a protective role in human anti-Mtb immunity and are significantly reduced in active TB patients[26]. Consistent with this, single-cell analysis of NHP granulomas that found that cytotoxic $CD8^+$ T cells are associated with low bacterial burden[24]. Our analysis of the virtual granuloma library identifies spatial features corresponding to prominent markers of infection previously reported in human and NHP literature[20,24,27–35] (see Methods).

Correlation between CFU and granuloma size and shape (Extended Data 4)

We also find that the granuloma size and shape have a complex relationship to CFU (Extended Data 4). Highlighting the importance of granuloma size, high-CFU granulomas are associated with large numbers of both macrophages and IFN-γ$^+$ T cells. Similarly, high-CFU granulomas are associated with larger diameters of macrophage and T-cell aggregates (including cytotoxic, regulatory, and IFN-γ$^+$). This suggests that the increased bacterial burden is driving increased cell recruitment or proliferation at the granuloma site, leading to larger granulomas as noted previously[36]. For granuloma shape, we find that granulomas with irregularly shaped T-cell aggregates (i.e., in contrast to relatively intact circular lymphocytic cuffs) correlate with lower bacterial burdens. On the cellular scale, larger angles between cytotoxic cells (i.e., both macrophages and cytotoxic T cells) and their closest same-type neighbors are related to higher CFU. Larger angles are indicative of cells arranged linearly along corridor-like shapes, possibly due to irregularly shaped caseum (e.g., the uncontrolled granuloma in Fig. 2B) as previously hypothesized[10]. Taken together, these results suggest that poorly-controlling granulomas become large and have T cells relegated to a more circular lymphocytic cuff, despite any local irregularities emerging from caseum. These simulated results highlight the importance of using both cellular- and tissue-scale features to predict infection outcomes.

Extended discussion of mechanisms driving granuloma- and cell-scale outcomes (Fig. 6)

The number of macrophages and the rate of activated macrophage production of IL-10 is associated with increased regulatory T cell levels and increased granuloma diameters (Fig. 6). Higher IL-10 diffusivity blunts this effect, likely due to weaker localized signals leading to less regulatory T cell recruitment. Granuloma caseum-to-cellular area ratio is primarily driven by the caseation rate (Fig. 6B), which characterizes the number of macrophages that must die to caseate a 20 μm × 20 μm area. Once adaptive immunity strengthens (beginning at 4 weeks post-infection and increasing thereafter), there are more Mtb-specific IFN-γ$^+$ T cells, leading to increased T-cell-induced macrophage apoptosis. This leads to decreased caseum formation (negative correlation with IFN-γ$^+$ T cells). The frequency of Mtb-specificity among IFN-γ$^+$ T cells is a negative correlate of both granuloma diameter and caseum-to-cellular area ratio, and this correlation becomes stronger over time.

Macrophage recruitment rate emerges as a significant driver increasing all tissue- and cellular-scale outputs over time, including both pro- and anti-inflammatory T cell levels. The only exception to this is during the formation phase (weeks 1-4), where macrophage recruitment negatively correlates with cytotoxic T cell counts (Fig. 6F). This reinforces the dual role of macrophages in Mtb infection[37,38].

Most of the same mechanisms drive cellular-scale outcomes (Fig. 6C–G). All cellular-scale outputs except CFU have one or more parameters that undergo statistically significant sign changes predominantly during the transient phase (weeks 5-14; Fig. 3). Parameters that positively correlate with total CFU, regardless of metabolic state, are related to macrophages and IL-10 levels. Mtb specificity among IFN-γ$^+$ T cells is a negative correlate of CFU throughout both the transient and maintenance phases, unlike its correlation with granuloma diameter. Elevated IL-10 and macrophage recruitment are also associated with larger cytotoxic and IFN-γ$^+$ T cell populations, indicative of ongoing infection[11,35,39,40]. In all cases, recruitment and activity of Mtb-specific IFN-γ$^+$ T cells negatively correlate with all outcomes except regulatory T cell levels (even that of IFN-γ$^+$-producing T cells themselves) as the immune response is self-limiting.

#### *GranSim* agent-to-cell mapping

Not all *GranSim* agents are identical to cell markers and instead simulates function-based phenotypes as shown in Suppl. Table 1 and Suppl. Fig. 1. Additionally, *GranSim* has simulated agents that were added for specific projects, meaning those project-specific agents have individually been used for past investigations. However, project-specific agents have not been fully calibrated together and therefore were not enabled for the simulations presented in this work.

Suppl. Table 1: **Grouping of MIBI-TOF cell phenotypes to *GranSim* agents**. * These *GranSim* project-specific agents were not simulated in this work.

| **Cell phenotype** | **Agent** |
|---|---|
| CD11c$^+$ Mac | Macrophage |
| CD14$^+$CD11c$^+$ Mac | Macrophage |
| CD14$^+$ Mac/Mono | Macrophage |
| CD163$^+$ Mac | Macrophage |
| CD203$^+$ Mac | Macrophage |
| CD68$^+$ Mac | Macrophage |
| CD4$^+$ Tcell | Regulatory T cell |
| CD8$^+$ Tcell | Cytotoxic T cell |
| HLADR$^+$ APC | Dendritic cell * |

| | |
|---|---|
| Neutrophil | Neutrophil * |
| FN1$^{+}$ Fibroblast | Fibroblast * |
| SMA$^{+}$ Fibroblast | Myofibroblast * |

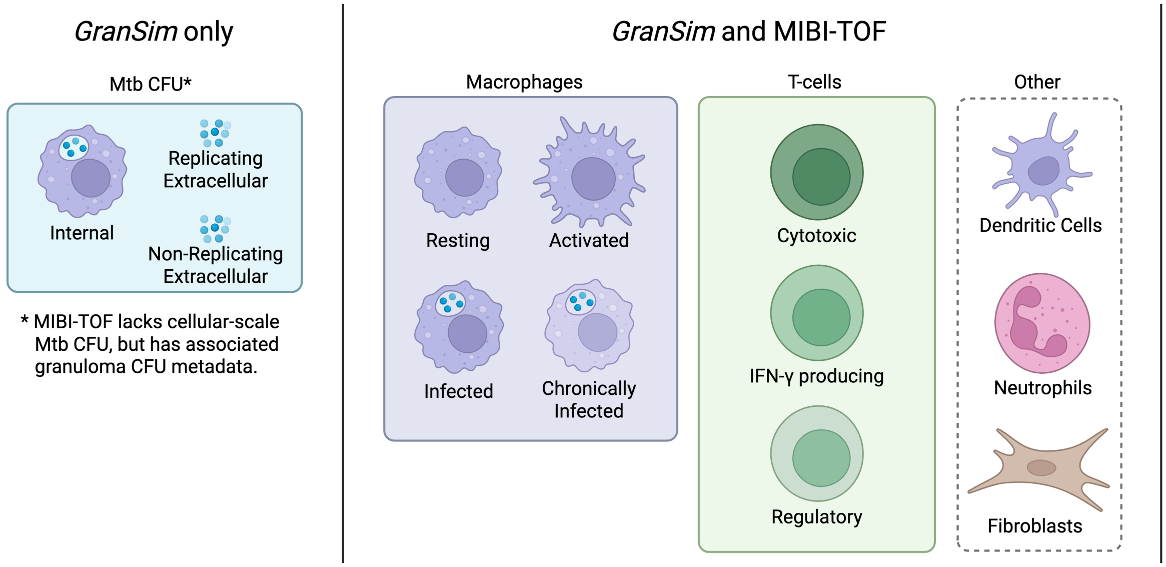


Suppl. Fig. 1 **Representation of cell types between *GranSim* model and MIBI-TOF panel**. The *GranSim* project-specific agents of dendritic cells, neutrophils, and fibroblasts were not simulated in this work. Only total CFU was compared between *GranSim* and CFU. Created in BioRender. Nanda, P. (2026) https://BioRender.com/59yjdap.

### *GranSim* granuloma types

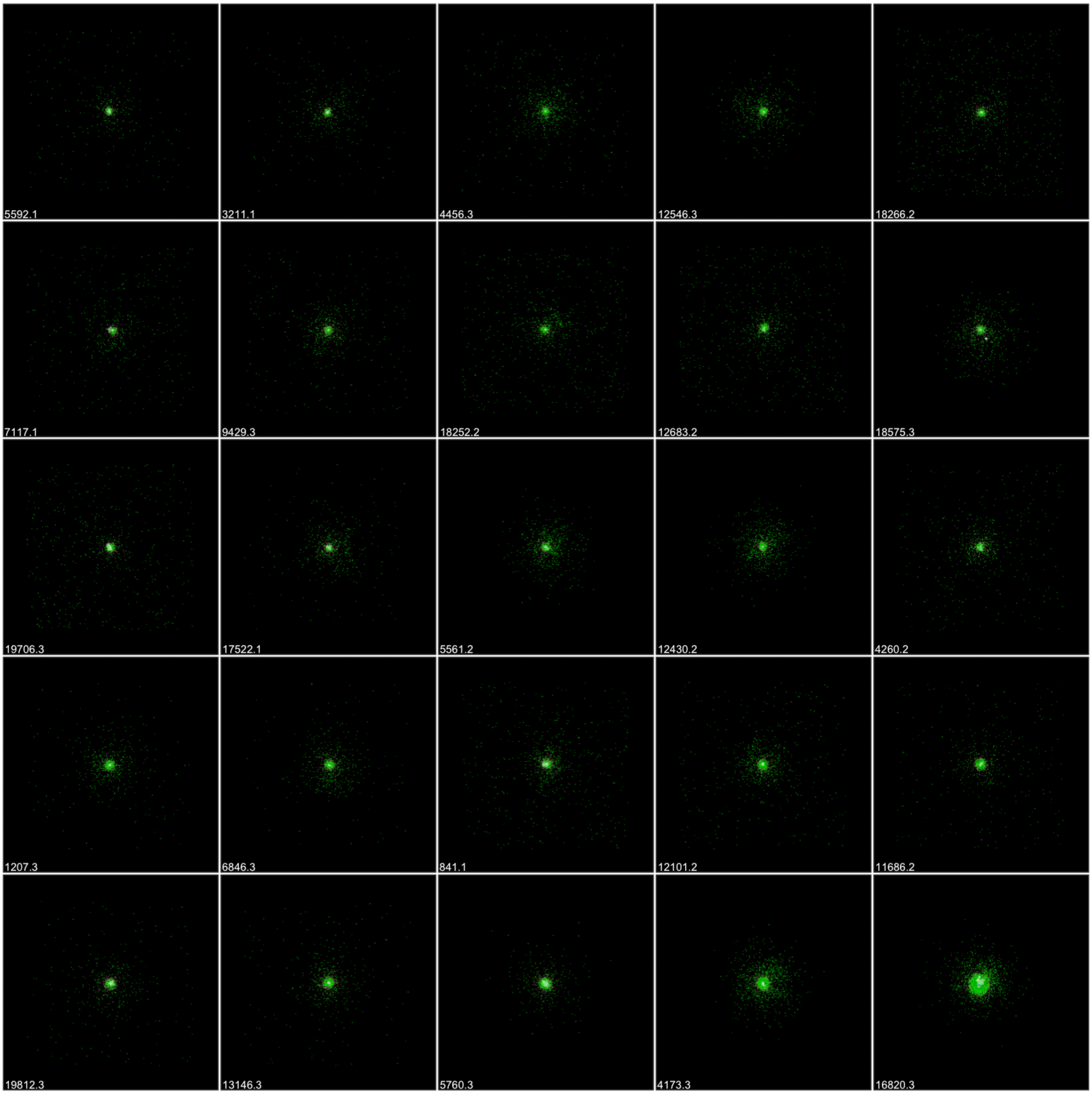


Suppl. Fig. 2 **25 virtual granuloma samples stratified by percentile of CFU in sterilizing *GranSim* granulomas at 22 weeks post-infection**. All images represent 6mm × 6mm of lung tissue. Numbers at bottom-left are *GranSim* sample identifiers (number before the decimal point is the parameter set among 1–20,000; number after the decimal point is the replicate among 1–3). Samples chosen for Fig. 2 are sterile = 16820.3, controlling = 6792.2, uncontrolled = 13222.2. Sterilizing samples that cleared on the first timestep are omitted; simulations are initialized with a single infected macrophage, which means there are instances of sterilizing at the first timestep when macrophages successfully kill the single Mtb bacillus. Such sterilizing events captured by our simulations are below the detection threshold of tissue-level data.

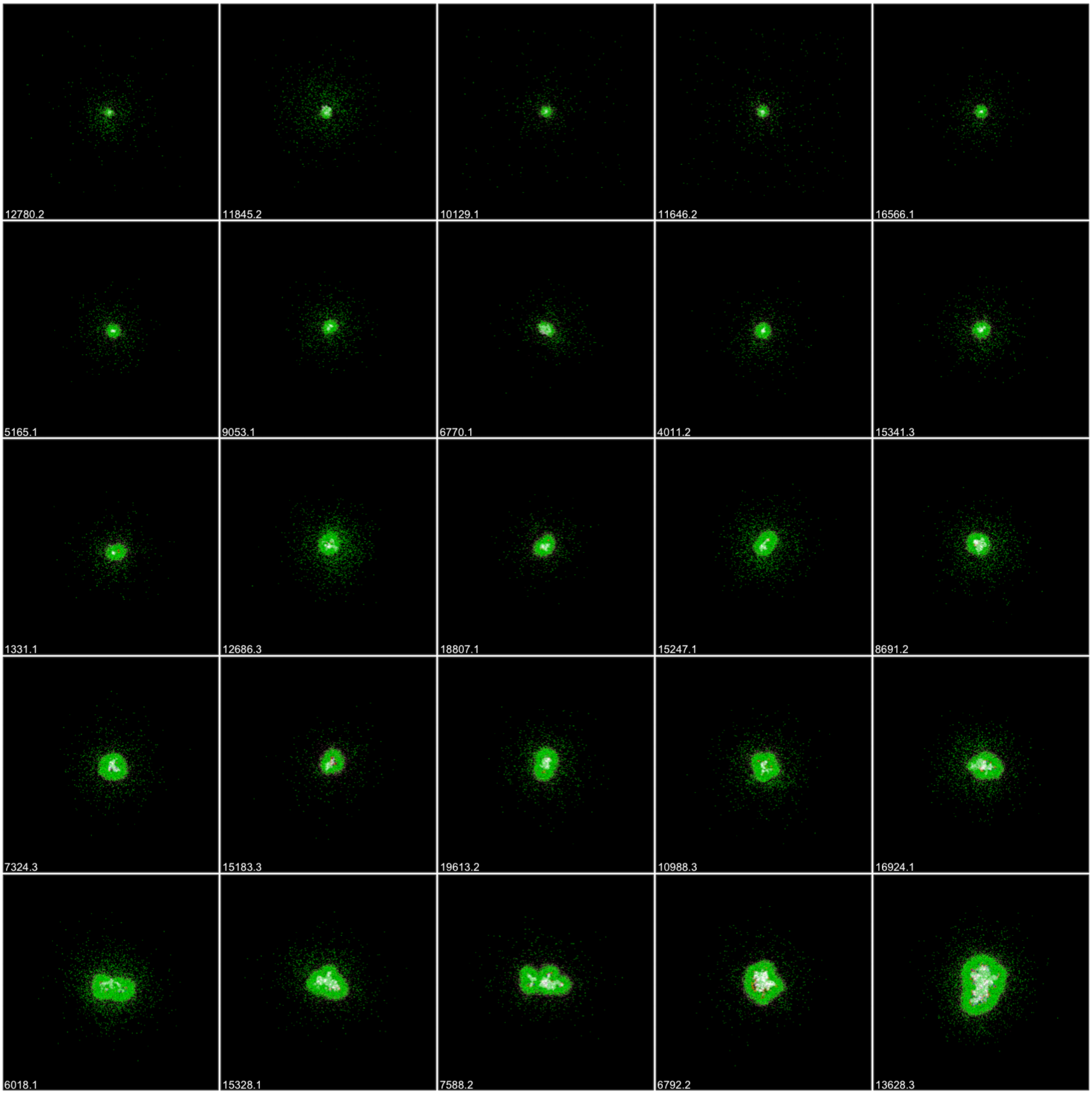


Suppl. Fig. 3: **As above for controlling granulomas.**

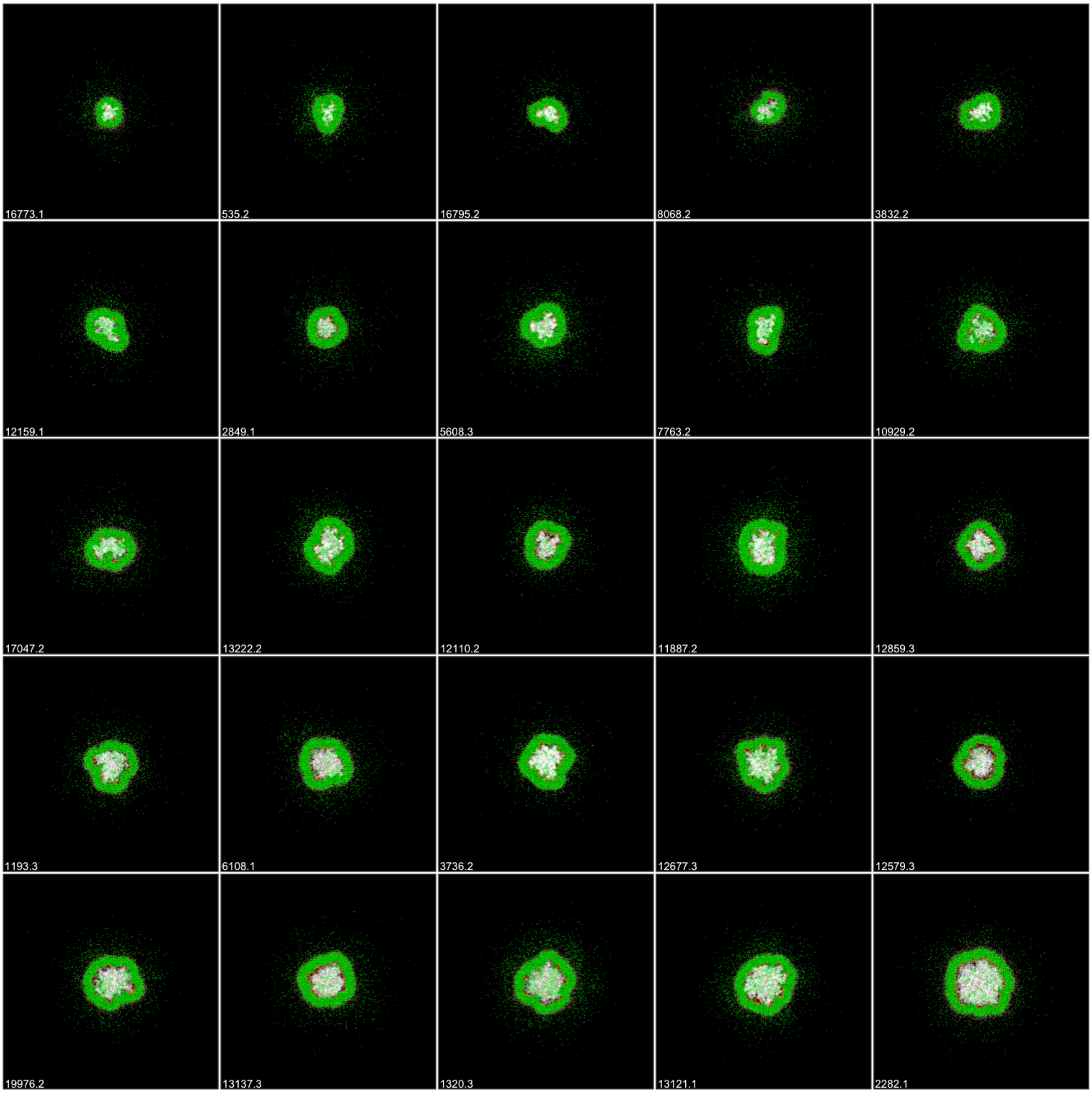


Suppl. Fig. 4 **As above for uncontrolled granulomas.**

### *GranSim* cross-sectional feature list

Suppl. Table 2: **CellProfiler names for top tissue and cellular scale features predictive of log-CFU in simulated granulomas**. Two sets of informative features emerged from analysis: those that appeared 4 or more times contiguously (shown in Fig. 3), and those shown in Extended Data 4. We highlight features appearing in both analyses by shading the table cells. * We chose 60 μm as the neighborhood boundary to be comparable to the 50 μm boundary, which we chose for MIBI-TOF neighborhood summary statistics. † Note that the informative summary statistic of this feature is the median, whereas all others are means. ‡ This is the Euler characteristic of the T cell aggregate, predominantly measuring (# T cells with no T-cell neighbors) - (# of holes in the lymphocytic cuff).

| CellProfiler module and Output statistic | Descriptive name | Scale |
|---|---|---|
| **(i) Features that appear 4 or more times contiguously (see Fig. 3; total of 15 features)** | | |
| MeasureObjectIntensityDistribution macs_RadialDistribution_FracAtD_ifng_3of4_mean | Fraction of IFN-γ near granuloma center | Tissue |
| * MeasureObjectNeighbors mac_Neighbors_SecondClosestDistance_3_median | Distance from macrophage to second-closest cell† | Cellular |
| MeasureObjectIntensity t_cyts_Intensity_UpperQuartileIntensity_tnf_mean | Cytotoxic T cells co-localized with high TNFα concentrations | Tissue |
| MeasureObjectSizeShape ts_AreaShape_EulerNumber_mean | Consolidation of T cells‡ | Tissue |
| MeasureObjectSizeShape macs_AreaShape_Perimeter_mean | Macrophage aggregate perimeter | Tissue |
| MeasureObjectIntensity mac_Intensity_IntegratedIntensityEdge_tnf_mean | Mean macrophage-colocalized TNFα | Cellular |
| * MeasureObjectNeighbors mac_Neighbors_SecondClosestDistance_60_mean | Distance (mean) from macrophage to second-closest cell | Cellular |
| * MeasureObjectNeighbors mac_Neighbors_AngleBetweenNeighbors_60_mean | Angle between neighboring macrophages | Cellular |
| MeasureObjectSizeShape macs_AreaShape_MeanRadius_mean | Macrophage aggregate radius | Tissue |
| MeasureObjectSizeShape t_gams_AreaShape_MajorAxisLength_mean | IFN-$\gamma^+$ T cell aggregate major-axis length | Tissue |
| MeasureObjectSizeShape t_gams_AreaShape_MinorAxisLength_mean | IFN-$\gamma^+$ T cell aggregate minor-axis length | Tissue |
| MeasureObjectSizeShape t_cyts_AreaShape_EquivalentDiameter_mean | Cytotoxic T cell aggregate diameter | Tissue |
| MeasureObjectSizeShape t_regs_AreaShape_EquivalentDiameter_mean | Regulatory T cell aggregate diameter | Tissue |
| SplitOrMergeObjects mac_Children_macs_Count_mean | 1 / (Number of macrophages) | Tissue |
| MeasureObjectIntensityDistribution macs_RadialDistribution_FracAtD_ifng_4of4_mean | Fraction of IFN-γ in outer granuloma | Tissue |
| **(ii) Features identified by LASSO and Bayesian variable selection at 22 weeks (see Extended Data 4; total of 12 features)** | | |
| MeasureObjectSizeShape t_gams_AreaShape_MajorAxisLength_mean | IFN-$\gamma^+$ T cell aggregate major-axis length | Tissue |
| MeasureObjectIntensityDistribution macs_RadialDistribution_FracAtD_ifng_4of4_mean | Fraction of IFN-γ in outer granuloma | Tissue |
| * MeasureObjectNeighbors mac_Neighbors_AngleBetweenNeighbors_60_mean | Angle between neighboring macrophages | Cellular |
| MeasureObjectSizeShape t_gams_AreaShape_MinorAxisLength_mean | IFN-$\gamma^+$ T cell aggregate minor-axis length | Tissue |
| SplitOrMergeObjects mac_Children_macs_Count_mean | 1 / (Number of macrophages) | Tissue |
| MeasureObjectSizeShape t_regs_AreaShape_EquivalentDiameter_mean | Regulatory T cell aggregate diameter | Tissue |
| MeasureObjectSizeShape macs_AreaShape_MeanRadius_mean | Macrophage aggregate radius | Tissue |
| MeasureObjectSizeShape t_cyts_AreaShape_EquivalentDiameter_mean | Cytotoxic T cell aggregate diameter | Tissue |
| SplitOrMergeObjects t_gam_Children_t_gams_Count_mean | 1 / (Number of IFN-$\gamma^+$ T cells) | Tissue |
| MeasureObjectIntensityDistribution macs_RadialDistribution_FracAtD_ifng_1of4_mean | Fraction of IFN-γ in granuloma center | Tissue |
| * MeasureObjectNeighbors t_cyt_Neighbors_AngleBetweenNeighbors_60_median | Angle between neighboring cytotoxic T cells† | Cellular |
| MeasureObjectSizeShape ts_AreaShape_FormFactor_mean | Circularity of aggregate T cells | Tissue |

***GranSim* cross-sectional feature selection**

Among the top 20 features of each week, we used unsupervised clustering to help identify features that appeared at multiple times. It is possible for different weeks to have different sets of top 20 features; in our case there were 64 features in total. We narrow down the top features by those that appear multiple weeks, assuming a spatial feature to granuloma state and highly-correlated to CFU would appear at multiple weeks. The number of weeks used to filter these results is a heuristic and the figures below explore thresholds that start as the most stringent (14 features using a threshold of 7 weeks) to the least stringent (all 64 features using no threshold) (Suppl. Fig. 5–Suppl. Fig. 7). We choose a threshold of 3 or more weeks based on the stability of the clusters.

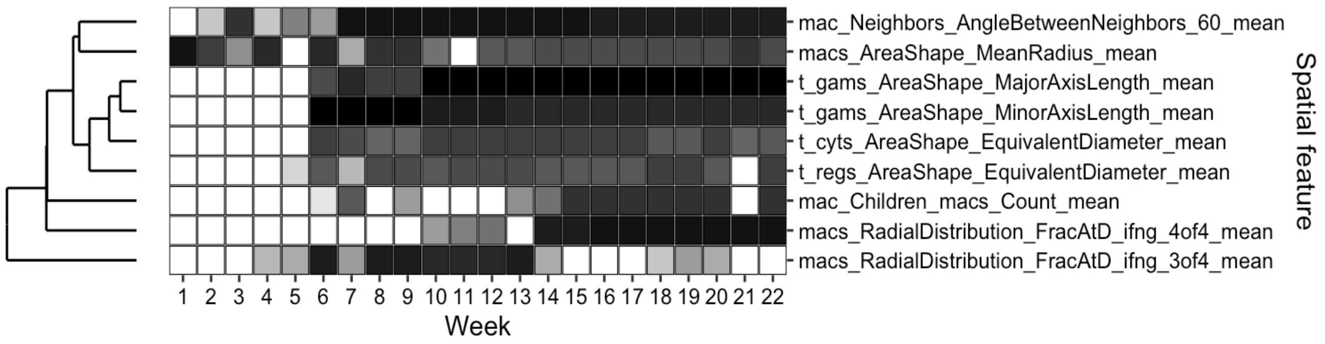


Suppl. Fig. 5 **Exploratory clustering only using features that appear 7 or more times contiguously**. To reveal prominent spatial features driving log-CFU, even after filtering to the top 20 features for each timepoint, there were 64 features in total with some features appearing only once. Therefore, here we show the effect of removing features appearing less frequently to identify drivers across multiple timepoints.

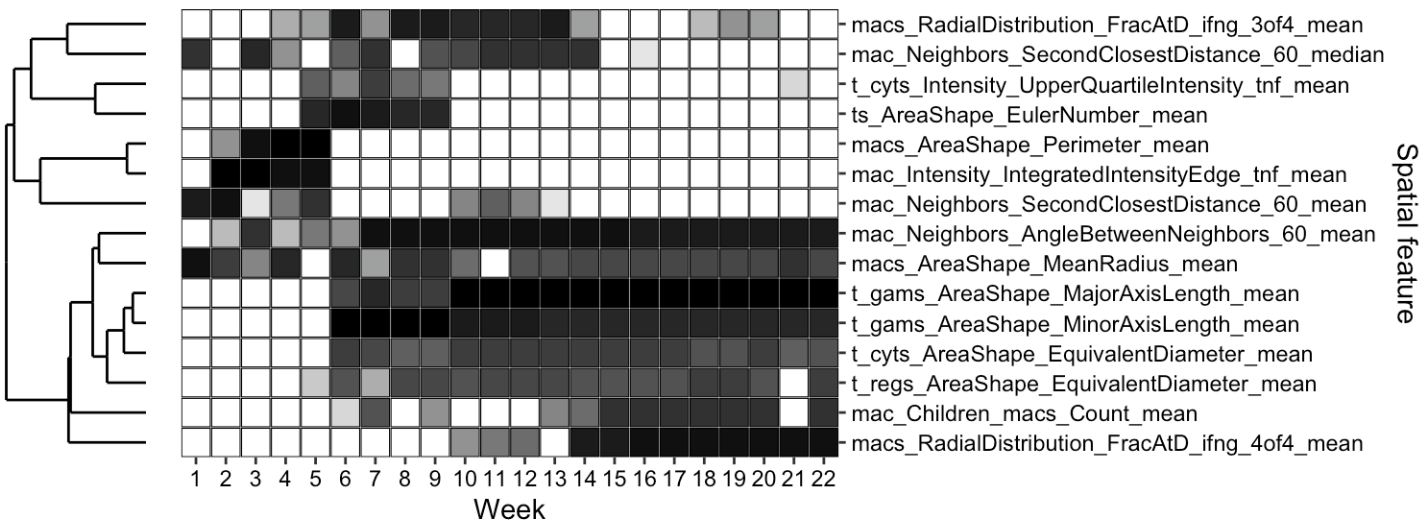


Suppl. Fig. 6: **Exploratory clustering only using features that appear 4 or more times contiguously.** This threshold was used for Fig. 3.

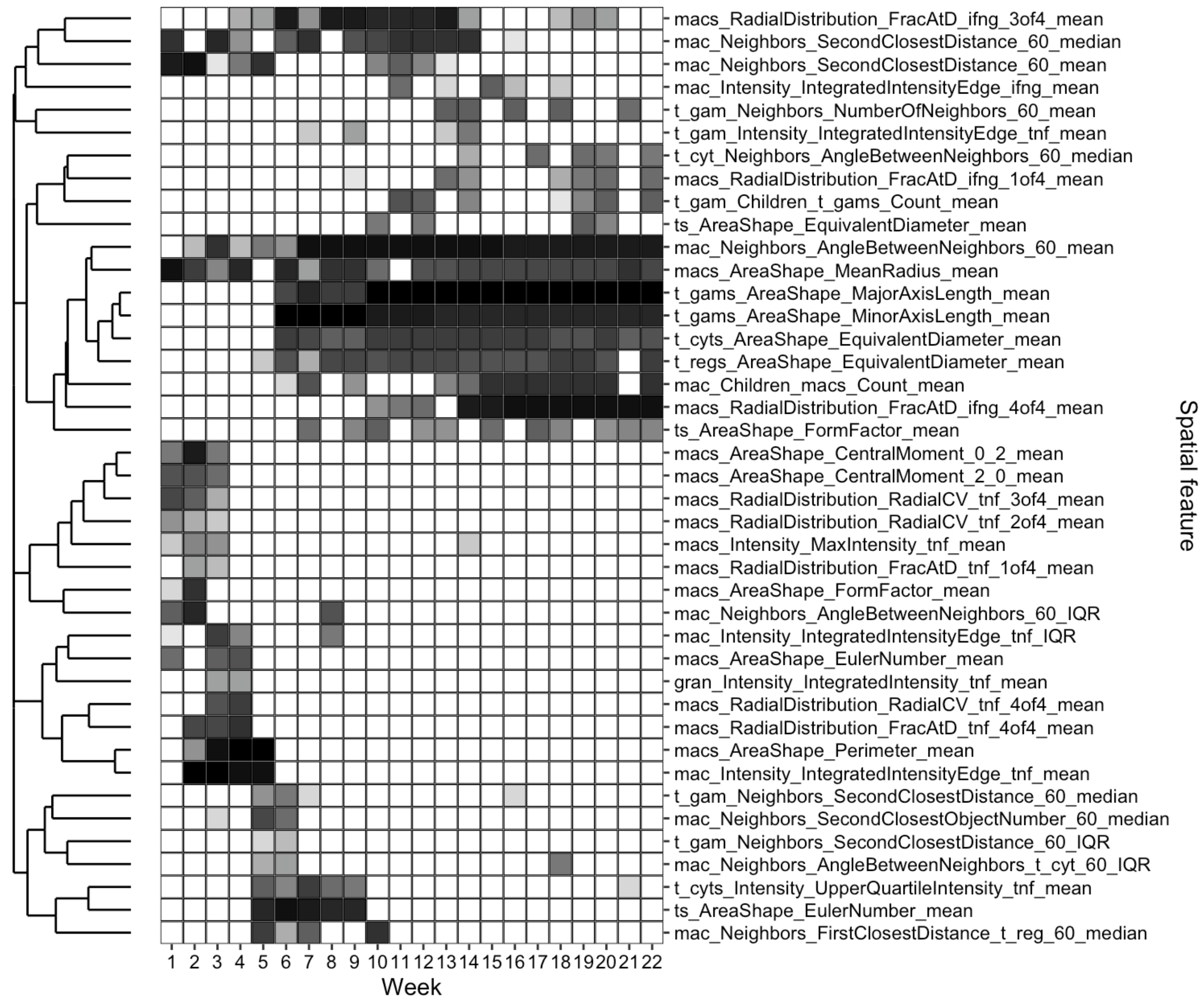


Suppl. Fig. 7: **Exploratory clustering only using features that appear 2 or more times contiguously**

To break ties among the LASSO-identified features, more computationally intensive Bayesian model selection was performed to order the most predictive variables for log-CFU output. At week 22, 12 variables were found sufficient to predict log-CFU output of *GranSim* simulations (Suppl. Fig. 8). The agreement among variables using 5-fold cross-validation is shown in Suppl. Fig. 9.

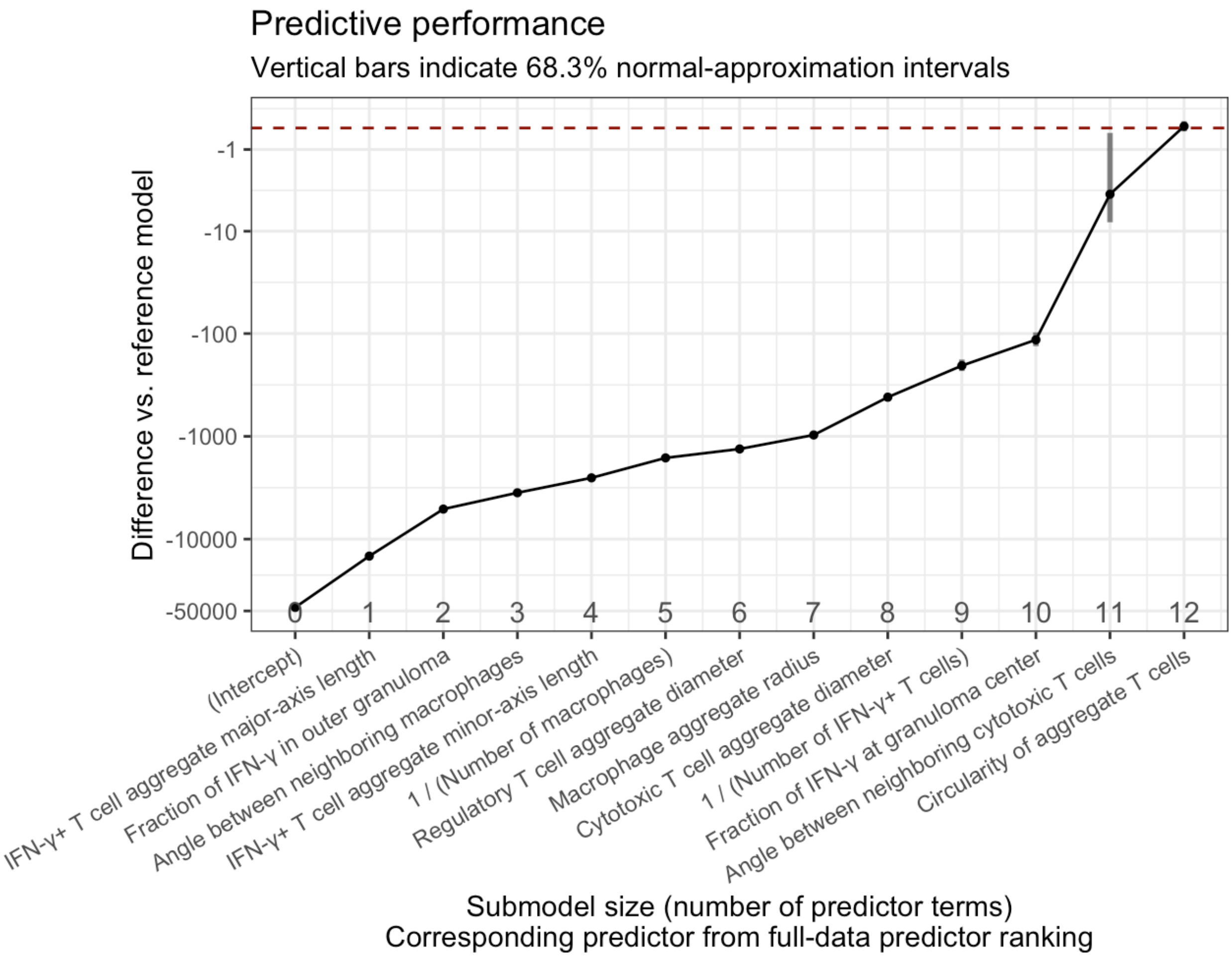


Suppl. Fig. 8: **Bayesian regression performance plot showing LASSO-identified variables ordered from most to least predictive**. We provide mappings of CellProfiler feature names to our descriptive feature names in Suppl. Table 2.

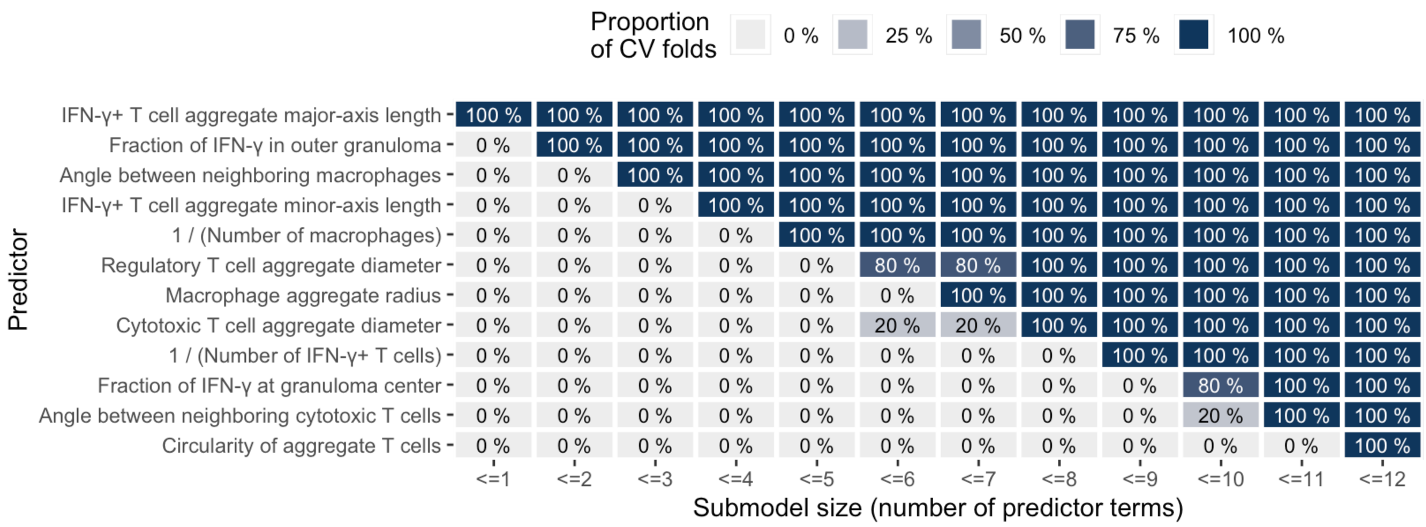


Suppl. Fig. 9 **Bayesian model selection of top LASSO variables showing agreement of 5-fold cross-validation.**

***GranSim* correlation of selected features with CFU outcome**

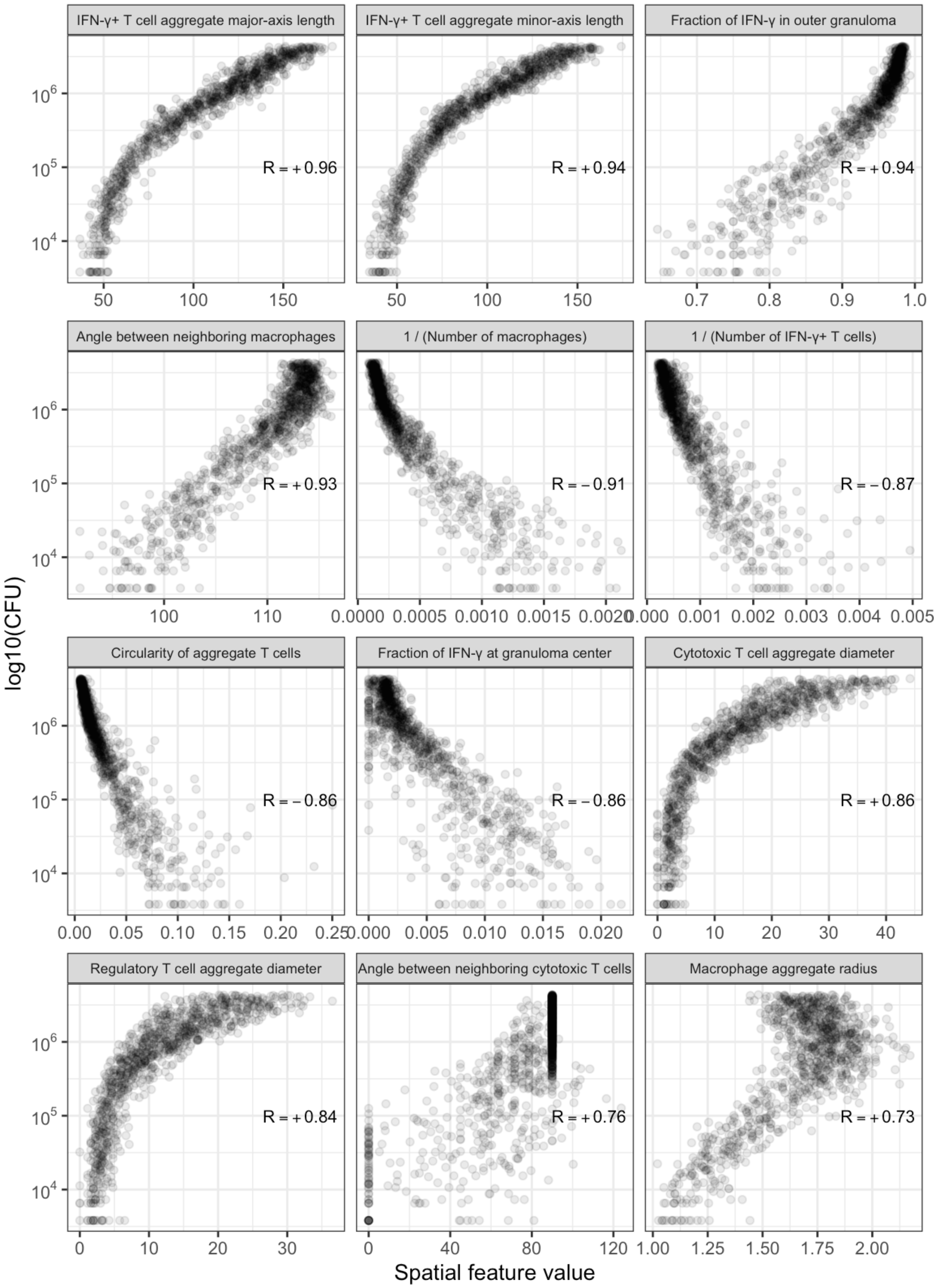


Suppl. Fig. 10 **Example correlations between spatial features and log-CFU.** We show scatter plots of spatial features against log-CFU with Pearson's correlation coefficient (R). Descriptions of the covariates are detailed in the same order as the plots moving left-to-right and top-to-bottom. Note that the R values are only provided as visual aids; the LASSO variable selection centers and scales the covariates to fit the linear model and does not use R. LASSO correlations like these are used to determine the most predictive spatial features, such as those presented in Fig. 3**.**

### Digital partner cost matrix and sample similarity

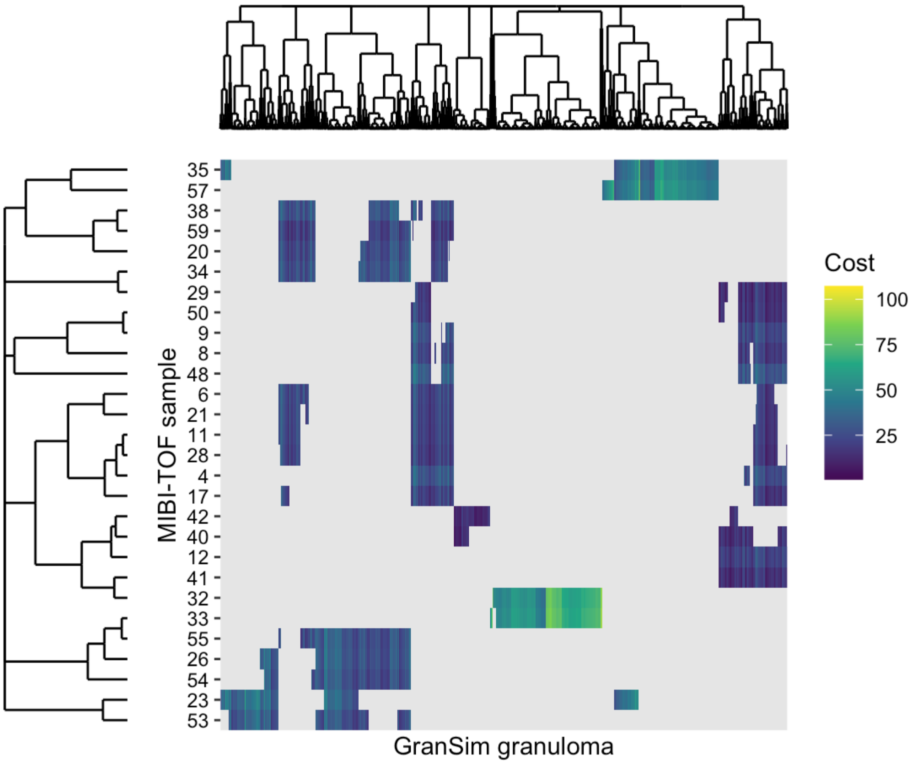


Suppl. Fig. 11 **Digital partner cost matrix visualization**. Hierarchical clustered cost matrix of feature distances between MIBI-TOF samples (rows) and simulated granulomas (columns). Gray areas are simulated granulomas omitted from being too far away from MIBI-TOF log-CFU (see methods). Samples 40 and 41 are most dissimilar to the simulations.

### Digital partner images of tissue samples

In Fig. 5, we showed the top three digital partners for granulomas 4, 33, and 57. Here, we show the top three digital partners for all other granulomas in the experimental library. We show controlling granulomas in Suppl. Fig. 12–Suppl. Fig. 18, the second uncontrolled-growth granuloma in Suppl. Fig. 19, and the indeterminate granulomas in Suppl. Fig. 20–Suppl. Fig. 21.

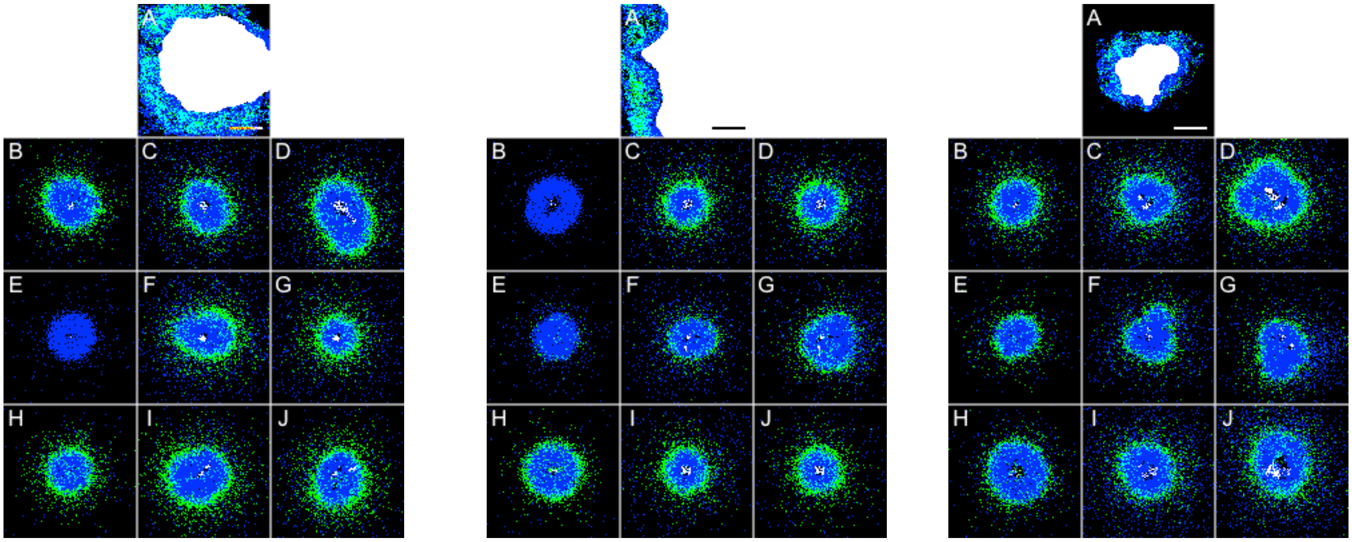

Suppl. Fig. 12 **Granulomas 6, 8, and 9 with their digital partners.** For each granuloma, all three digital partners predict controlled, non-sterile infection. Figure coloration, organization, and scale are described in Fig. 5.

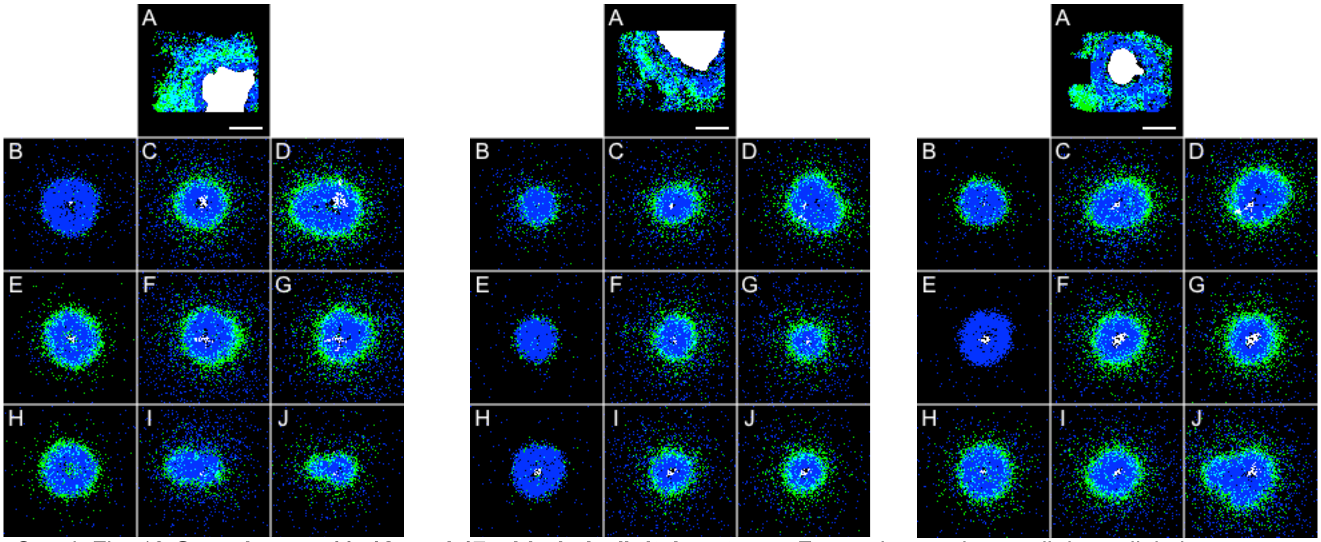

Suppl. Fig. 13 **Granulomas 11, 12, and 17 with their digital partners.** For each granuloma, all three digital partners predict controlled, non-sterile infection. Figure coloration, organization, and scale are described in Fig. 5.

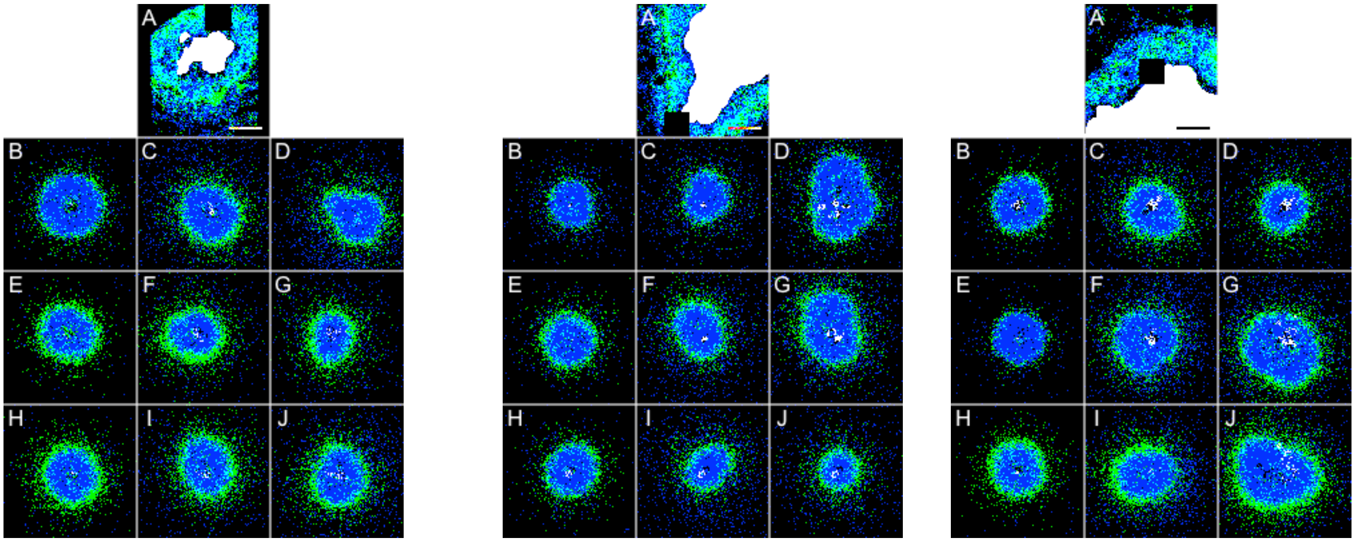


Suppl. Fig. 14 **Granulomas 20, 21, and 26 with their digital partners.** For each granuloma, all three digital partners predict controlled, non-sterile infection. Figure coloration, organization, and scale are described in Fig. 5.

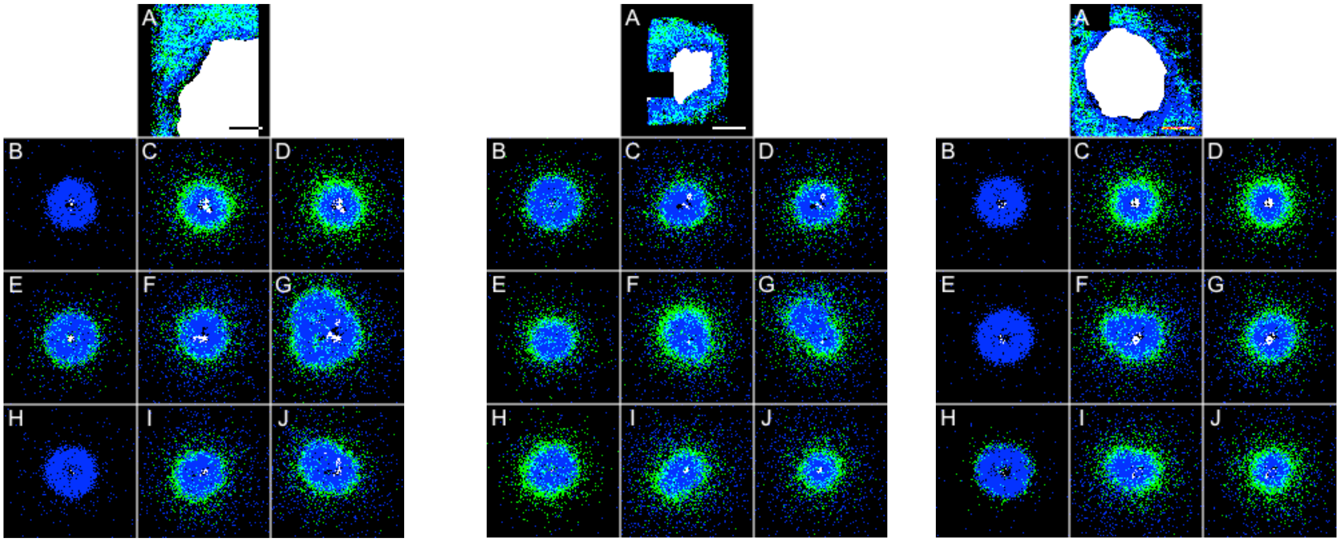


Suppl. Fig. 15 **Granulomas 28, 29, and 38 with their digital partners.** For each granuloma, all three digital partners predict controlled, non-sterile infection. Figure coloration, organization, and scale are described in Fig. 5.

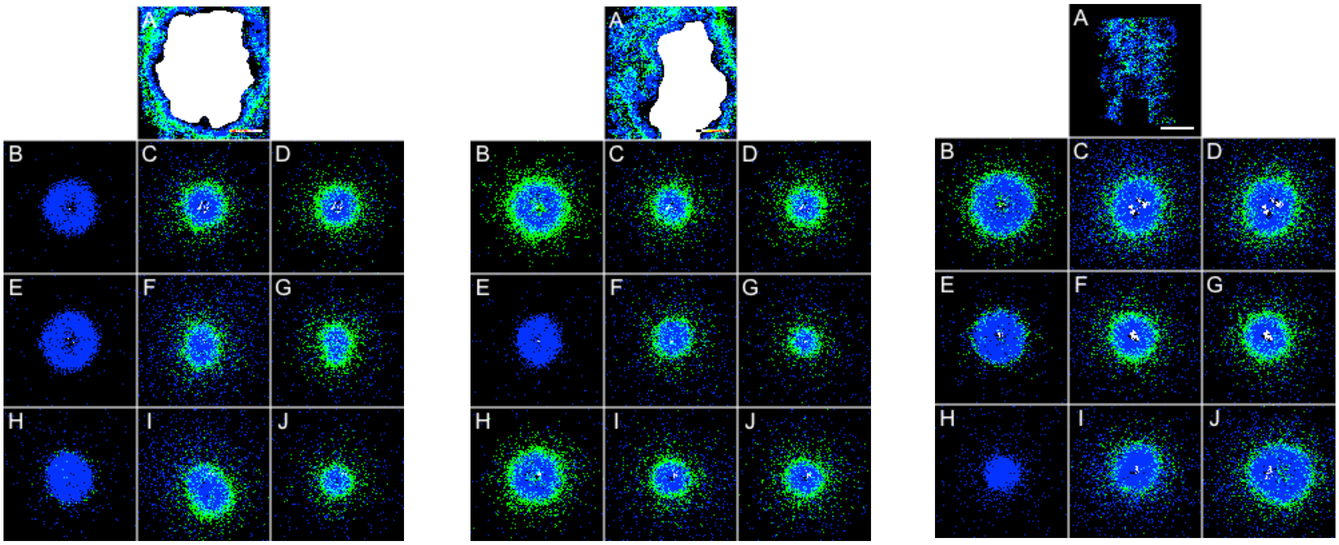


Suppl. Fig. 16 **Granulomas 41, 42, and 48 with their digital partners.** For each granuloma, all three digital partners predict controlled, non-sterile infection. Figure coloration, organization, and scale are described in Fig. 5.

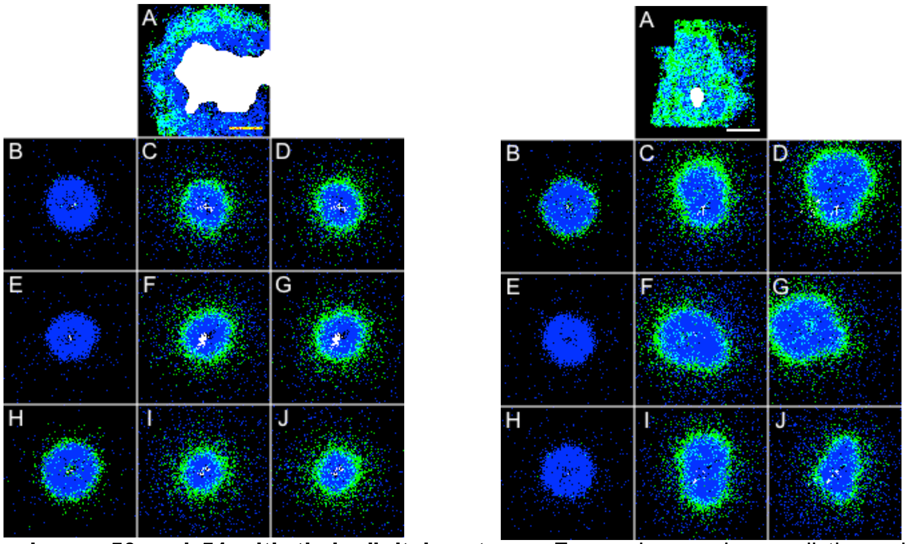


Suppl. Fig. 17 **Granulomas 50 and 54 with their digital partners.** For each granuloma, all three digital partners predict controlled, non-sterile infection. Figure coloration, organization, and scale are described in Fig. 5.

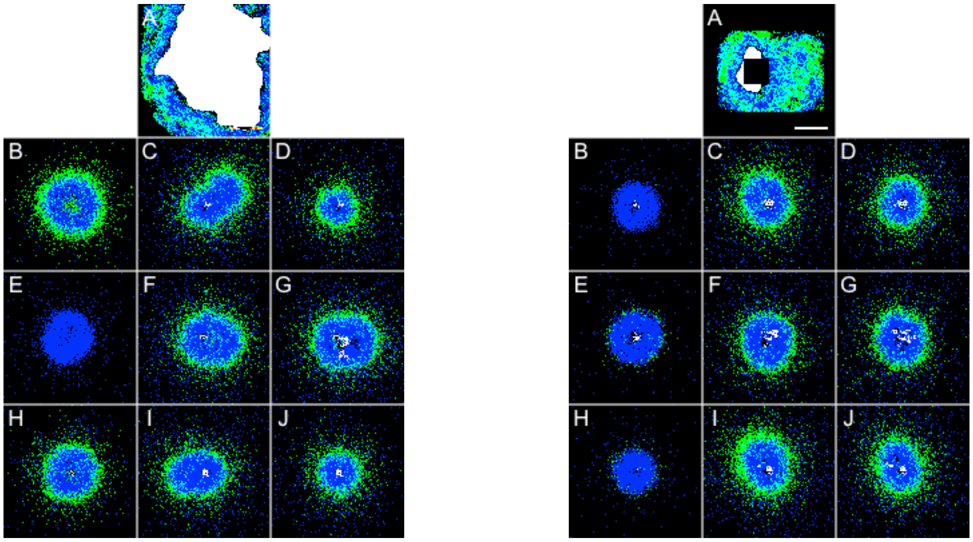


Suppl. Fig. 18 **Granulomas 55 and 59 with their digital partners.** For each granuloma, all three digital partners predict controlled, non-sterile infection. Figure coloration, organization, and scale are described in Fig. 5.

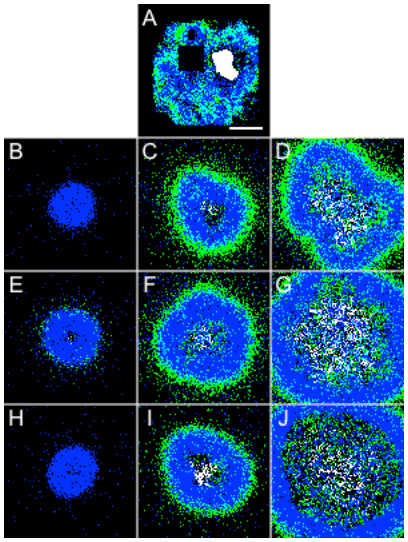


Suppl. Fig. 19 **Granuloma 32 with its digital partners.** All three digital partners of this granuloma predict uncontrolled growth. Figure coloration, organization, and scale are described in Fig. 5.

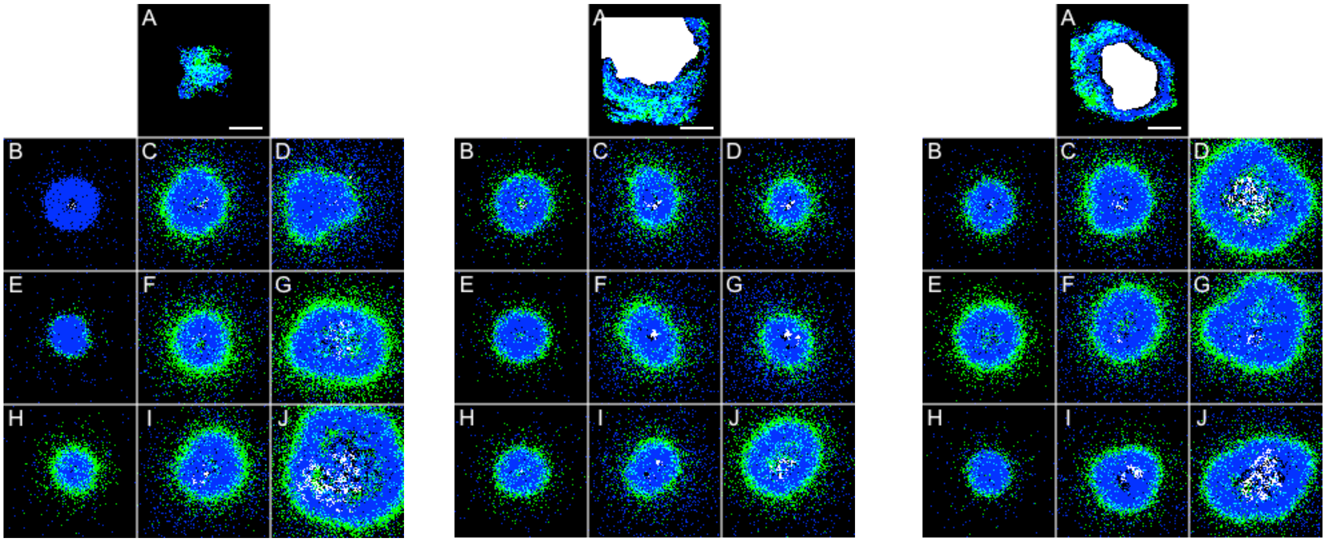

Suppl. Fig. 20 **Granulomas 23, 34, and 35 with their digital partners.** For each granuloma, the top three digital partners are shown as in Fig. 5. The digital partners of these granulomas did not yield consensus prediction and return indeterminate predictions. Figure coloration, organization, and scale are described in Fig. 5.

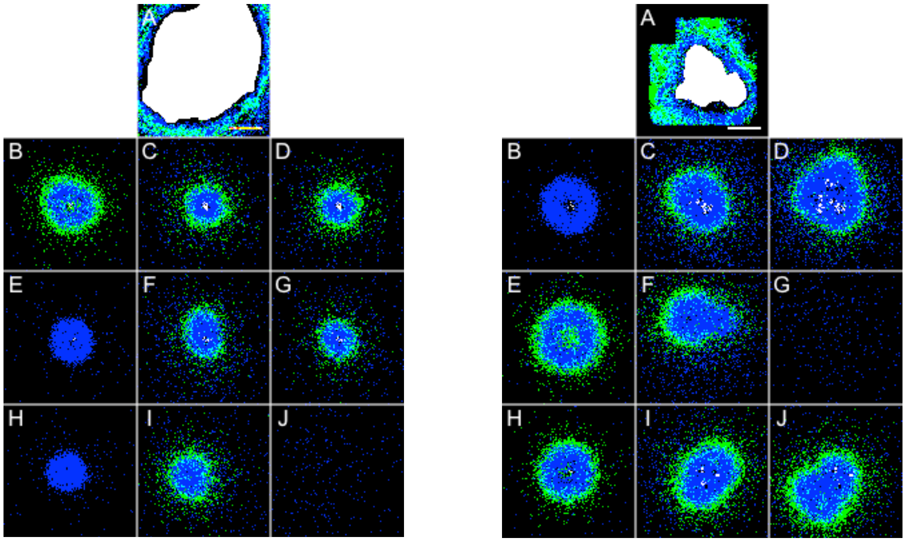

Suppl. Fig. 21 **Granulomas 40 and 53 with their digital partners.** For each granuloma, the top three digital partners are shown as in Fig. 5. The digital partners of these granulomas did not yield consensus prediction and return indeterminate predictions. Figure coloration, organization, and scale are described in Fig. 5.

#### Low correlation of parameters to spatial features

Directly comparing spatial features of non-sterilizing granulomas with model parameters has poor correlation (Suppl. Fig. 22). Therefore, one must instead use partial ranked correlation to discern model parameters driving model outputs. Additionally, this correlation was done against all non-sterilizing granulomas, not the n = 1,100 subset of MIBI-TOF digital partners (Fig. 4).

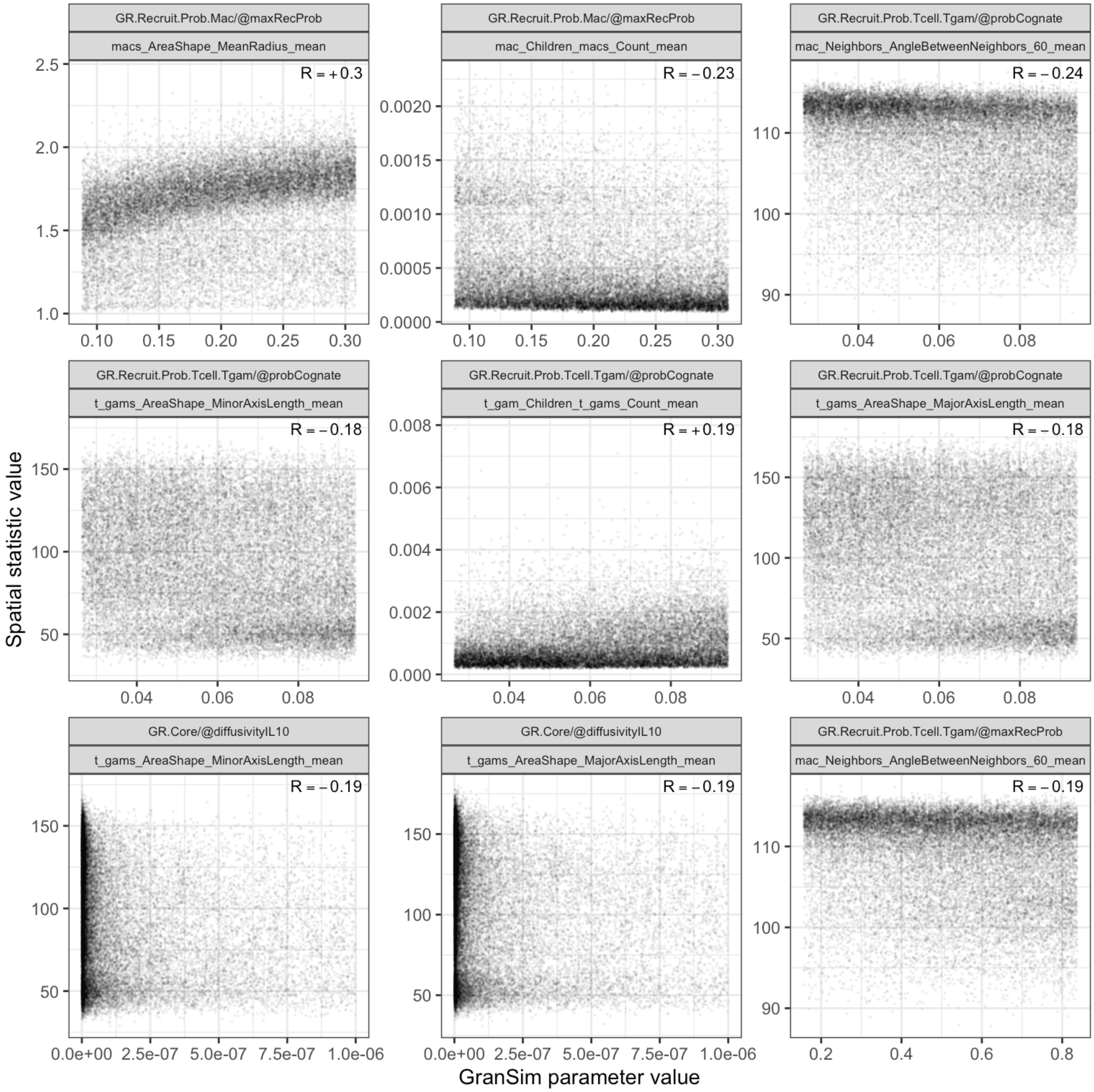


Suppl. Fig. 22 **Top relational scatter plots of spatial statistics against the PRCC identified model parameters using PRCC**.

### Supplementary Material References


1. Davis JM, Ramakrishnan L. The role of the granuloma in expansion and dissemination of early tuberculous infection. Cell. 2009 Jan 9;136(1):37–49. doi:S0092-8674(08)01443-8 [pii] 10.1016/j.cell.2008.11.014 PubMed PMID: 19135887.
2. Lin PL, Rodgers M, Smith L, Bigbee M, Myers A, Bigbee C, et al. Quantitative comparison of active and latent tuberculosis in the cynomolgus macaque model. Infect Immun. 2009 Oct;77(10):4631–42. doi:IAI.00592-09 [pii] 10.1128/IAI.00592-09 PubMed PMID: 19620341.
3. Barry CE, Boshoff HI, Dartois V, Dick T, Ehrt S, Flynn J, et al. The spectrum of latent tuberculosis: rethinking the biology and intervention strategies. Nat Rev Microbiol. 2009 Dec;7(12):845–55. doi:nrmicro2236 [pii] 10.1038/nrmicro2236 PubMed PMID: 19855401.
4. Lin PL, Flynn JL. Understanding latent tuberculosis: a moving target. J Immunol. 2010 Jul 1;185(1):15–22. doi:185/1/15 [pii] 10.4049/jimmunol.0903856 PubMed PMID: 20562268.
5. Kaplan G, Post FA, Moreira AL, Wainwright H, Kreiswirth BN, Tanverdi M, et al. *Mycobacterium tuberculosis* Growth at theCavity Surface: a Microenvironment with FailedImmunity. Infect Immun. 2003 Dec;71(12):7099–108. doi:10.1128/IAI.71.12.7099-7108.2003 PubMed PMID: 14638800; PubMed Central PMCID: PMC308931.
6. Ganchua SKC, White AG, Klein EC, Flynn JL. Lymph nodes-The neglected battlefield in tuberculosis. PLoS Pathog. 2020 Aug;16(8):e1008632. doi:10.1371/journal.ppat.1008632 PubMed PMID: 32790739; PubMed Central PMCID: PMC7425845.
7. Rolo M, González-Blanco B, Reyes CA, Rosillo N, López-Roa P. Epidemiology and factors associated with Extra-pulmonary tuberculosis in a Low-prevalence area. Journal of Clinical Tuberculosis and Other Mycobacterial Diseases. 2023 Aug;32:100377. doi:10.1016/j.jctube.2023.100377 PubMed PMID: 37252369; PubMed Central PMCID: PMC10209530.
8. McCaffrey EF, Donato M, Keren L, Chen Z, Delmastro A, Fitzpatrick MB, et al. The immunoregulatory landscape of human tuberculosis granulomas. Nat Immunol. 2022 Feb;23(2):318–29. doi:10.1038/s41590-021-01121-x PubMed PMID: 35058616; PubMed Central PMCID: PMC8810384.
9. Millar JA, Butler JR, Evans S, Grant NL, Mattila JT, Linderman JJ, et al. Spatial Organization and Recruitment of Non-Specific T Cells May Limit T Cell-Macrophage Interactions Within Mycobacterium tuberculosis Granulomas. Front Immunol. 2021 Jan 20;11. doi:10.3389/fimmu.2020.613638 PubMed PMID: 33552077; PubMed Central PMCID: PMC7855029.
10. Joslyn LR, Renardy M, Weissman C, Grant NL, Flynn JL, Butler JR, et al. Temporal and Spatial Analyses of TB Granulomas to Predict Long-Term Outcomes. In: Vodovotz Y, An G, editors. Complex Systems and Computational Biology Approaches to Acute Inflammation [Internet]. Cham: Springer International Publishing; 2021 [cited 2026 Sep 15]. p. 273–91. Available from: http://link.springer.com/10.1007/978-3-030-56510-7_15 doi:10.1007/978-3-030-56510-7_15
11. Lyu J, Narum DE, Baldwin SL, Larsen SE, Bai X, Griffith DE, et al. Understanding the development of tuberculous granulomas: insights into host protection and pathogenesis, a review in humans and animals. Front Immunol. 2024 Dec 9;15:1427559. doi:10.3389/fimmu.2024.1427559 PubMed PMID: 39717773; PubMed Central PMCID: PMC11663721.
12. Cicchese JM, Evans S, Hult C, Joslyn LR, Wessler T, Millar JA, et al. Dynamic balance of pro- and anti-inflammatory signals controls disease and limits pathology. Immunological Reviews. 2018;285(1):147–67. doi:10.1111/imr.12671 PubMed PMID: 30129209; PubMed Central PMCID: PMC6292442.
13. Lin PL, Ford CB, Coleman MT, Myers AJ, Gawande R, Ioerger T, et al. Sterilization of granulomas is common in active and latent tuberculosis despite within-host variability in bacterial killing. Nat Med. 2014 Jan;20(1):75–9. doi:10.1038/nm.3412 PubMed PMID: 24336248.
14. Co DO, Hogan LH, Karman J, Heninger E, Vang S, Wells K, et al. Interactions between T Cells Responding to Concurrent Mycobacterial and Influenza Infections. The Journal of Immunology. 2006 Dec 15;177(12):8456–65. doi:10.4049/jimmunol.177.12.8456 PubMed PMID: 17142743.
15. Marakalala MJ, Raju RM, Sharma K, Zhang YJ, Eugenin EA, Prideaux B, et al. Inflammatory signaling in human tuberculosis granulomas is spatially organized. Nat Med. 2016 May;22(5):531–8. doi:10.1038/nm.4073 PubMed PMID: 27043495; PubMed Central PMCID: PMC4860068.
16. Liu H, Li P, He S, Yang H, Yuan T, He X, et al. The role and mechanisms of multiple immunoregulatory cells in pulmonary tuberculosis. Front Immunol. 2026 Jun 2;17:1841581. doi:10.3389/fimmu.2026.1841581
17. Capuano SV, Croix DA, Pawar S, Zinovik A, Myers A, Lin PL, et al. Experimental Mycobacterium tuberculosis infection of cynomolgus macaques closely resembles the various manifestations of human M. tuberculosis infection. Infect Immun. 2003 Oct;71(10):5831–44.
18. Lin PL, Pawar S, Myers A, Pegu A, Fuhrman C, Reinhart TA, et al. Early events in Mycobacterium tuberculosis infection in cynomolgus macaques. Infect Immun. 2006 Jul;74(7):3790–803. doi:74/7/3790 [pii] 10.1128/IAI.00064-06 PubMed PMID: 16790751.

19. Flynn JL, Gideon HP, Mattila JT, Lin PL. Immunology studies in non-human primate models of tuberculosis. Immunol Rev. 2015 Mar;264(1):60–73. doi:10.1111/imr.12258 PubMed PMID: 25703552.
20. Gideon HP, Phuah J, Myers AJ, Bryson BD, Rodgers MA, Coleman MT, et al. Variability in tuberculosis granuloma T cell responses exists, but a balance of pro- and anti-inflammatory cytokines is associated with sterilization. PLoS Pathog. 2015 Jan;11(1):e1004603. doi:10.1371/journal.ppat.1004603 PubMed PMID: 25611466.
21. Martin CJ, Cadena AM, Leung VW, Lin PL, Maiello P, Hicks N, et al. Digitally Barcoding Mycobacterium tuberculosis Reveals In Vivo Infection Dynamics in the Macaque Model of Tuberculosis. MBio. 2017 May 9;8(3). doi:10.1128/mBio.00312-17 PubMed PMID: 28487426.
22. Cadena AM, Fortune SM, Flynn JL. Heterogeneity in tuberculosis. Nature reviews Immunology. 2017 Nov;17(11):691–702. doi:10.1038/nri.2017.69 PubMed PMID: 28736436.
23. Young DB, Gideon HP, Wilkinson RJ. Eliminating latent tuberculosis. Trends Microbiol. 2009 May;17(5):183–8. doi:10.1016/j.tim.2009.02.005 PubMed PMID: 19375916.
24. Gideon HP, Hughes TK, Tzouanas CN, Wadsworth MH, Tu AA, Gierahn TM, et al. Multimodal profiling of lung granulomas in macaques reveals cellular correlates of tuberculosis control. Immunity. 2022 May;55(5):827-846.e10. doi:10.1016/j.immuni.2022.04.004 PubMed PMID: 35483355; PubMed Central PMCID: PMC9122264.
25. McCaffrey EF, Delmastro AC, Fitzhugh I, Ranek JS, Douglas S, Peters JM, et al. The immunometabolic topography of cellular organization and bacterial control in tuberculosis granulomas. Nat Immunol. 2026 Apr;27(4):867–80. doi:10.1038/s41590-026-02431-8 PubMed PMID: 41731147; PubMed Central PMCID: PMC13361869.
26. Shen X, Wu T, Ji X, Yang K, Wang L, Peng Y, et al. Mycobacterium tuberculosis infection depressed cytotoxic T cells activity owing to decreasing NKG2C and increasing NKG2A expression. Molecular Immunology. 2023 Oct;162:133–42. doi:10.1016/j.molimm.2023.08.014 PubMed PMID: 37683324.
27. Algood HM, Lin PL, Flynn JL. Tumor necrosis factor and chemokine interactions in the formation and maintenance of granulomas in tuberculosis. Clin Infect Dis. 2005 Aug 1;41 Suppl 3:S189-93. doi:CID35769 [pii] 10.1086/429994 PubMed PMID: 15983898.
28. Khader SA, Bell GK, Pearl JE, Fountain JJ, Rangel-Moreno J, Cilley GE, et al. IL-23 and IL-17 in the establishment of protective pulmonary CD4+ T cell responses after vaccination and during Mycobacterium tuberculosis challenge. Nat Immunol. 2007 Apr;8(4):369–77.
29. Lin PL, Plessner HL, Voitenok NN, Flynn JL. Tumor necrosis factor and tuberculosis. J Investig Dermatol Symp Proc. 2007 May;12(1):22–5.
30. Millington KA, Innes JA, Hackforth S, Hinks TSC, Deeks JJ, Dosanjh DPS, et al. Dynamic Relationship between IFN-γ and IL-2 Profile of *Mycobacterium tuberculosis* -Specific T Cells and Antigen Load. The Journal of Immunology. 2007 Apr 15;178(8):5217–26. doi:10.4049/jimmunol.178.8.5217 PubMed PMID: 17404305; PubMed Central PMCID: PMC2743164.
31. Green AM, Difazio R, Flynn JL. IFN-gamma from CD4 T cells is essential for host survival and enhances CD8 T cell function during Mycobacterium tuberculosis infection. J Immunol. 2013 Jan 1;190(1):270–7. doi:10.4049/jimmunol.1200061 PubMed PMID: 23233724.
32. O'Garra A, Redford PS, McNab FW, Bloom CI, Wilkinson RJ, Berry MP. The immune response in tuberculosis. Annu Rev Immunol. 2013;31:475–527. doi:10.1146/annurev-immunol-032712-095939 PubMed PMID: 23516984.
33. Lyadova IV. Neutrophils in Tuberculosis: Heterogeneity Shapes the Way? Mediators of inflammation. 2017;2017:8619307. doi:10.1155/2017/8619307 PubMed PMID: 28626346.
34. Scriba TJ, Coussens AK, Fletcher HA. Human Immunology of Tuberculosis. Microbiol Spectr. 2017 Jan;5(1). doi:10.1128/microbiolspec.TBTB2-0016-2016 PubMed PMID: 28155806.
35. Esaulova E, Das S, Singh DK, Choreño-Parra JA, Swain A, Arthur L, et al. The immune landscape in tuberculosis reveals populations linked to disease and latency. Cell Host & Microbe. 2021 Feb;29(2):165-178.e8. doi:10.1016/j.chom.2020.11.013 PubMed PMID: 33340449; PubMed Central PMCID: PMC7878437.
36. Lin PL, Coleman T, Carney JP, Lopresti BJ, Tomko J, Fillmore D, et al. Radiologic responses in cynomolgous macaques for assessing tuberculosis chemotherapy regimens. Antimicrob Agents Chemother. 2013 Jun 24. doi:10.1128/AAC.00277-13 PubMed PMID: 23796926.
37. Smith JA. Neutrophils, host defense, and inflammation: a double-edged sword. Journal of Leukocyte Biology. 1994 Dec 1;56(6):672–86. doi:10.1002/jlb.56.6.672 PubMed PMID: 7996043.
38. Ghoshal A, Verma A, Bhaskar A, Dwivedi VP. The uncharted territory of host-pathogen interaction in tuberculosis. Front Immunol. 2024 Jan 19;15:1339467. doi:10.3389/fimmu.2024.1339467 PubMed PMID: 38312835; PubMed Central PMCID: PMC10834760.
39. Gammack D, Doering CR, Kirschner DE. Macrophage response to Mycobacteriumtuberculosis infection. J Math Biol. 2004 Feb 1;48(2):218–42. doi:10.1007/s00285-003-0232-8

40. Hunter L, Hingley-Wilson S, Stewart GR, Sharpe SA, Salguero FJ. Dynamics of Macrophage, T and B Cell Infiltration Within Pulmonary Granulomas Induced by Mycobacterium tuberculosis in Two Non-Human Primate Models of Aerosol Infection. Front Immunol. 2022 Jan 6;12:776913. doi:10.3389/fimmu.2021.776913 PubMed PMID: 35069548; PubMed Central PMCID: PMC8770544.
